# A Hierarchy of Biomolecular Proportional-Integral-Derivative Feedback Controllers for Robust Perfect Adaptation and Dynamic Performance

**DOI:** 10.1101/2021.03.21.436342

**Authors:** Maurice Filo, Sant Kumar, Mustafa Khammash

## Abstract

Proportional-Integral-Derivative (PID) feedback controllers have been the most widely used controllers in industry for almost a century due to their good performance, simplicity, and ease of tuning. Motivated by their success in various engineering disciplines, PID controllers recently found their way into synthetic biology, where the design of feedback molecular control systems has been identified as an important goal. In this paper, we consider the mathematical realization of PID controllers via biomolecular interactions. We propose an array of topologies that offer a compromise between simplicity and high performance. We first demonstrate that different Proportional-Integral (PI) controllers exhibit different capabilities for enhancing the dynamics and reducing variance (cell-to-cell variability). Next, we introduce several derivative controllers that are realized based on incoherent feedforward loops acting in a feedback configuration. Alternatively, we show that differentiators can be realized by placing molecular integrators in a negative feedback loop—an arrangement that can then be augmented by PI components to yield PID feedback controllers. We demonstrate that the derivative component can be exploited for enhancing system stability, dramatically increasing the molecular control system’s dynamic performance, and for reducing the noise effect on the output of interest. The PID controller features are established through various deterministic and stochastic analyses as well as numerical simulations. Finally, we provide an experimental demonstration using a recently developed hybrid setup, the cyberloop, where the controller is implemented *in silico* to control a biological genetic circuit *in vivo*. The large array of novel biomolecular PID controllers introduced here forms a basis for the design and construction of advanced high-performance biomolecular control systems that robustly regulate the dynamics of living systems.

---

**O**ne of the most salient features of biological systems is their ability to adapt to their noisy environments. For example, cells often regulate gene expression to counteract intrinsic and extrinsic noise in order to maintain a desirable behavior in a precise and timely fashion. This resilience toward undesired disturbances is often achieved via feedback control that has proved to be ubiquitous in both natural (e.g. [2–4]) and engineered systems (e.g. [5,6]). In fact, synthetic engineering of biomolecular feedback controllers is gaining a wide attention from biologists and engineers (e.g. [7–15]).

A standard general setup for feedback controllers is depicted as a block diagram in Figure 1(a) (refer to Box 1. A Primer on Block Diagrams in the SI). The “Plant” block represents the process to be controlled. It can be actuated through its input, denoted here by u, to dynamically manipulate its output of interest, denoted here by *y*. The objective of such control systems is to design a feedback controller that *automatically* actuates the plant in a smart autonomous fashion which guarantees that the output *y* meets certain performance goals despite the presence of disturbances in the plant. These performance goals, described in Figure 1(b), include (but are not limited to) Robust Perfect Adaptation (RPA), stability enhancement, desirable transient response and variance reduction. Control theory developed a wide set of tools to design feedback controllers that meet certain performance objectives. For instance, it is well known in control theory (Internal Model Principle [16]) that a controller should include Integral (I) action to be able to achieve RPA. Furthermore, Proportional-Integral-Derivative (PID) feedback controllers—first rigorously introduced by Nicolas Minorsky [17] around a hundred years ago—adds Proportional (P) and Derivative (D) action to the Integrator (I) to be able to tune the transient dynamics and enhance stability while preserving RPA. Interestingly, after almost a century, PID controllers are still the most widely used controllers in industrial applications spanning a broad range of engineering disciplines such as chemical, mechanical, and electrical engineering [18–20].

**Figure 1:**
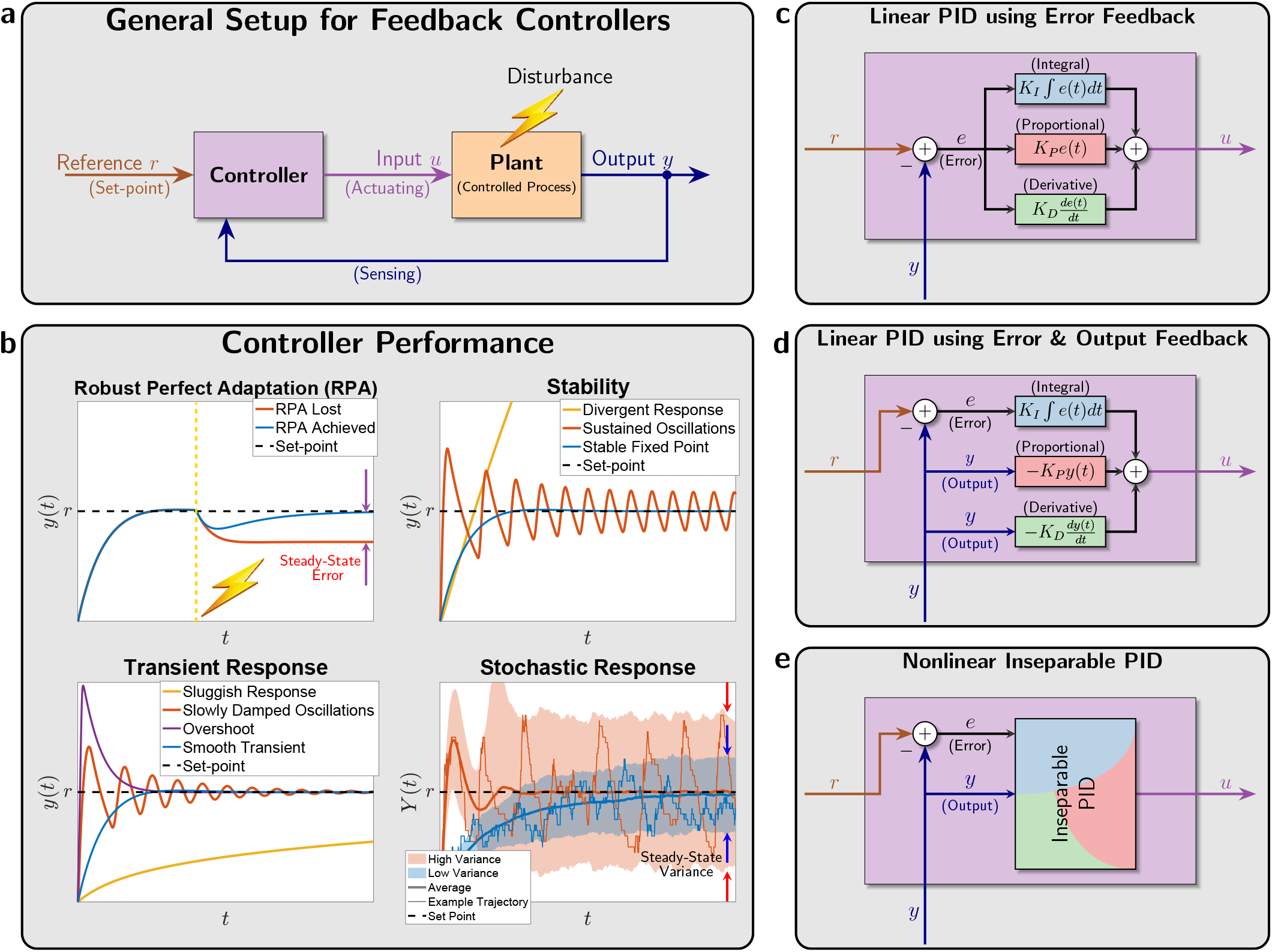
Feedback controller design and performance. **(a)** The output to be controlled is fed back into the controller via a sensing mechanism. The controller exploits the set-point, that is typically “dialed in” by the user and computes the suitable control action to be applied to the plant (or process) via an actuation mechanism. The goal of the control action is to steer the output to the desired set-point despite external or even internal disturbances. **(b)** A demonstration of four performance goals that are typically targeted when designing the controller. **Robust Perfect Adaptation (RPA)**: This is the biological analogue of the notion of *Robust Steady*-*State Tracking* (RSST) that is well known in control theory [1]. A controller achieves RPA if it drives the steady state of the plant output *y* to the set-point (or reference, denoted by *r*) despite varying initial conditions, plant uncertainties and/or constant disturbances. **Stability Enhancement**: A typical goal of a controller is to stabilize the dynamics. That is, it forces the output *y* to converge to a fixed steady-state value thus avoiding divergent responses and sustained oscillations. **Desirable Transient Response**: Another typical control objective is to yield a smooth transient response which is fast enough but doesn’t overshoot or oscillate too much. **Variance Reduction**: For stochastic dynamics, it is common to study the time evolution of the output probability distribution and its moments such as the mean and variance. A natural performance objective is to design a controller that tightens the probability distribution around the mean, e.g. reduce the variance (cell-to-cell variability). **(c), (d), and (e) Various PID control architectures.**The classical designs in (c) and (d) involve separate linear P, I and D operations that are added together to yield the control action *u*. The difference between (c) and (d) is in the controller input: in (c) the error signal is the only input, while in (d) the error signal is fed into the I component whereas the output signal is fed into the P and D components. In this paper, we propose PID control architectures that fit in the more general class depicted in (e) where the PID components may be nonlinear and inseparable. This gives more mathematical realization flexibility for biomolecular controllers.

Originally, PID feedback controllers were designed to control mechanical (later, electrical and chemical) systems such as automatic ship steering [21]. Such control systems involve controlling quantities that can take both negative and positive values such as angles, velocities, electric currents, voltages, etc. Furthermore, traditional PID controllers possess linear dynamics since all three operations of a PID are linear. Two classes of linear PID controllers, adopted from [22, Chapter 10], are shown in Figures 1(c) and (d). In Figure 1(c), the error signal *e*(*t*):= *r* – *y*(*t*) is fed into the three (P, I, and D) components. The outputs of the three components are summed up to yield the control action *u* which serves as the actuation input to the plant. However, in Figure 1(d), the controller has two degrees of freedom since both the error e and the output *y* are used separately and simultaneously. Particularly, the error is fed into the integrator, while the output is fed into the proportional and derivative components. Observe that both architectures require that the integrator operates on the error (and not the output). This is necessary to achieve RPA and can be easily seen using a very simple argument explained next. Let *u_I_*(*t*) denote the output of the integrator, that is

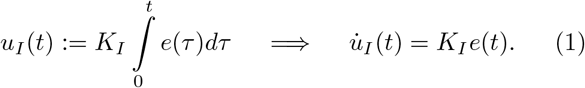

Assuming that the dynamics are stable, then at steady state we have 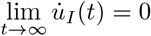. This implies that, at steady state, the error *e*:= *r* – *y* has to be zero, and thus 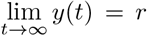, hence achieving the steady-state tracking property. Observe that this argument does not depend on the plant, hence achieving the robustness property.

For mechanical and electrical systems, the linearity of the PID controllers is convenient because of the availability of basic physical parts (e.g. dampers, springs, RLC circuits, op-amps, etc.) that are capable of realizing these linear dynamics. However, this realization quickly becomes challenging when designing biomolecular controllers. This difficulty arises because (a) biomolecular controllers have to respect the structure of BioChemical Reaction Networks (BCRN), and (b) the quantities to be controlled (protein copy numbers or concentrations) cannot be negative (see [23] for positive integral control). Furthermore, the dynamics of biochemical reactions are inherently nonlinear. To achieve RPA, BCRN realizations of standalone Integral (I) controllers received considerable attention [24–29]. In previous work [26], the antithetic Integral (*a*I) feedback controller was introduced to realize integral action that ensures RPA. More recently, it was shown in [10] that the antithetic motif is necessary to achieve RPA in arbitrary intracellular networks with noisy dynamics. A detailed mathematical analysis of the performance tradeoffs that may arise in the *a*I controller is presented in [30,31], and optimal tuning is treated in [32]. Furthermore, practical design aspects, particularly the dilution effect of controller species, are addressed in [10,28]. Biological implementations of various biomolecular integral and quasi-integral controllers appeared in bacteria *in vivo* [7,9,10] and in *vitro* [14], and more recently in mammalian cells *in vivo* [15] and in yeast using the cyberloop *in silico* [33].

In the pursuit of designing high performance controllers while maintaining the RPA property, BCRN realizations of PI and PID controllers are starting to receive more focused attention [34–39]. Particularly in [34], a proportional component is separately appended to the antithetic integral motif via a repressing hill-type function to tune the transient dynamics and reduce the variance. The resulting PI controller follows the concept of Figure 1(d) where error and output feedback are used to build separate (but nonlinear) P and I components. Several successful attempts were carried out to devise BCRN realizations that approximate derivatives [40–44]. A BCRN realization of a full PID controller was reported in [36], where the authors introduced additional controller species to obtain a derivative component. The resulting PID controller uses error feedback (similar to the concept of Figure 1(c)) to build separate nonlinear P, I, and D components and successfully improves the dynamic performance in the deterministic setting. Using a different approach, [38] and [39] exploit the dual-rail representation from [24], where additional species are introduced to overcome the non-negativity challenge of the realized PID controller. The authors demonstrate the resulting improvement of the performance in the deterministic setting. On a different note, [37] analyzed the effects of separate proportional and derivative controllers on (bursty) gene expression models in the stochastic setting.

Interestingly, previous research in this direction shares two intimately related aspects. Firstly, the P, I, and D components are realized separately such that they enter the dynamics additively. This aspect is motivated by traditional PID controllers where the controller dynamics are constrained to be linear, and thus the three components have to be added up (rather than multiplied for example).

However, since feedback mechanisms in BCRNs are inherently nonlinear, there is no reason to restrict the controller to have linear dynamics and/or additive components. Secondly, the proposed designs introduce additional species to mathematically realize the controller, and thus making the biological implementation more difficult. To this end, we consider in this paper (more general) nonlinear PID controllers that do not have to be explicitly separable into their three (P, I and D) components. This allows controllers to involve P, I, and D architectures in one (inseparable) block as depicted in Figure 1(e) where both, error and output, feedbacks are allowed. The nonlinearity and inseparability features of the proposed PI and PID controllers provide more flexibility in the BCRN design and allows simpler architectures that do not require introducing additional species to the standalone integral controller. Next, we slightly increase the complexity (or order) of the controller designs by introducing up to two additional controller species. This provides more degrees of freedom for the controller and, as a result, offers a higher achievable performance. Furthermore, the higher the order of the controller, the more separable it is which facilitates the tuning of the PID gains by the biomolecular parameters. This hierarchical approach offers the designer a natural compromise between simplicity and performance enhancement. A rich library of biomolecular PID controllers of variable complexity is presented in this paper to offer the biologists a flexible wide range of designs that can be selected depending on the desired performance and application at hand.

## Results

### General framework for biomolecular feedback controllers

The framework for feedback control systems is traditionally described through block diagrams (e.g. Figure 1(a)). In this section, we lay down a general framework for feedback control systems where both the plant and the controller are represented by Biochemical Reaction Networks (BCRN). With this framework, the controller can either represent an actual biomolecular circuit or it can be implemented as a mathematical algorithm *in silico* [45–47] to regulate a biological circuit (through light for example [33,48]).

Consider a general plant, depicted in Figure 2, comprised of *L* species **X**:= {**X_1_**,…, **X_L_** }that react with each other through *K* reaction channels labeled as 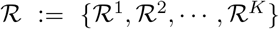. Each reaction 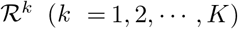 has a stoichiometry vector denoted by 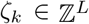 and a propensity function 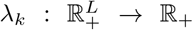. Let 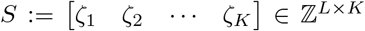 denote the stoichiometry matrix and let *λ*:= [*λ*_1_ *λ*_2_… *λ*_*K*_]^*T*^ denote the (vector-valued) propensity function. Then, the plant constitutes a BCRN that is fully characterized by the triplet 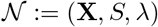 which we shall call the “open-loop” system.

**Figure 2:**
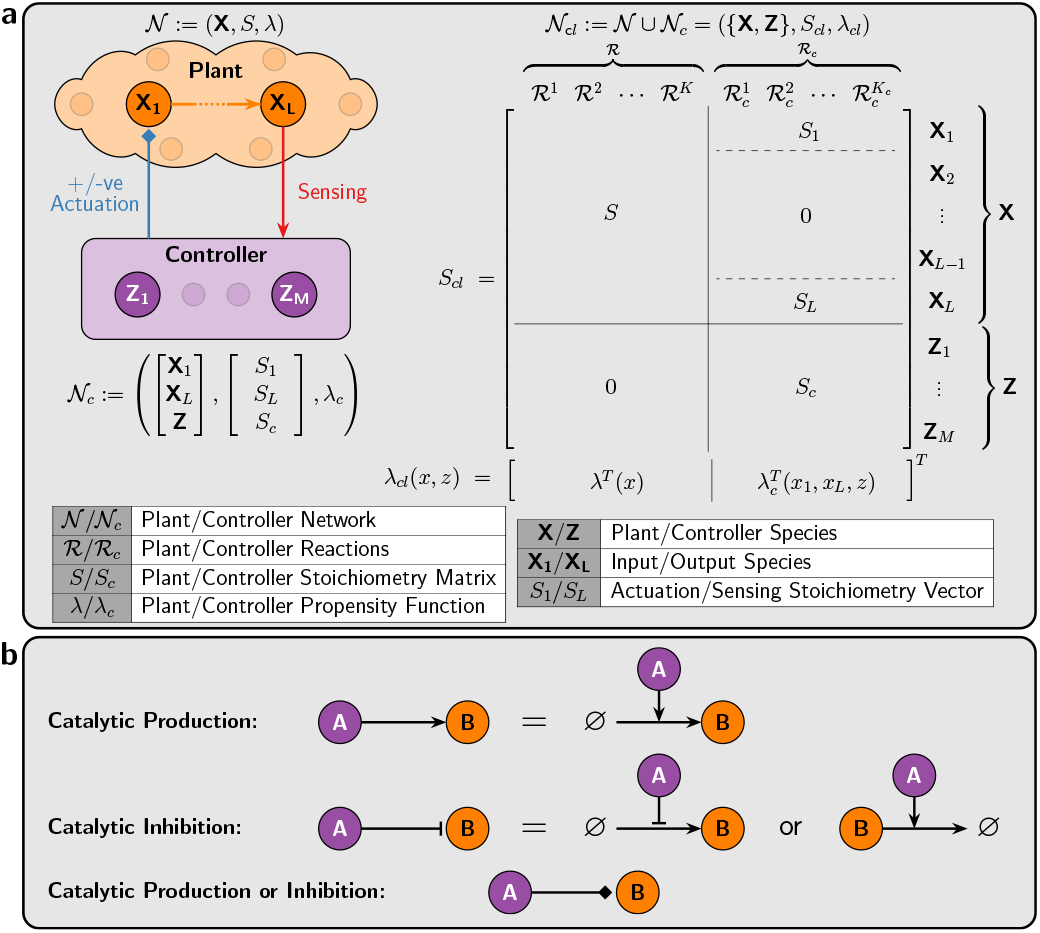
A framework for feedback control of Chemical Reaction Networks. **(a)** An arbitrary plant is comprised of *L* species {**X_i_**,… **X_L_**} reacting with each other. Species **X_L_**, by definition, is the output of interest to be controlled, while **X_1_** is assumed to be the only accessible input species that can be “actuated” (positively and/or negatively) by the controller network which is comprised of *M* species {**Z_1_**,…, **Z_M_**}. The closed-loop system, with stoichiometry matrix **S_cl_** and propensity function *λ_cl_*, denotes the overall feedback interconnection between the plant and controller networks. The partitioning of *S_cl_* and *λ*_*cl*_ describes the various components of the closed-loop network. **(b)** A description of the compact graphical notation that is adopted throughout the paper. Arrows directed toward species indicate catalytic productions, whereas T-shaped lines indicate catalytic inhibitions that encompass either repressive production or degradation. Note that the propensities of degradation reactions are considered to be either *kAB*/(*B* + *κ*) if two parameters (*k*, *κ*) are indicated on the arrow, or *ηAB* if only one parameter *η* is indicated on the arrow. Finally, diamonds indicate either production or inhibition.

The goal of this work is to design a controller network, denoted by 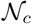, that is connected in feedback with the plant network 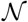, as illustrated in Figure 2(a), to meet certain performance objectives such as those mentioned in Figure 1(b). We assume that all the plant species are inaccessible by the controller except for species **X_1_** and **X_L_**. Particularly, the controller “senses” the plant output species **X_L_**, then “processes” the sensed signal via the controller species **Z**:= {**Z_1_**,…, **Z_M_**}, and “actuates” the plant input species **X_1_**. The controller species are allowed to react with each other and with the plant input/output species through *K_c_* reaction channels labeled as 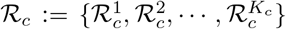. Let 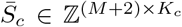 and 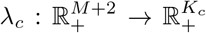 denote the stoichiometry matrix and propensity function of the controller, respectively. Since the controller reactions 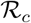 involve the controller species **Z** and the plant input/output species **X_1_**/**X_L_**, the stoichiometry matrix 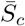 can be partitioned as

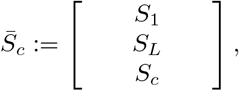

where *S*_1_ and 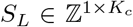 encrypt the stoichiometry coefficients of the plant input and output species **X_1_** and **X_L_**, respectively, among the controller reaction channels 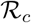. Furthermore,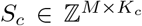 encrypts the stoichiom etry coefficients of the controller species **Z_1_**,…, **Z_M_**. Hence, the controller design problem boils down to de-signing *S*_1_, *S_L_*, *S_c_* and *λ*_*c*_. Note that, for simplicity, we consider plants with Single-Input-Single-Output (SISO) species. However, this can be straightforwardly generalized to Multiple-Input-Multiple-Output (MIMO) species by adding more rows to *S*_1_ and *S_L_*. Finally, the closed-loop system constitutes the open-loop network augmented with the controller network so that it includes all the plant and controller species **X_cl_**:= **X** ∪ **Z** and reactions 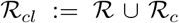. Thus, the closed-loop network, 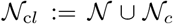, can be fully represented by the closed-loop stoichiometry matrix *S_cl_* and propensity function *λ*_*cl*_ described in Figure 2(a). We close this section, by noting that our proposed controllers range from simple designs involving *M* = 2 controller species, up to more complex designs involving *M* = 4 controller species.

### Antithetic Proportional-Integral (*a*PI) feedback controllers

Equipped with the BCRN framework for feedback control systems, we are now ready to propose several PI feedback controllers that are capable of achieving various performance objectives. All of the proposed controllers involve the antithetic integral motif introduced in [26] to ensure RPA. However, other additional motifs are appended to mathematically realize a Proportional (P) control action.

Consider the closed-loop network, depicted in Figure 3, where an arbitrary plant is connected in feedback with a class of controllers that we shall call *a*PI controllers. Observe that there are three different inhibition actions that are color coded. Each inhibition action gives rise to a single class of the proposed *a*PI controllers. Particularly, when no inhibition is present, we obtain the standalone antithetic Integral (*a*I) controller of [26] whose reactions are summarized in the left table of Figure 3. Whereas, *a*PI of Class 1 (resp. Class 2) involves the inhibition of **X_1_** by **X_L_** (resp. **Z_2_**), and *a*PI of Class 3 involves the inhibition of **Z_1_** by **X_L_**. Furthermore, each *a*PI class encompasses various types of controllers depending on the inhibition mechanisms that enter the controller network as actuation reactions. We consider three types of biologically-relevant inhibition mechanisms detailed in Figure 3: additive, multiplicative (competitive) and degradation. Considering all three *a*PI classes with the various inhibition mechanisms, Figure 3 proposes eight different *a*PI control architectures. Note that, it can be shown that a degradation inhibition in the case of *a*PI Class 3 would destroy the RPA property and is thus omitted. All of these controllers are compactly represented by a single general closed-loop stoichiometry matrix *S_cl_* and propensity function *λ*_*cl*_ depicted in Figure 3. The various architectures can be easily obtained by suitably selecting the functions *h*:= *h*^+^ – *h*^−^ and *g* from the tables of Figure 3. A theoretical linear perturbation analysis is carried out in Section S1.1 of the SI to verify the proportional-integral control structure of the proposed controllers. In fact, the analysis applies to any smooth function *h* which is monotonically increasing (resp. decreasing) in *z*_1_ (resp. *z*_2_, *x*_1_ and *x_L_*), and any smooth function *g* which is monotonically increasing (resp. decreasing) in *μ* (resp. *x_L_*).

**Figure 3:**
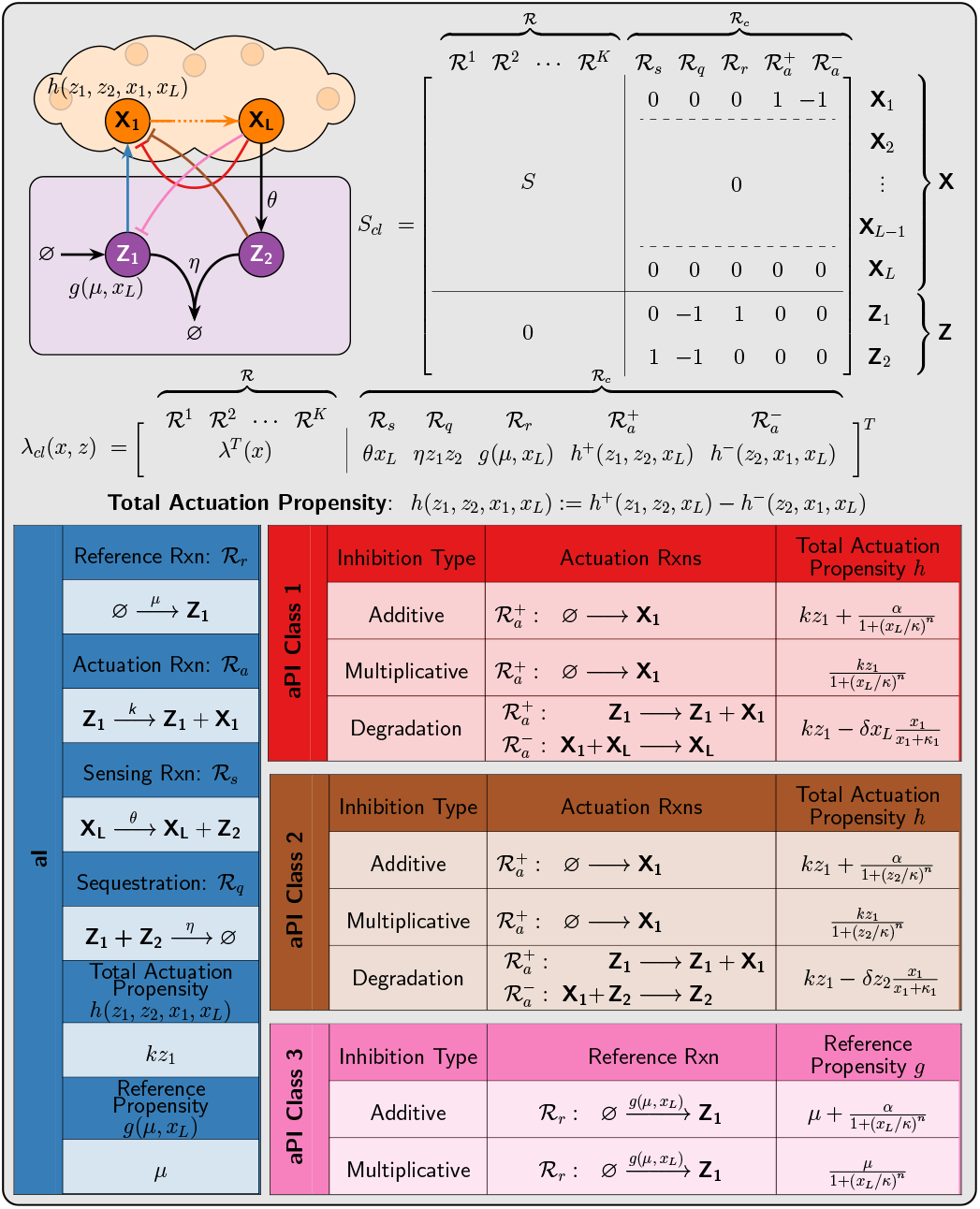
Antithetic Proportional-Integral (*a*PI) feedback controllers. Three different classes of *a*PI controllers are designed by appending the standalone *a*I controller with three inhibitions. Three biologically-relevant inhibition mechanisms are considered. **Additive Inhibition**: The inhibitor species produces the inhibited species separately at a decreasing rate. For instance, in the case of *a*PI Class 1 with additive inhibition, both **Z_1_** and **X_L_** produce **X_L_** separately, but **Z_1_** acts as an activator while **X_L_** acts as a repressor. This separate inhibition can be modeled as the production of **X_1_** as a positive actuation reaction 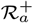 with an *additive* hill-type propensity given by 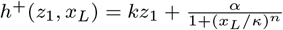, where *n*, *α* and *k* denote the hill coefficient, maximal production rate and repression coefficient, respectively. This *a*PI is the closest control architecture to [34] and [36], since the P and I components are additive and separable (see Figures 1(c) and (d)). **Multiplicative Inhibition**: The inhibitor *competes* with an activator over a production reaction. In the case of *a* PI Class 1 with multiplicative inhibition, **X_L_** inhibits the production of **X_1_** by **Z_1_**. This can be modeled as the production of **X_1_** with a *multiplicative* hill-type propensity given by 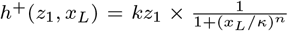. Observe that in this scenario, the Proportional (P) and Integral (I) control actions are inseparable, and the actuation reaction 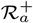 encodes both PI actions simultaneously. **Degradation Inhibition**: The inhibitor invokes a negative actuation reaction that degrades the inhibited species. For instance, in the case of *a*PI Class 1 with degradation inhibition, **Z_1_** produces **X_1_** (positive actuation reaction 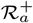), while **X_L_** degrades **X_1_** (negative actuation reaction 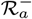). The dynamics can be captured by using a positive actuation with propensity *h*^+^(*z*_1_) = *kz*_1_ and a negative actuation with propensity 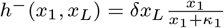. The total actuation propensity is defined as *h*(*z*_1_, *x*_1_, *x_L_*):= *h*^+^(*z*_1_) – *h*^−^(*x*_1_, *x_L_*). The three classes with different inhibition mechanisms give rise to eight controllers that are compactly represented by the closed-loop stoichiometry matrix *S_cl_* and propensity function *λ*_*cl*_ by choosing the suitable *h* and *g* functions from the tables.

### Deterministic steady-state analysis: Robust Perfect Adaptation (RPA) of *a*PI controllers

The deterministic dynamics of the closed-loop systems, for all the *a*PI controllers given in Figure 3 can be compactly written as a set of *Ordinary Differential Equations* (ODEs) given by

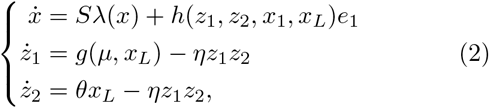

where 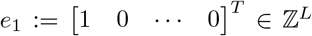. Note that the total actuation and reference propensities *h* and *g* take different forms for different *a*PI control architectures as depicted in Figure 3. The fixed point of the closed-loop dynamics cannot be calculated explicitly for a general plant; however, the output component (*x_L_*) of the fixed point solves the following algebraic equation

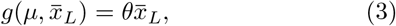

where over-bars denote steady-state values (if they exist), that is 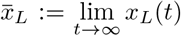. Two observations can be made based on (3). The first observation is that (3) has a unique nonnegative solution since *g* is a monotonically decreasing function in *x_L_*. The second observation is that (3) does not depend on the plant. As a result, if the closed-loop system is stable (i.e. the dynamics converge to a fixed point), then the output concentration converges to a unique set-point that is independent of the plant. This property is valid for any initial condition, and is referred to as *Robust Perfect Adaptation* (RPA). Particularly, for the *a*I and *a*PI controllers of Class 1 and 2, the reference propensity is *g*(*μ*, *x_L_*) = *μ*, and thus 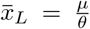. Furthermore, for the *a*PI of Class 3, 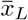 solves a polynomial equation of degree *n* + 1 (see Section S3 in the SI). In conclusion, all the proposed *a*PI controllers maintain the RPA property that is obtained by the antithetic integral motif, while introducing additional control knobs as extra degrees of freedom to enhance other performance objectives.

### Deterministic stability analysis & performance assessment of *a*PI controllers

To compare the stability properties of the various proposed *a*PI controllers, we consider a particular plant, depicted in Figure 4(a), that is comprised of two species **X_1_** and **X_2_**(*L* = 2). This plant may represent a gene expression network where **X_1_** is the mRNA that is translated to a protein **X_2_** at a rate *k*_1_. The degradation rates of **X_1_** and **X_2_** are denoted by *γ*_1_ and *γ*_2_, respectively. The closed-loop stoichiometry matrix and propensity function are also shown in Figure 4(a). Using the Routh-Hurwitz stability criterion [49], one can establish the exact conditions of local stability of the fixed point (Equation (S17) in Section S3 of the SI) for the various proposed *a*PI controllers. These conditions, once satisfied, guarantee that the dynamics locally converge to the fixed point.

**Figure 4:**
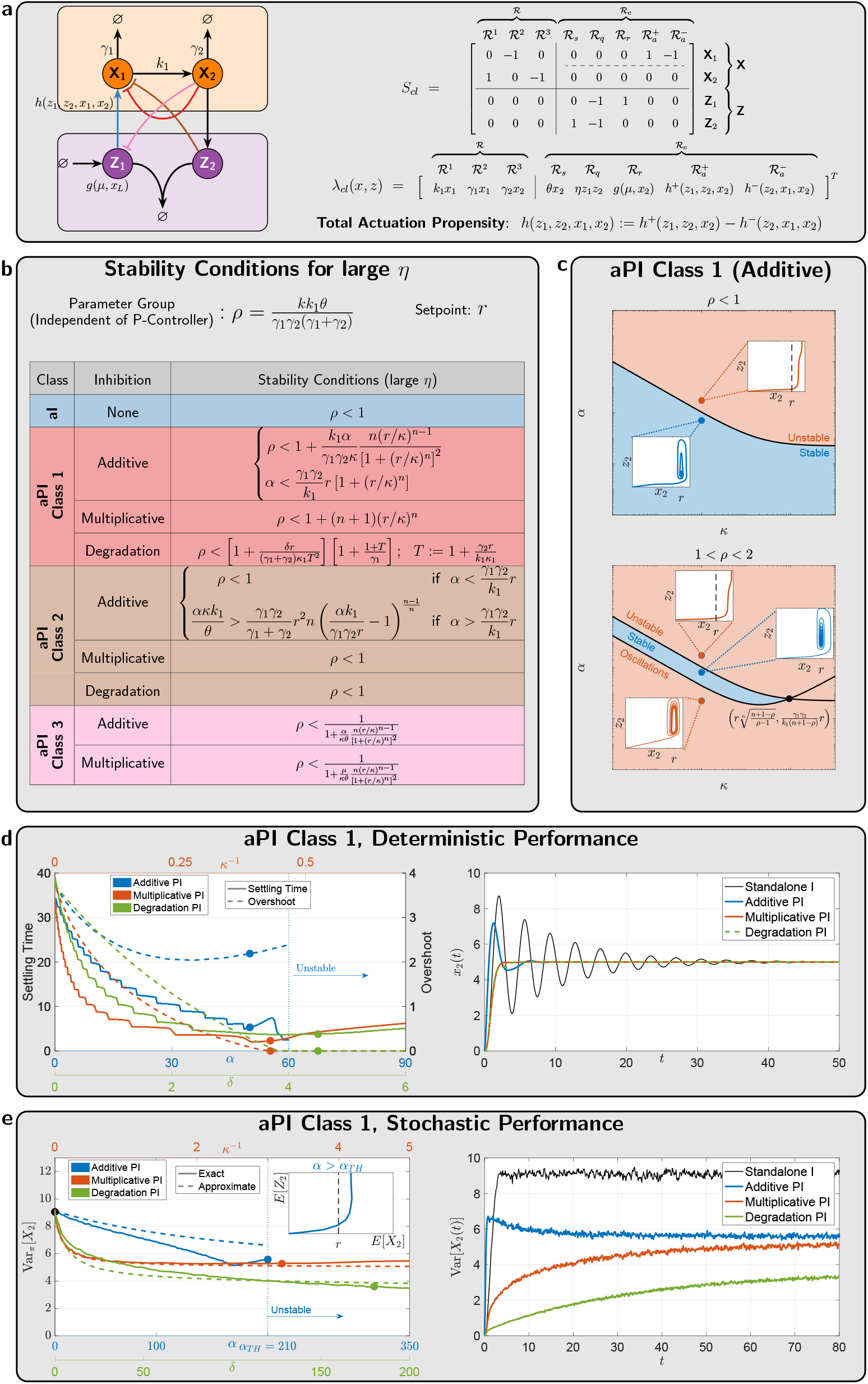
Performance of *a*PI feedback controllers. **(a)** Gene expression network controlled by *a*PI controllers. **(b)**Inequalities that need to be respected by the various controllers (with *η* being large enough) to guarantee closed-loop stability in the deterministic setting. Multiplicative and degradation inhibition mechanisms exhibit superior structural stability properties. **(c)** *a*PI controllers of class 1 with an additive inhibition mechanism, exhibit different stability properties for different ranges of the parameter group *ρ* (that depends solely on the plant and the standalone *a*I controller). In particular, for *ρ* < 2, the additive proportional control action can stabilize the dynamics, while for *ρ* > 2, it cannot stabilize without re-tuning the integral component. **(d)** Settling time and over-shoot for the output (**X_2_**) response as a function of controller parameters that are related to the appended proportional components. Multiplicative and degradation in-hibition mechanisms are capable of ameliorating the performance without risking instability as opposed to the additive inhibition mechanism. **(e)** Reduction of the output stationary variance with *a*PI controllers. The *a*PI controllers of Class 1 with all three inhibition mechanisms are capable of reducing the stationary variance of the output. This is demonstrated here via the simulations and the approximate formula shown in Table 1 as well. For additive inhibition, *α* has a threshold value *α_TH_* above which ergodicity is lost similar to the deterministic setting. Furthermore, observe that for values of *α* that are close to *α_TH_*, the analytic approximation is less accurate. In contrast, the multiplicative and degradation mechanisms are capable of reducing the variance without the risk of losing ergodicity, and the analytic approximation remains accurate. The numerical values of all the parameters can be found in Section S9 in the SI.

For the remainder of this section, we consider fast sequestration reactions, that is, *η* is large. Under this assumption, one can obtain simpler stability conditions that are calculated in Section S3 of the SI, and tabulated in Figure 4(b). The stability conditions are given as inequalities that have to be satisfied by the various parameters of the closed-loop systems. A particularly significant lumped parameter group is 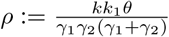 that depends only on the plant and standalone *a*I controller parameters. To study the stabilizing effect of the appended proportional (P) component, we fix all the parameters related to the plant and standalone *a*I controller (hence *ρ* is fixed), and investigate the effect of the other controller parameters related to the appended proportional component. By examining the table in Figure 4(b), one can see that, compared to the standalone *a*I, the *a*PI controller of Class 1 with multiplicative (resp. degradation) inhibition enhances stability regardless of the exact values of *κ* (resp. *δ*) and *n*. This gives rise to a structural stability property: adding these types of proportional components guarantees better stability without having to fine-tune parameters.

In contrast, although the *a*PI controller of Class 1 with additive inhibition may enhance stability, special care has to be taken when tuning *α*. In fact, if *α* is tuned to be larger than a threshold given by 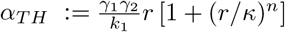, then stability is lost. Figure 4(c) elaborates more on this type of *a*PI controller. Three cases arise here. Firstly, if *ρ* < 1, that is the standalone *a*I already stabilizes the closed-loop dynamics, then the (*α*, *κ*)-parameter space is split into a stable and unstable region. In the latter (*α* > *α_TH_*), *z*_2_ grows to infinity, and the output *x*_2_ never reaches the desired set-point *r* = μ *ρ*/*θ*. Secondly, if 1 < *ρ* < 2, that is the standalone *a*I is unstable, then the (*α*, *κ*)-parameter space is split into three regions: (1) a stable region, (2) an unstable region with divergent response similar to the previous scenario where *ρ* < 1, and (3) another unstable region where sustained oscillations emerge as depicted in the bottom plot of Figure 4(c). Note that the closer *ρ* is to 2, the narrower the stable region is. Thirdly, for *ρ* > 2, the stable region disappears and thus this *a*PI controller has no hope of stabilizing the dynamics without re-tuning the parameters related to the standalone *a*I controller (e.g. *k* and/or *θ*). Clearly, multiplicative and degradation inhibitions outperform additive inhibition if stability is a critical objective. To this end, Figure 4(d) shows how the settling time and overshoot can be tuned by the controller parameters *α*, *κ*, and *δ* for additive, multiplicative, and degradation inhibitions, respectively. It is shown that with multiplicative and degradation inhibitions, one can simultaneously suppress oscillations (settling time) and remove overshoots. In contrast, a proportional component with additive inhibition can suppress oscillations but is not capable of removing overshoots as illustrated in the simulations of Figure 4(d) right panel. Furthermore, one can lose stability if *α* is increased above a threshold, as mentioned earlier. Nevertheless, for multiplicative and degradation inhibitions, increasing the controller parameters (*κ*^-1^, *δ*) too much can make the response slower but can never destroy stability.

It can be shown that the other two classes (2 and 3) are undesirable in enhancing stability. For instance, observe that for Class 2, the stability conditions are the same as the standalone *a*I controller (in the limit as *η* → *∞*) with the exception of the case of additive inhibition when 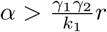. In this case, the inequality is structurally very different from all other stability conditions. In fact, the actuation via **Z_2_** dominates **Z_1_**, and hence **Z_2_** becomes responsible for the Integral (I) action instead of **Z_1_**. The detailed analysis of this network is not within the scope of this paper, and is left for future work. Finally, *a*PI controllers of Class 3 deteriorate the stability margin, since the right hand side of the inequalities is strictly less than one. However, this class of controllers can be useful for slow plants if the objective is to speed up the dynamics.

### Stochastic analysis of the *a*PI controllers: RPA& stationary variance

We now investigate the effect of the *a*PI controllers on the stationary (steady-state) behavior of the output species **X_L_** in the stochastic setting. Par-ticularly, we examine the stationary expectation 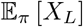 and variance Var_*π*_[*X_L_*]. The evolution of the expectations of the various species in the closed-loop network of Figure 3 are simply given by the differential equation 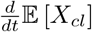. By substituting for the closed-loop stoichiometry matrix *S_cl_* and propensity function *λ_cl_* given in Figure 3, we obtain the following set of differential equations that describe the evolution of the expectations for an arbitraty plant.

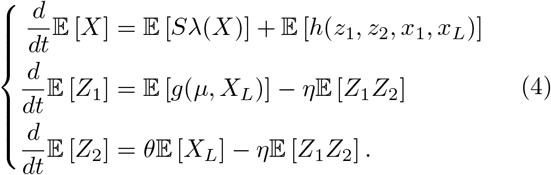

At stationarity, assuming that the closed-loop network is ergodic, the time derivatives are set to zero. Particularly, we have

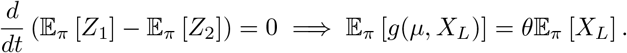

To achieve RPA at the population level (i.e. expectations), the stationary expectation 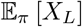 of the output species should not depend on the plant parameters. Clearly, this depends on the function *g*. In fact, if *g* is nonlinear in *X_L_*, then there is no guarantee that RPA is achieved because the nonlinearity couples higher order moments (that may depend on the plant parameters) with 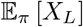. As a result, RPA is not guaranteed for the *a*PI controllers of Class 3 in the stochastic setting, although it is guaranteed in the deterministic setting. Nonetheless if *g* is affine in *X_L_*, then RPA is guaranteed (once again, assuming ergodicity). In particular, for the *a*PI controllers of Class 1 and 2, we have *g*(*μ*, *X_L_*) = *μ* and as a result 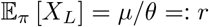. Clearly, for these classes of controllers, 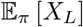 depends only on the control parameters *μ* and *θ* (like the deterministic setting), and thus RPA is ensured as long as the closed-loop network is ergodic.

Next, we examine the variance of the output species **X_L_**. Unfortunately, a general analysis for an arbitrary plant cannot be done. As a case study, we consider again the particular plant given in Figure 4(a) in feedback with the *a*PI controller of Class 1. Note that the subsequent analysis can be generalized to any (affine-linear) plant with mono-molecular reactions. Even for this particular plant, one cannot derive an exact expression for Var_*π*_[*X*_2_]. This is a consequence of the moment closure problem that stems from the inherent nonlinear nature of the antithetic motif (quadratic propensity: *ηz*_1_ *z*_2_) and the proportional control (propensity: *h*(*z*_1_, *z_2_*, *x*_1_, *x*_2_)). However, a tailored moment closure technique was proposed in [34] to give an approximate expression for Var_*π*_[*X*_2_] in the case of the *a*PI controller of Class 1 with additive inhibition and *n* = *k* = 1. This approximate technique exploits the fact that 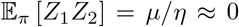 for large *η*; and as a result assumes that *Z*_2_ remains close to zero. Furthermore, a linearized approximation of the function *h* is also exploited to circumvent the moment closure problem. Extending this approximate technique to our more general controllers allows us to give a general (approximate) expression for Var_*π*_[*X*_2_] that encompasses all three types of inhibitions with an arbitrary hill coefficient *n* ≥ 1. The results are summarized in Table 1, where a general formula is given for any choice of *h*. One can easily see from the general expression in Table 1 that Var_*π*_[*X*_2_] is monotonically increasing in the integral gain *K_I_* ≈ *σ*_1_ and monotonically decreasing in the proportional gain *K*_*P*_1__ = *σ*_4_. This conclusion extends the results in [34] to more general proportional actuations involving different mechanisms of inhibitions and with cooperativity (*n* ≥ 1). Figure 4(e) demonstrates this stationary variance reduction via simulations and the approximate formula given in Table 1. Unlike additive inhibition, multiplicative and degradation inhibitions provide a structural property of decreasing the stationary variance of the output species**X_2_** without risking the loss of ergodicity (similar to the deterministic setting).

**Table 1:** An approximate analytic formula for the output stationary variance of a gene expression network regulated by the *a*PI controllers of Class 1 with various inhibition mechanisms. Recall that the total actuation propensity is defined as *h*:= *h*^+^ – *h*^−^, and its Jacobian is defined by 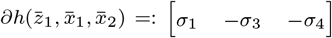 with *σ*_1_ > 0 and *σ*_3_, *σ*_4_ ≥ 0. Furthermore, recall from Equation (S2) in the SI that the proportional gain *K*_*p*_1__ = *σ*_4_ and the integral gain 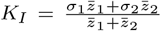. Since in the limit of large *η*, 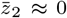 (refer to Section S7 in the SI), then *K_I_* ≈ *σ*_1_. Observe that the denominator of the variance expression is positive if the deterministic setting is stable (see Equation (S18) in the SI). Hence this expression is only valid when the deterministic setting is stable; otherwise, this approximation is meaningless. For additive inhibition (which is similar to the previous works in [34] and [36] with *κ* = *n* =1), *α* tunes the proportional gain separately. In fact, increasing *α* decreases the stationary variance as demonstrated in Figure 4(e) through both stochastic simulations and the approximate analytic formula. In contrast, for the case of multiplicative inhibition, tuning *κ* automatically tunes both the proportional and integral gains in a beneficial manner to the stationary variance. More precisely, decreasing *κ* increases the proportional gain *K*_*P*_1__ = *σ*_4_ and decreases the integral gain *K_I_* ≈ *σ*_1_, simultaneously. This has the effect of decreasing the variance without risking loss of stability as demonstrated in Figure 4(e). Finally, for degradation inhibition, increasing *δ* also increases the proportional gain and thus reduces the stationary variance as well.

| Stationary Variance $\text{Var}_\pi[X_2] \approx r \left[ \frac{(\gamma_1 + \gamma_2 + \sigma_3)(\gamma_1 \gamma_2 + \gamma_2 \sigma_3 + \sigma_1 k_1) + k_1 \left( \gamma_2 (\gamma_1 + \sigma_4) + \frac{k_1}{r} h^- \left( \frac{\gamma_2}{k_1} r, r \right) \right)}{(\gamma_1 + \gamma_2 + \sigma_3)(\gamma_1 \gamma_2 + \gamma_2 \sigma_3 + k_1 \sigma_4) - \sigma_1 k_1 \theta} \right]$ | | | | | |
| --- | --- | --- | --- | --- | --- |
| Controller | $h^+(z_1, x_2)$ | $h^-(x_1, x_2)$ | $\sigma_1 = K_I$ | $\sigma_3$ | $\sigma_4 = K_{P_1}$ |
| $aI$ | $kz_1$ | 0 | $k$ | 0 | 0 |
| $aPI$ Class 1 (Additive Inhibition) | $kz_1 + \frac{\alpha}{1 + (x_2/\kappa)^n}$ | 0 | $k$ | 0 | $\frac{\alpha}{r} \frac{n(r/\kappa)^n}{[1 + (r/\kappa)^n]^2}$ |
| $aPI$ Class 1 (Multiplicative Inhibition) | $\frac{kz_1}{1 + (x_2/\kappa)^n}$ | 0 | $\frac{k}{1 + (r/\kappa)^n}$ | 0 | $\frac{\gamma_1 \gamma_2}{k_1} \frac{n(r/\kappa)^n}{1 + (r/\kappa)^n}$ |
| $aPI$ Class 1 (Degradation Inhibition) | $kz_1$ | $\delta x_2 \frac{x_1}{x_1 + \kappa_1}$ | $k$ | $\delta r \frac{\kappa_1}{\left( \frac{\gamma_2}{k_1} r + \kappa_1 \right)^2}$ | $r \frac{\delta \gamma_2 / k_1}{\frac{\gamma_2}{k_1} r + \kappa_1}$ |

### Antithetic Proportional-Integral-Derivative feedback (*a*PID) control topologies

In this section, we append a Derivative (D) control action to the *a*PI (Class 1) controller of Figure 3 to obtain an array of *a*PID controllers depicted in Figure 5. The proposed *a*PID controllers range from simple second order (involving only two controller species **Z**_1_and **Z_2_**) up to fourth order (involving four controller species **Z_1_** to **Z_4_**). Furthermore the various controllers are categorized as two types: N-Type and P-Type. N-Type (Negative feedback) controllers are usually suitable for plants with positive gain (increasing the input yields an increase in the output), while P-Type (Positive feedback) controllers are usually suitable for plants with negative gains. This ensures that the overall control loops realize negative feedback. Note that one can easily construct hybrid PN-Type controllers, where the individual P, I and D components have different P/N-Types. This hybrid design is shown to be very useful for certain plants (see Figure 7(d) for example).

**Figure 5:**
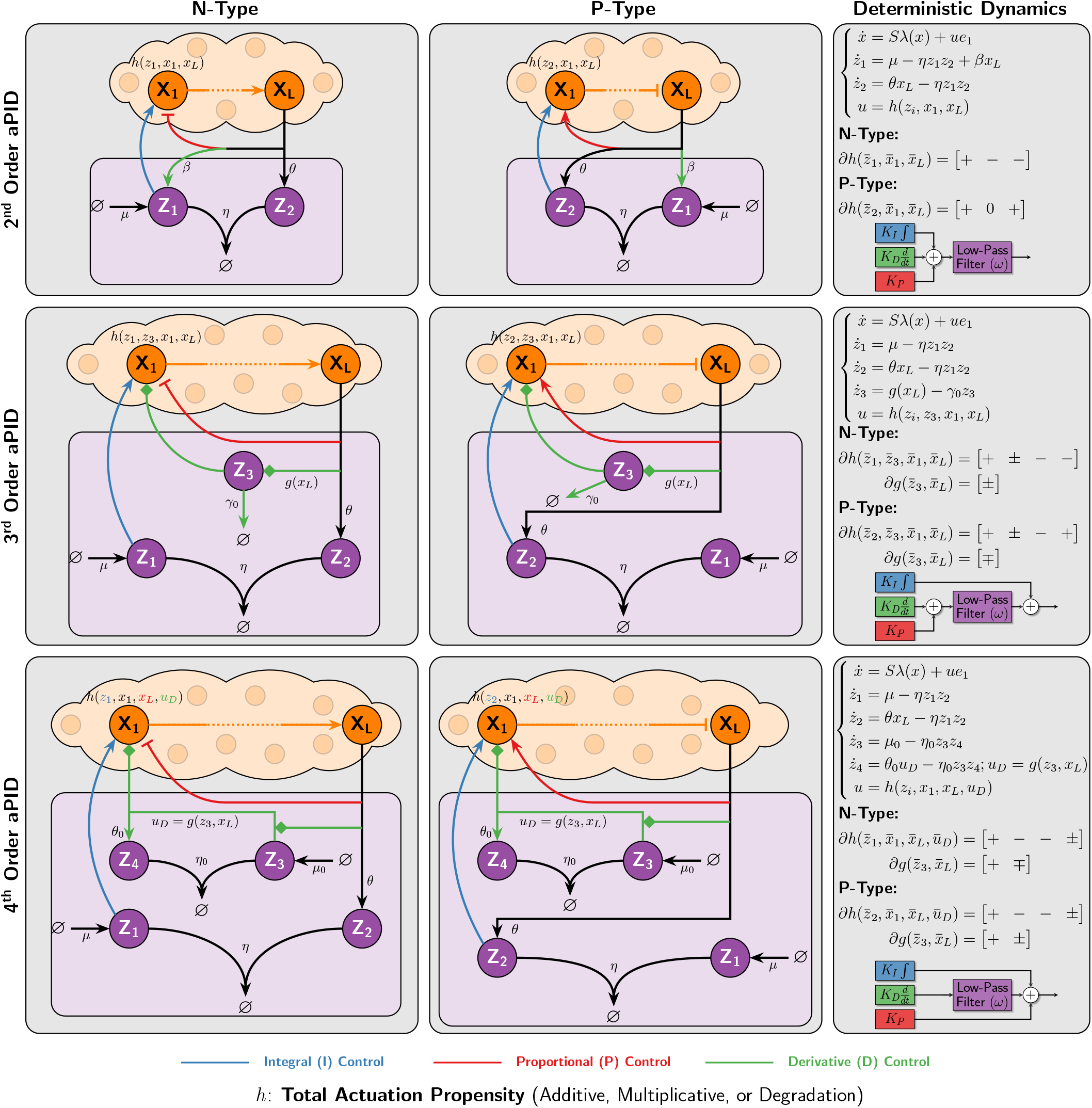
Antithetic Proportional-Integral-Derivative (*a*PID) feedback controllers. N-Type (Negative feedback) controllers are usually suitable for plants with positive gain (increasing the input yields an increase in the output), while P-Type (Positive feedback) controllers are usually suitable for plants with negative gains. The order of the controllers indicate the number of controller species ***Z_i_***. The second order *a*PID controller has the simplest design where no additional species are added to the *a*PI design, and only one reaction is added to produce **Z_1_** catalytically from **X_L_** at a rate *β* < *θ*. The third order *a*PID controller adds a single species to the *a*PI design. This intermediate species **Z_3_** is produced by the output **X_L_** and actuates the input species **X_1_**. These actions (indicated by the diamonds) are allowed to be either activations or both inhibitions. Finally, the fourth order *a*PID controller adds two species to the aPI design. These two species form an “antithetic differentiator” where **Z_3_** is constitutively produced at a rate *μ*_0_ and participates with **Z_4_** in a sequestration reaction with a rate *η*_0_. For the N-Type design, the derivative action enters the plant either by mutually producing **Z_4_** and **X_1_** at a rate proportional to *g*(*z*_3_, *x_L_*) (see Figure S4) such that *g* is monotonically increasing (resp. decreasing) in *z*_3_ (resp. *x_L_*), or by producing **Z_4_** while degrading **X_1_** at a rate proportional to *g*(*z*_3_, *x_L_*) (see Figure S5) such that *g* is monotonically increasing in both *z*_3_ and *x_L_*. Intuitively, the second and third order *a*PID controllers mathematically realize a derivative action using an incoherent feedforward loop from **X_L_** to **X_1_** via **Z_1_** and **Z_3_**, respectively; whereas the fourth order *a*PID controller realizes a derivative action by placing an additional antithetic integral motif in feedback with the plant and itself (**Z_3_** feeds back into **Z_4_**).

**Figure 6:**
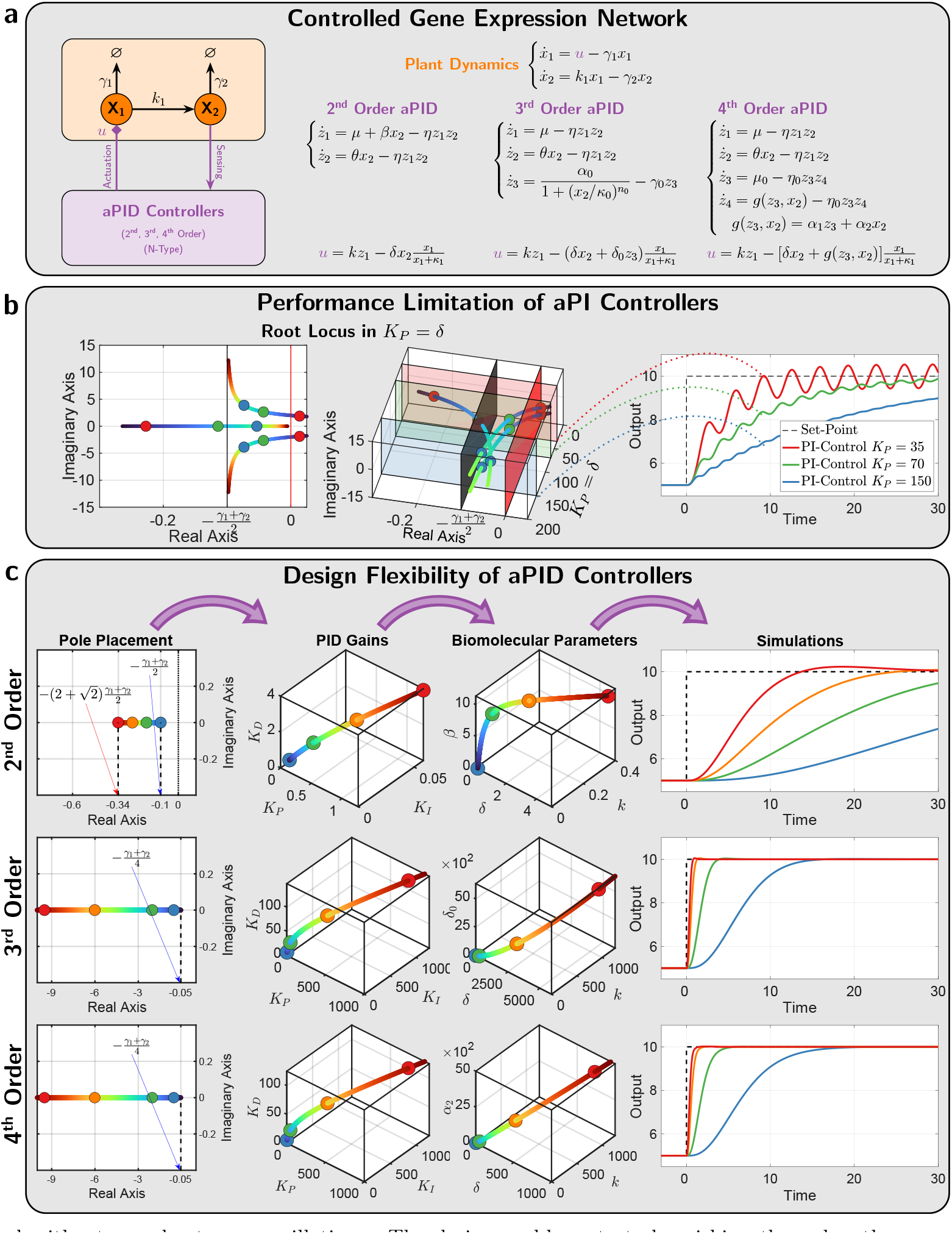
*a*PID control of a gene expression network. **(a) Closed-loop dynamics.** A gene expression network is controlled by the various N-Type *a*PID controllers of Figure 5. The deterministic dynamics and the overall control action *u* are shown here for each controller to explicitly specify the adopted propensity functions *h* and *g* in this example. **(b) Fundamental limitation of *a*PI controllers.** The left and middle plots demonstrate the same root locus of the linearized dynamics as *K_P_* is increased. The left plot depicts the complex plane, while the middle plot explicitly shows the complex plane together with the values of the proportional gain *K_P_* which is shown to be approximately equal to *δ*. These plots verify that two eigenvalues are confined within a small region close to the imaginary axis when *γ*_1_ and *γ*_2_ are small, and thus imposing a limitation on the achievable performance as demonstrated in the simulations shown in the right plot. **(c) Design flexibility offered by derivative control actions.** Exploiting all the components of the full *a*PID controllers offers more flexibility in achieving superior performance compared to the *a*PI controllers. This panel shows the steps of a pole-placement, control design problem where the four dominant poles are placed on the real axis of the left-half plane to ensure a stable and non-oscillating response with a minimal overshoot. The second order *a*PID exhibits a restriction on how far to the left the poles can be placed; whereas the higher order controllers can place the poles arbitrarily as far to the left as desired and thus achieving a response that is as fast as desi-red without overshoots nor oscillations. The design problem starts by picking the poles, then computing the PID gains (shown here) and cutoff frequency (not shown here), and finally computing the actual biomolecular parameters that allows us to obtain the nonlinear simulations to the right.

**Figure 7:**
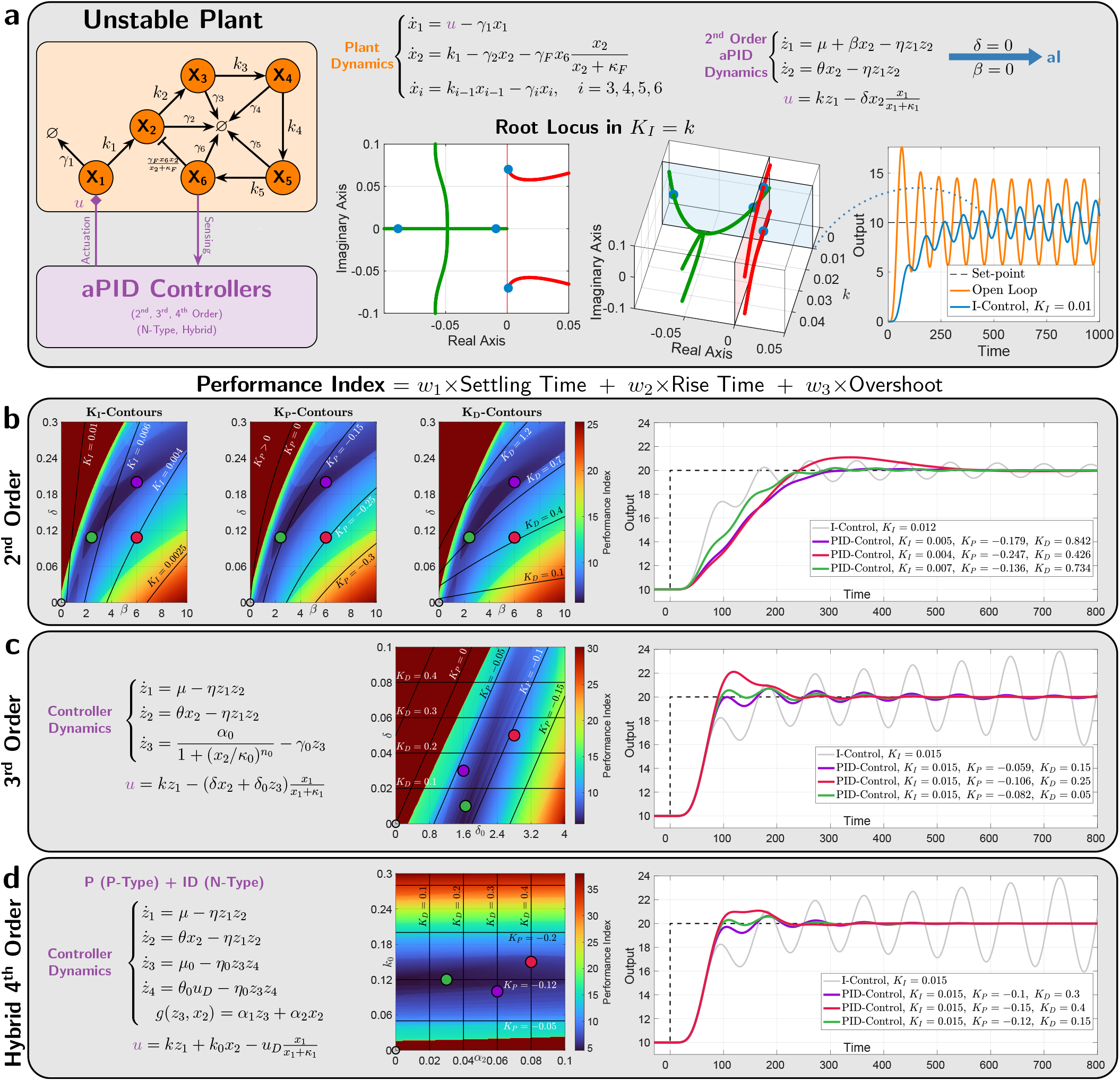
*a*PID control of an unstable and more complex plant. (a) Plant description. The plant considered here involves *L* = 6 species and embeds a negative feedback from **X_6_** to **X_2_** via an active degradation reaction. The underlying deterministic dynamics of the plant and the second order *a*PID controller are shown in this panel. It is demonstrated that the open loop is unstable (orange response), and integral control alone cannot stabilize the dynamics since two eigenvalues carry on a positive real part for any *k* < 0. **(b), (c) and (d) Performance of the various *a*PID controllers.** The intensity plots show the Performance Index over a range of biomolecular parameter values. These plots are overlaid with contours where the PID gains *K_P_*, *K_I_* or *K_D_* are constant. For the third and fourth order *a*PID in **(c)** and **(d)**, *K_I_* ≈ *k* and thus can be tuned separately with *k* which is held constant throughout this figure. For the fourth order *a*PID in **(d)**, the *K_P_*- and *K_D_*-contours are orthogonal to the *k*_0_- and *α*_2_-axes, respectively, and hence can also be tuned separately. For the third order *a*PID in **(c)**, the *K_D_*-contours are orthogonal to the *δ*-axis and hence *K_P_* can be tuned separately with *δ*_0_; whereas, the inseparability of the PD components forces the *K_P_*-contours to be oblique and thus *δ* tunes both *K_P_* and *K_D_* simultaneously. Finally, for the second order *a*PID in **(b)**, all three contours are not orthogonal to the axes and, as a result, all three PID gains have to be mutually tuned by the biomolecular parameters. This is due to the inseparability of all PID components. Note that each set of contours are displayed on a separate intensity plot here for clarity. Observe that the optimal performance for each controller is located in the dark blue regions where the proportional gains *K_P_* are negative. Three different examples, red, green and purple (along with the unstable standalone *a*I control in gray), are picked to demonstrate the achievable high performances depicted in the response plots to the right. For the second and third order *a*PID, negative *K_P_* can be achieved by properly tuning the biomolecular parameters without having to switch the topology from N-Type to P-Type. However, for the (separable) fourth order *a*PID controller, a hybrid design with N-Type ID and P-Type P can also achieve a negative *K_P_* which is critical for controlling this plant.

We begin with the N-Type second order design (first row of Figure 5) whose main advantage is its simplicity. Note that the rationale behind the various P-Type designs is similar. Intuitively, the antithetic integral motif is cascaded with an Incoherent FeedForward Loop (IFFL) to yield a PID architecture whose P, I and D components are inseparable as described in Figure 1(e). More precisely, the output species **X_L_** directly inhibits **X_1_** and simultaneously produces it via the intermediate species **Z_1_**. As a result, **Z_1_** simultaneously plays the role of both an intermediate species for the IFFL and the AIF control action. It is shown in Section S1.2.1 in the SI that this simple design embeds a low-pass-filtered PID controller. The N-Type third order design (second row of Figure 5) involves one additional controller species **Z_3_** to realize an IFFL that is disjoint from the antithetic motif. This yields an inseparable PD component appended to the separate I controller. It is shown in Section S1.2.2 in the SI that this design embeds a low-pass-filtered PD + I controller when *η* is large enough. In contrast, the N-Type fourth order design (third row of Figure 5) involves two additional controller species **Z_3_** and **Z_4_** to realize a completely separable PID control architecture. It is shown in Sections S1.2.3 and S1.2.4 in the SI that this design embeds a PI + low-pass-filtered D controller when *η* and *η*_0_ are large enough.

The key idea behind mathematically realizing the derivative component here is fundamentally different from the previous two designs. This controller realizes an “antithetic differentiator,” whereby the antithetic motif feeds back into itself: **Z_3_** feeds back into **Z_4_** via the rate function *g*(*z*_3_, *x_L_*). In fact, this idea is inspired by a mathematical trick in control theory (see Section S6 in the SI) which basically exploits an integral controller, in feedback with itself to implement a low-pass-filtered derivative controller. For this fourth order design, the derivative action can be achieved in two ways. One way is by mutually producing **Z_4_** and **X_1_** at a rate proportional to *g*(*z*_3_, *x_L_*) such that *g* is monotonically increasing (resp. decreasing) in *z*_3_ (resp. *x_L_*). This implementation is treated separately in Section S1.2.3 of the SI. The other way is by producing **Z_4_** while degrading **X_1_** at a mutual rate of *g*(*z*_3_, *x_L_*) such that *g* is monotonically increasing in both *z*_3_ and *x_L_*. This implementation is treated separately in Section S1.2.4 of the SI. Both designs have the same underlying PID control structure, but one might be easier to experimentally implement than the other.

### Deterministic analysis& properties of the various *a*PID controllers

It is straightforward to show that the set-point for the second-order design is given by 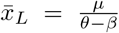 with the requirement that *β* < *θ*; whereas the set-point for both higher order designs are given by 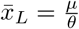. Furthermore, the effective PID gains, denoted by (*K_P_*, *K_I_*, *K_D_*), and cutoff frequency *ω* of the embedded low-pass filter for each of the proposed *a*PID controllers can be designed by tuning the various biomolecular parameters: *β*, *η*, *η*_0_, *γ*_0_, *μ*_0_ and the parameters of the propensity functions *h* and *g*. These functions depend on the specific implementation adopted. In particular, they can be picked in a similar fashion to the functions used to realize the three inhibition mechanisms (additive, multiplicative, or degradation) of the *a*PI controllers shown in Figure 3. In the subsequent examples, we use degradation inhibitions, but the other mechanisms can also be used.

Next, we demonstrate various properties of the proposed controller designs in the deterministic setting. The mappings between the effective PID parameters (*KxP*, *K_I_*, *K_D_*, *ω*) and the biomolecular parameters (*μ*, *θ*, *η*, *β*, *γ*_0_, *η*_0_,…) are given in S4 of the SI for each controller. It is fairly straightforward to go back and forth between the two parameter spaces. For control analysis, these mappings can compute the various PID parameters from the biomolecular parameters; whereas, for control design, these mappings can compute the various biomolecular parameters that achieve some desired PID gains and cutoff frequency. As a result, one can use existing methods in the literature (e.g. [50]) to carry out the controller tuning in the PID parameter space, and then map them to the actual biomolecular parameter space. Nevertheless, it is of critical importance to note that different controllers yield different coverage over the PID parameter space. For instance, for the fourth order design, there are enough biomolecular degrees of freedom to design any desired positive 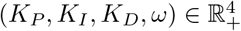. The lower the order of the controller, the fewer the biomolecular degrees of freedom, and hence the more constrained the coverage in the PID parameter space. For instance, for the third order design, the achievable PID parameters are constrained to satisfy *K_P_* ≤ *K_D_ω*. For the second order design, the constraint becomes even stricter. The details are all rigorously reported in Section S4 of the SI.

### Flexibility of *a*PID controllers

We first show the limitation of *a*PI controllers, and then demonstrate the flexibility that comes with an added derivative component. We also show that the higher order controllers exhibit more flexibility in shaping the transient response. Consider the controlled gene expression network depicted in Figure 6(a) where the ordinary differential equations of the various controllers are shown to explicitly specify the adopted propensity functions *h* and *g*. In this example, we consider both the P and D components acting on the input species **X_1_** as negative actuation via degradation reactions. We start by highlighting the fundamental limitation of *a*PI controllers alone (without a D) in Figure 6(b). Using simple root locus arguments (see Section S5.1), it is shown that as *K_P_* is increased, two complex eigenvalues of the linearized dynamics around the fixed point approach a vertical asymptote intersecting the real axis at 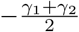, while one real eigenvalue approaches the origin (due to integral control). This is numerically demonstrated in the two root-locus plots of Figure 6(b), where *K_P_* ≈ *δ* (for a sufficiently small *κ*_1_). Clearly, the asymptotic limit is independent of all other parameters, including the integral gain *K_I_*. This analysis highlights a fundamental limitation of the *a*PI controller, because no matter how we tune *K_P_* and *K_I_*, two of the eigenvalues are constrained to remain close to the imaginary axis when *γ*_1_ and *γ*_2_ are small. In the time domain, these constraints impose either a slowly-rising response or a faster rising response but with lightly-damped oscillations, as illustrated in the simulation examples of Figure 6(b). In all cases, there is a fundamental limit on how fast the response can be. These limitations can be mitigated by appending a derivative control action via the various *a*PID controllers. To demonstrate this, we consider a design problem where the end goal is to achieve a fast response without oscillations and with minimal overshoot. This can be achieved by placing the eigenvalues far to the left on the real axis. Hence the design problem can essentially be translated to the following objective: place the four most dominant eigenvalues (or poles) at *s* = –*a* where *a* > 0 and make *a* sufficiently large. The design steps start by first (1) deciding where to place the poles *s* = –*a* for some desired *a*, then (2) computing the PID parameters so as to place the poles as desired; this can be achieved using equations (S46) in the SI, and finally (3) mapping the PID parameters to the actual biomolecular parameters using the formulas in Section S4. This is pictorially demonstrated in Figure 6(c) for each *a*PID controller. However, it is shown in Section S5.2 of the SI that the second-order *a*PID imposes a lower bound on the achievable poles given by 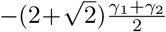 as demonstrated in Figure 6(c). As a result, with a second-order *a*PID, the performance can be made better than the *a*PI controller; however, the performance is also limited and cannot be made faster than a threshold, dictated by *γ*_1_ and *γ*_2_, without causing overshoots and/or oscillations. In contrast, it is also shown in Section S5.2 of the SI that the third- and fourth-order *a*PID controllers can make *a* as large as desired without any theoretical upper bound. This means that the added complexity of the higher order controllers are capable of shaping the response of the gene expression network freely and as fast as desired with no overshoots nor oscillations. This is also demonstrated in the simulations depicted in Figure 6(c).

### Deterministic performance of *a*PID control of a complex network

We consider a more complex plant to be controlled. The plant, comprised of *L* = 6 species, is depicted in Figure 7(a) where **X_i_** degrades at a rate *γ_i_* and catalytically produces **X_*i*+1_** at a rate *k_i_*. Furthermore, the output species *X_6_* feeds back into **X_2_** by catalytically degrading it at a rate *γ_F_*. This plant is adopted from [36]; however, to challenge our controllers in a way that demonstrates their features, the feedback degradation rate *γ_F_* is chosen to be larger than that reported in [36], which yields a plant that is unstable when operating in open loop, as shown in Figure 7(a). In fact, the root locus in the integral gain *K_I_* ≈ *k* (for large *η*) demonstrates that this plant cannot be stabilized with a standalone *a*I controller, that is no matter how we tune *k*, the response will remain unstable. It was shown in [36] that, for this plant, the P control is not useful. This is the case because the proportional gain *K_P_* was restricted to have a positive value. One of the nice features of our proposed second- and third-order *a*PID controllers is their ability to achieve negative proportional gains *K_P_* (see (S27) and (S32) in the SI) without having to rewire, that is without switching topologically from N-Type to P-Type. This is a consequence of the inseparability of the P component from other components (I and D for the second order, and D for the third order). In Figures 7(b) and (c) we show that, for this plant, tuning *K_P_* to be negative is critical to achieving high performance, whereby oscillations and overshoots are almost completely removed while maintaining a fast response. This is demonstrated using the intensity plots of a performance index that quantifies the overshoot, settling time, and rise time of the output response over a range of the relevant biomolecular controller parameters. With the completely separable fourth order *a*PID, the gains cannot be tuned to be negative; however, one can always switch between N-Type and P-Type topologies or even resort to hybrid designs where different PID components are of different P/N Types. Indeed, Figure 7(d) shows that by using a fourth-order hybrid *a*PID controller, very high performance can be attained.

To demonstrate the effectiveness of *a*PID control of high dimensional plants, we consider the control of cholesterol levels in the plasma depicted in Figure 8. High levels of cholesterol, particularly Low-Density Lipoprotein-Cholesterol (LDL-C), serve as a major trigger for cardiovascular disease. To circumvent that, the body possesses natural pathways to regulate cholesterol homeostasis [52]. However, these regulatory mechanisms have the tendency to fail with age leading to elevated levels of plasma cholesterol due to metabolic disorders and/or poor nutrition [53]. A whole body mathematical model of cholesterol metabolism is adopted from [51] and is briefly described in Figure 8 (see [51]for details). The mathematical model of the cholesterol network is comprised of 34 state variables (species) and 43 parameters whose values are taken from https://www.ebi.ac.uk/biomodels/ where the model is coded in SBML format (MODEL 1206010000). Two different disturbances are applied: (1) an increase in daily dietary intake; and (2) an increase in intestinal absorption. A second-order *a*PID controller is connected in feedback with the network to regulate the LDL-C level and keep it at 100 mg/dL despite the disturbances. A positive actuation is realized by the production of the Intestinal Cholesterol (IC) via **Z_1_** (blue arrow); whereas, a negative actuation is realized by the degradation of the IC via LDL-C (red T-shaped line). With positive integral actuation alone, the controller cannot achieve RPA because both disturbances increase the levels of LDL-C. Hence, without negative actuation (degradation of IC) the controller cannot restore the set-point. However, the *a*PID controller appends a negative actuation and is capable of achieving RPA of LDL-C levels and also of suppressing the overshoots and oscillations for both IC and LDL-C.

**Figure 8:**
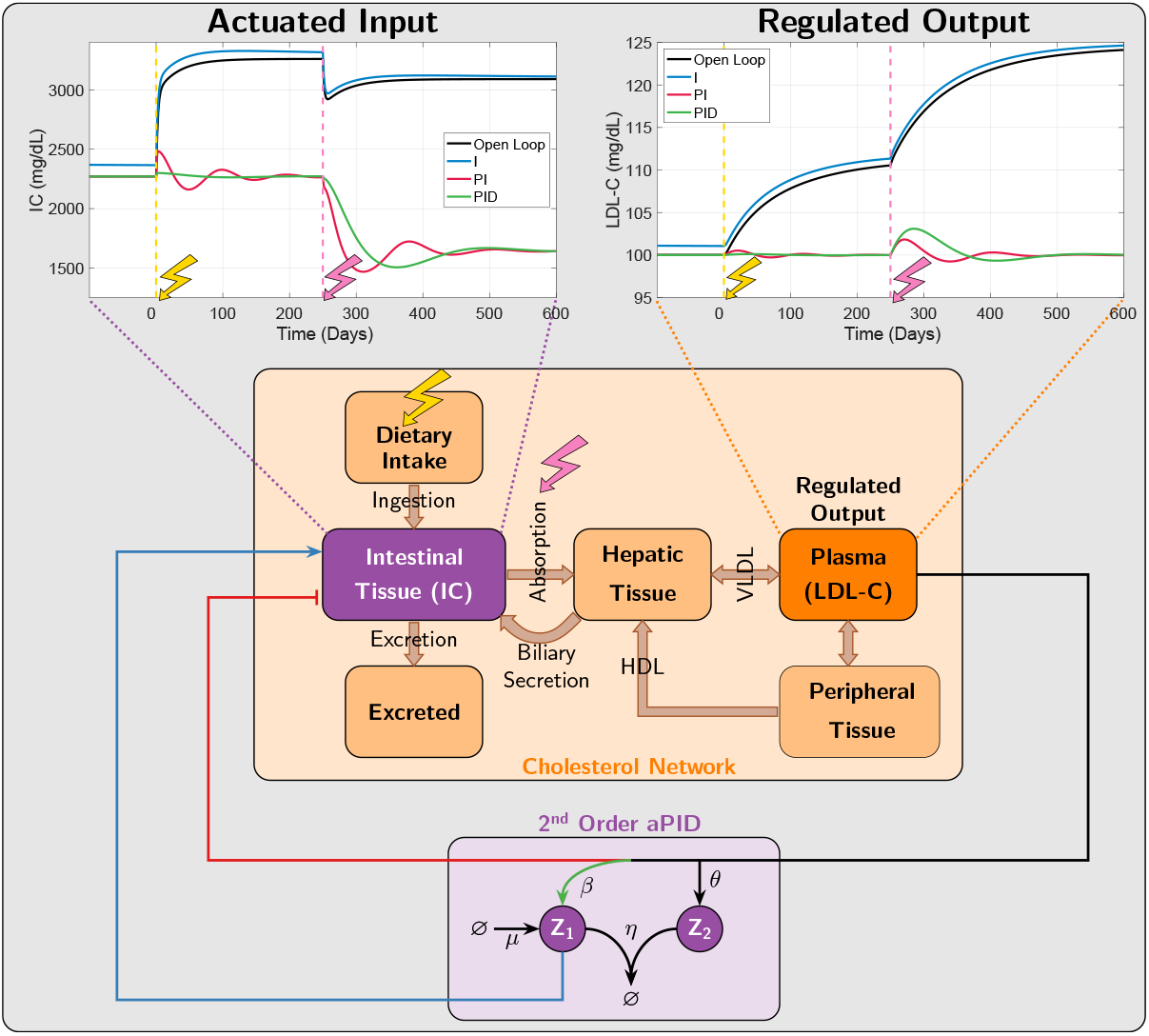
*a*PID Control of a whole-body model of Cholesterol metabolism. A whole-body model that describes the dynamics of cholesterol metabolism is adopted from [51]. The model tracks the flow of cholesterol, in its different forms, around the body. In summary, the Intestinal Cholesterol (IC) comes from the daily dietary intake or via biliary secretion from the liver (Hepatic Tissue). It is also locally synthesized in the intestine as well. The IC is either excreted from the body or absorbed and transported to the liver where it is then exported into the plasma via the Very-Low-Density Lipoproteins (VLDL). Excess cholesterol in the peripheral tissue is transferred to the liver via the High-Density Lipoproteins (HDL). Two exogenous disturbances are considered here: a 304 mg/day increase of dietary intake at *t*_1_ = 0 and 25% increase in absorption at *t*_2_ = 250 days. The first disturbance reflects a change in the daily diet, while the second reflects an increase of intestinal absorption efficiency. Both of these disturbances give rise to an increase in cholesterol levels in the plasma when no feedback control is applied as demonstrated in the black curve of the right plot. To implement the simplest (second order) *a*PID control, the IC is considered here to be the actuated input species, such that **Z_1_** produces IC, while the output species **LDL–C** degrades IC. The response of both the input and regulated output are shown here to demonstrate that an I alone (with positive actuation) is incapable of achieving RPA; whereas adding a PD (with negative actuation), not only achieves RPA, but also reduces oscillations.

### Effect of derivative control on the stationary variance

In this section, we examine the effect of the derivative component in the various *a*PID controllers on the cell-to-cell variability (e.g. stationary variance). We consider two plants: the gene expression network of Figure 6(a) and the six-species network of Figure 7(a). We fix the integral gain *K_I_* to be a constant and set the proportional gain *K_P_* to zero while we sweep the derivative gain *K_D_*. The biomolecular parameters that achieve these gains can be easily calculated using the mappings in Section S4 in the SI. The simulation results are depicted in Figure 9. Unlike the higher order *a*PID controllers, the second order *a*PID exhibits a hard upper limit on the achievable values of *K_D_* (see S27 in the SI). Figure 9 demonstrates that, for both plants, the third- and fourth-order *a*PID controllers are capable of reducing the stationary variance; whereas, the second-order *a*PID increases it. This conclusion is based on simulations only. Further stochastic analysis similar to [54] that exploits linear noise approximations is left for future work.

**Figure 9:**
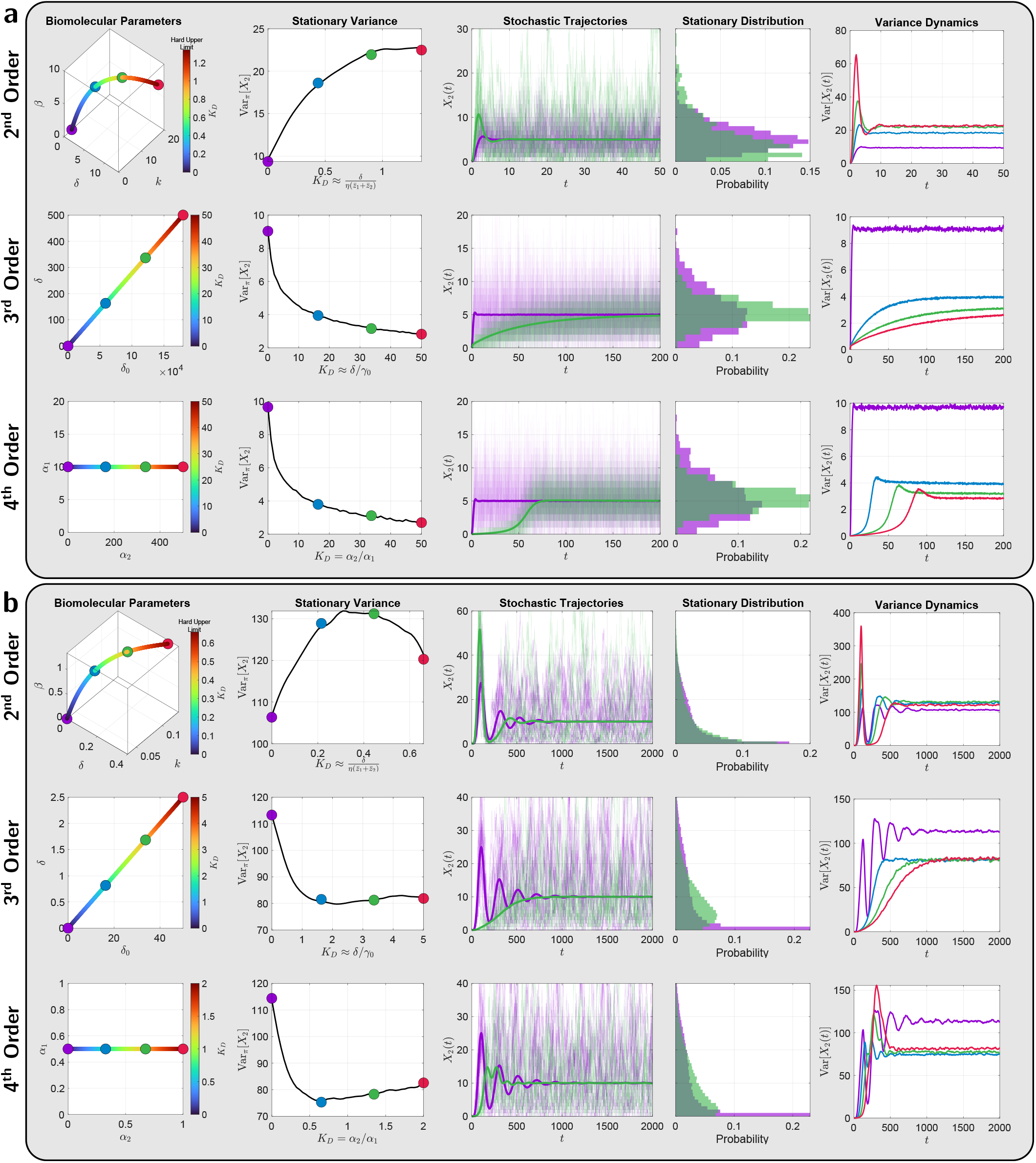
Stochastic performance of the *a*PID feedback controllers. In **(a)**, the plant is the gene expression network; whereas in **(b)**, the plant is the 6-Species network from Figure 7(a) but with stable open-loop dynamics (*γ_F_* is chosen to be smaller). For both networks, the biomolecular parameters are chosen such that the integral gain *K_I_* is constant and the proportional gain *K_P_* = 0, while the derivative gain *K_D_* is increased to track the effect of the derivative control action on the stationary variance of the output Var*π*[*X*_2_]. The first column shows the selected biomolecular parameters that achieve the desired values of *K_D_*, while the second column shows the stationary variance as a function of *K_D_*. The third and fourth columns show the stochastic (single-cell) trajectories and stationary distribution of the output for two particular values of *K_D_* (in green and purple). The fifth column shows the evolution of the variance in time for the four selected values of *K_D_*. These simulations demonstrate that the derivative control action increases the stationary variance of the output for second order *a*PID design, while it is capable of reducing the variance considerably for the higher order *a*PID designs. In fact, for the gene expression plant, the variance can be reduced to a level lower than the mean value = 5.

### Genetic circuit designs

Here we propose and describe a particular genetic design in E.Coli that realizes the third order *a*PID controller topology presented in Figure 5 (See Section S8 in the SI for another genetic design of the second order *a*PID controller). We also perform numerical simulations using biologically realistic parameters to demonstrate the effectiveness of the controllers in ameliorating the dynamic performance. The genetic circuit is depicted in Figure 10(a) where the controller circuit augments the I-control module (in blue), which is based on [10], with additional circuitry to implement additional P and D controls (in red and green). The only difference between our I-control module and that of [10] is the choice of the promoter P_RM_ driving the expression of the anti-*σ* factor (rsiW) which is activated by the transcription factor cI acting as a dual activator for P_RM_ and repressor for P_R_ (see [55], [56]). The P and D control modules are implemented via the Mflon protease which is capable of degrading the input species **X_1_**. The additional disturbance circuit (in yellow) serves as a source of external perturbation to the closed-loop circuit by degrading the regulated output **X_2_**. The set of ODEs describing the deterministic dynamics are also shown Figure 10(a) and the various parameters are chosen to reflect biologically realistic regimes and account for controller species dilution *δ_c_*(see Section S9 in the SI). Figure 10(b) shows the simulation results for I, PI and PID control. The responses are shown for a step change of setpoint *μ*/*θ*, which is tunable with HSL [10], at *t* = 8 hrs and for a step change of disturbance △, which is tunable with aTc, at *t* = 16 hrs. The simulations demonstrate that the full PID controller is capable of dramatically enhancing the stability and performance by not only shaping the transient dynamics but also reducing the steady-state error that can be incurred by the dilution effect (see [26], [10], [28]).

**Figure 10:**
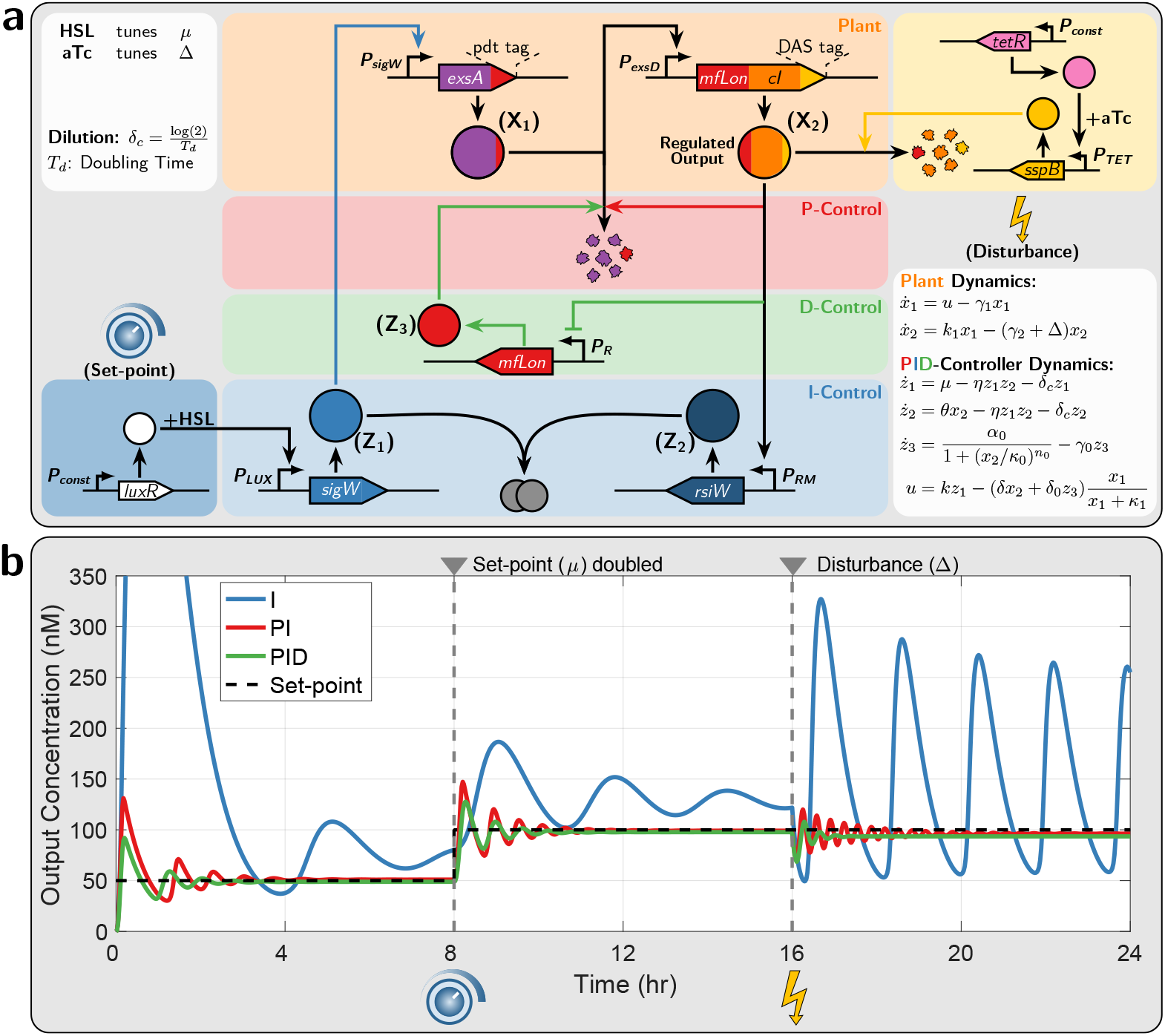
A genetic implementation of the third order *a*PID controller. **(a) Circuit design.** The genetic closed-loop circuit is comprised of the plant (in orange) involving two species **X_1_** and **X_2_**, the *a*PID controller involving **Z_1_**, **Z_2_** and **Z_3_** and the disturbance network (in yellow). The objective is to force the regulated output **X_2_** to track a tunable set-point, despite the injected disturbance, and enhance the transient dynamic response. The antithetic integral control is implemented via the sequestration between the *σ* factor (SigW denoted by **Z_1_**) and anti-*σ* factor (RsiW denoted by **Z_2_**) [10] that are driven by the promoters *P_LUX_* and *P_RM_* [55], respectively. The set-point is encrypted in the expression rate *μ* of **Z_1_** which is tunable with homoserine lactone (HSL). The plant is comprised of two genes. The first gene encodes *exsA* [57] fused to the degradation pdt tag (recognized by Mflon), and is driven by a SigW-responsive promoter *P_sig_W*. The second gene is driven by the ExsA-responsive promoter *P_exs_D* [57] and encodes the protease MfLon and the transcription factor *cI* fused to the ssrA(DAS) degradation tag. The disturbance circuit (in yellow) increases the degradation rate of **X_2_** by expressing *sspB* at a tunable rate, with anhydrotetracycline (aTc), which in turn recognizes the DAS tag in **X_2_** and sends it to the endogenous degradation machinery [58]. The MfLon in **X_2_** is capable of degrading **X_1_**, while the dual activator/repressor cI is capable of activating *P_RM_* and repressing *P_R_* [55]. The promoter *P_RM_* drives the integral control module, while *P_R_* drives another gene that expresses MfLon denoted by **Z_3_** which is also capable of degrading *X_1_*. Note that the protease MfLon encoded in **X_2_** and **Z_3_** implements an incoherent feedforward loop connected in feedback with the plant and thus realizing a PD-controller. **(b) Deterministic simulations.** Simulation of the closed-loop dynamics with I, PI and PID control. The plot shows the dynamic response of the regulated output **X_2_** to a step change in the set-point at *t* = 8 hrs and to a disturbance injection at *t* =16 hrs. The simulations are carried out using biologically realistic numerical values for the various parameters (see Section S9 in the SI).

### Experimental demonstration - Cyberloop implementation

To validate the performance benefits of the proposed *a*PID controllers, we implemented and tested our fourth order PID controller (presented in Figure 5) in a hybrid *in vivo - in silico* optogenetic platform [59] using the r*a*PID prototyping “Cyberloop” framework developed in [33]. This platform provides an interface (at singlecell resolution) between real biological circuits (*in vivo*) in cells placed under the microscope and stochastic computer simulation (*in silico*) of controllers via light stimulation and fluorescence measurement. Multiple cells can be targeted and observed individually in parallel on this platform. Under the cyberloop framework, at first, individual cellular outputs are observed and quantified via fluorescence imaging and subsequent image processing. The quantified value for each cell is then fed to a stochastic simulation [60] of controller reactions (one controller simulation per cell) which computes the light intensity (based on the controller species abundance) which the corresponding cell should be stimulated with. The light intensity data, once computed for every target cell in the microscope field of view, are sent to n specialized custom-built projection hardware [59] which then stimulates target cells with their corresponding light intensities in a parallel fashion, thus closing the control loop. This fluorescence measurement and subsequent light stimulation steps are repeated every fixed intervals providing us single-cell time-course data for the controller performance and output behavior.

In our cyberloop experiments, we used a target biological circuit genetically engineered in *Saccharomyces cere-visiae* (previously presented and used in [33,59]). As shown in Figure 11(a), this circuit includes an optoge-netic tool for gene expression regulation designed in such a way that the transcription rate in a target cell can be changed by varying blue light intensity which the cell is stimulated with. This provides a light control over nascent RNA abundance in the cell. The nascent RNAs are engineered with multiple stem loops which can bind with available fluorescent proteins in the cell and hence, they can be observed and quantified via fluorescence imaging under the microscope. The reader is referred to [59] and [33] for further technical details about this target circuit and the cyberloop framework, respectively.

**Figure 11:**
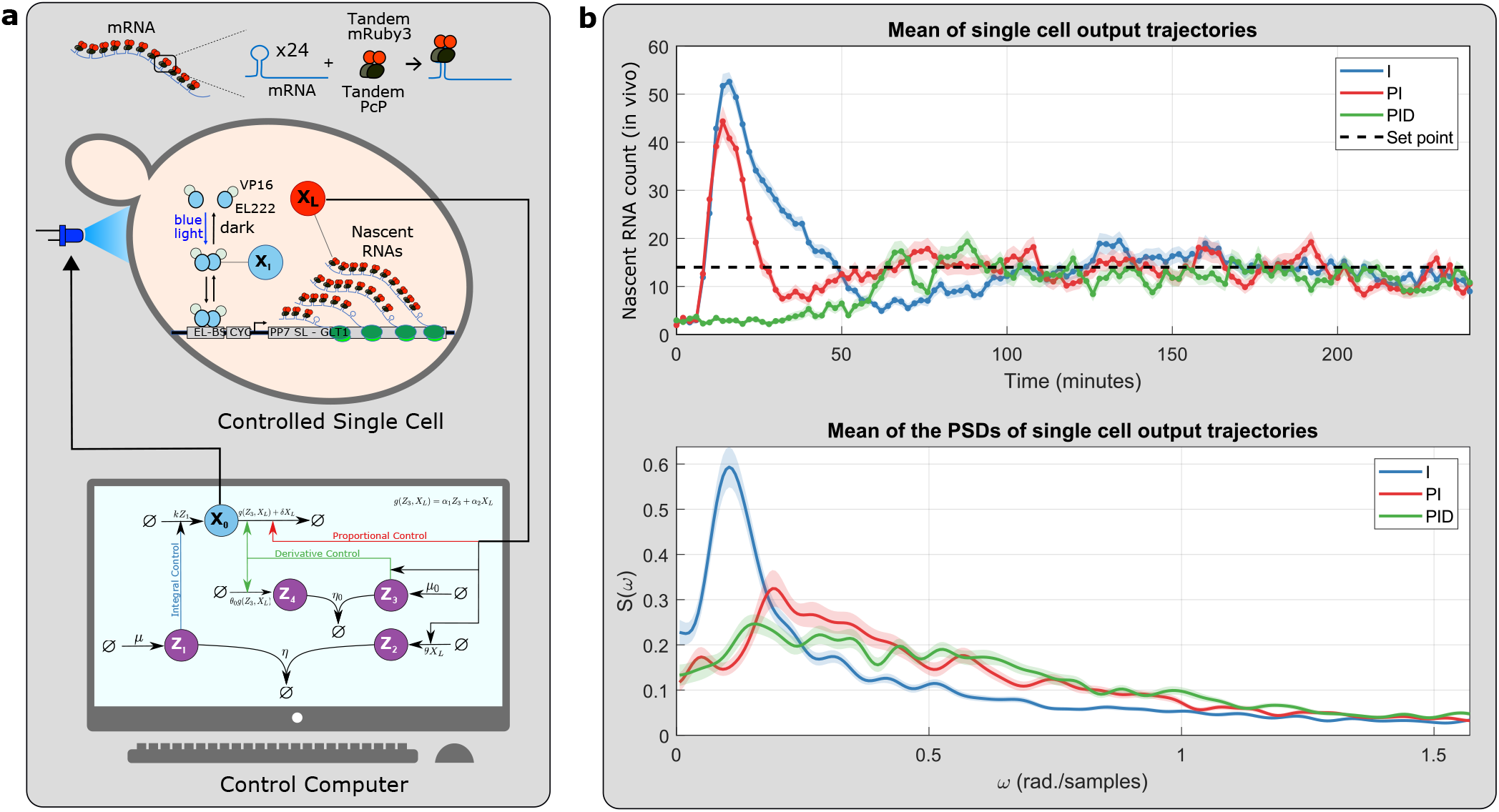
Experimental demonstration of the performance of *a*PID controllers in the Cyberloop platform [33] with a transcription circuit in *Saccharomyces cerevisiae*. **(a)** A Cyberloop implementation of the fourth order *a*PID controller. Using an optogenetic framework proposed in [33,59], an *in silico* stochastic simulation of the proposed fourth order *a*PID controller (**one controller per cell**) is interfaced with a real biological circuit/plant (in vivo) genetically engineered in *Saccharomyces cerevisiae* cells placed under a microscope. The circuit includes a blue-light optogenetic tool allowing light-stimulated regulation of transcription, making nascent RNAs (**X_L_**) as the output of interest to be controlled. These nascent RNAs (fused with fluorescent proteins) can be quantified via fluorescence imaging under the microscope and subsequent image processing in the control computer. The single-cell nascent RNA measurements are used to simulate stochastic dynamics of the controller network for each cell. Besides controller species (**Z_1_**, **Z_2_**, **Z_3_** and **Z_4_**), an additional *in silico* plant species X_0_ was added to the network to facilitate implementation of proportional and derivative control reactions. **X_0_** acts as an actuation species whose abundance defines the blue light intensity which the corresponding cell is then stimulated with using a custom-built projector set-up [59] attached to the microscope. This set-up allows one to stimulate and observe multiple cells in parallel. The single-cell measurements, controller simulations, and blue light intensity updates are done every 2 minutes interval. **(b)** The top plot shows the mean temporal response with the I-controller (across 168 cells), the PI-controller (across 128 cells) and the fourth-order PID-controller (across 131 cells). The shaded region represents mean ± standard error. This plot demonstrates the effectiveness of the PI-controller in reducing the oscillations of the mean response across the cells. It also demonstrates the added benefit of the PID-controller in reducing the overshoot as well. The bottom plot shows the mean Power Spectral Density (PSD) of various responses. The shaded region represents mean ± standard error. The PSD is useful in uncovering the stochastic oscillations on the single-cell level: a sharp peak in the PSD reveals the persistence of stochastic single-cell oscillations. The plot demonstrates the effectiveness of the PID controller in smoothing out the peak and thus considerably reducing the single-cell oscillations. The experimental parameters are provided in Supplementary Table S2. Source data are available in the source data file.

Following the approach in [33], we implemented our fourth order PID controller network. As shown in Figure 11(b), the PID controller was capable of reducing the oscillations on both the population and single-cell levels. This is demonstrated by plotting the time response of the population average, and the power spectral density (PSD) where sharp peaks indicate single-cell oscillations [61]. Particularly, the added derivative control action was capable of considerably enhancing the response by getting rid of the overshoot of the mean response across the cells, while simultaneously smoothing out the peak of the PSD and as a result suppressing the stochastic singlesingle oscillations.

### Alternative differentiators

In Figure 5, the antithetic integral motif is exploited to yield an antithetic differentiator; however, other integral motifs such as zero-order [62], [63] and auto-catalytic [25] integrators can also be similarly exploited as depicted in Figure S8 of the SI. These differentiators can be carefully appended to the *a*PI controllers of Class 1 (see Figure 3) to obtain an alternative set of *a*PID controllers depicted in Figure 12. Observe that these differentiators act on the concentration *x_L_* of the output species to approximate its derivative as a rate *u_D_*:= *g*(*z*_3_, *x_L_*). This is one of the differences between our differentiators and those proposed in [44] where the computed derivative is encoded as a concentration of another species. Having the computed derivative encoded directly as a rate rather than a concentration is particularly convenient for controllers with a fewer number of species. Another technical difference is that our differentiators realize a derivative with a first order low-pass filter; whereas, the differentiators in [44] realize derivatives with a second order low-pass filter due to the additional species introduced. We close this section by noting that it is also possible to replace the antithetic integral motif by other integrators to design yet another collection of PID controllers (see Figure S9 of the SI).

**Figure 12:**
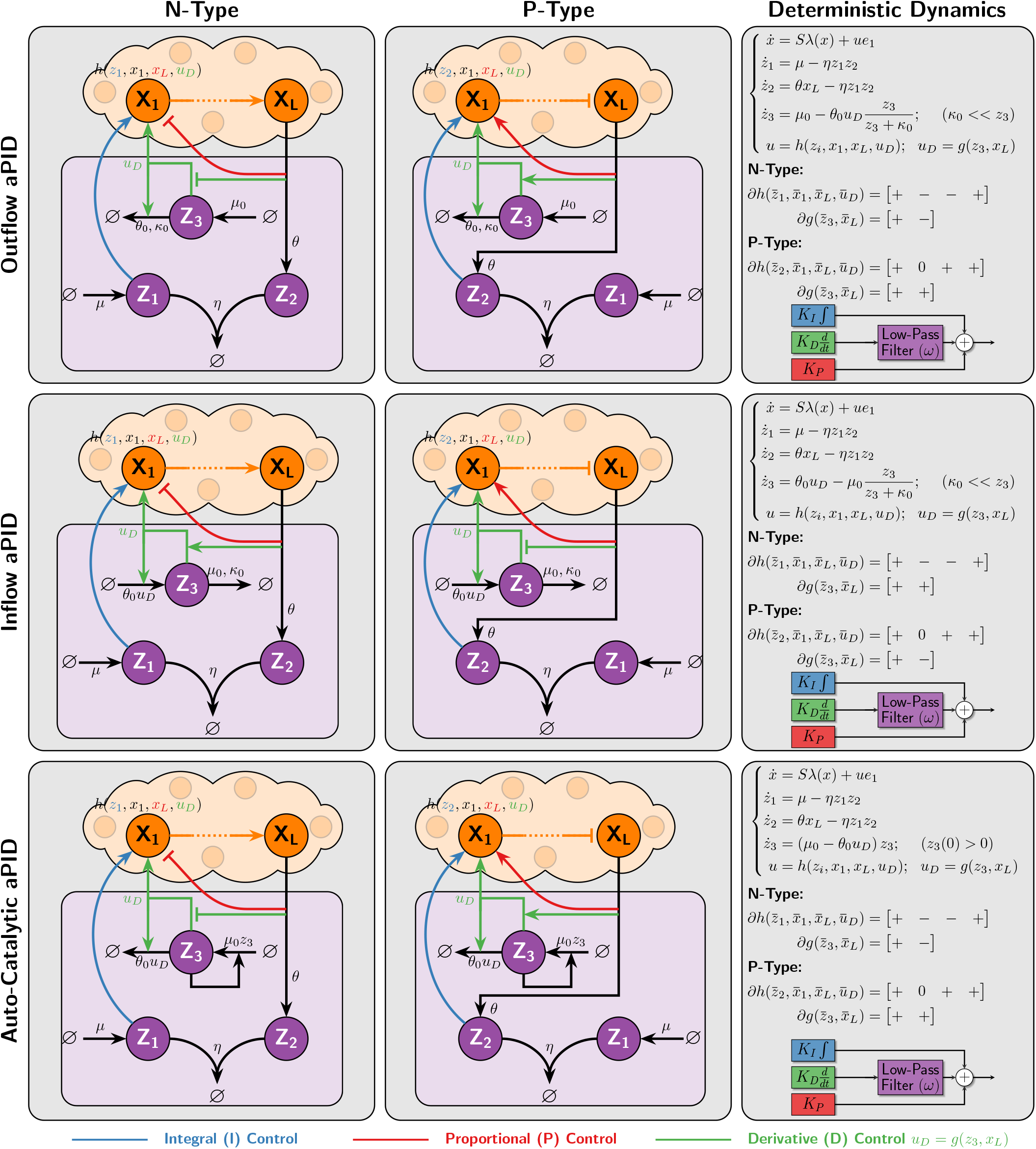
PID controllers using integral-based differentiators. Three differentiators are constructed based on three different integrators. The differentiators appended to the *a*PI controllers of Class 1 (see Figure 3) give rise to another collection of *a*PID controllers of both N- and P-Types. The Inflow and Outflow *a*PID controllers are based on integrators realized via zeroth-order degradation reactions [63], [62]. It is shown in Section S6 in the SI that if these degradation reactions are tuned to operate in a saturating regime (*κ*_0_ ≪ *z*_3_), then a low-pass filtered derivative action is mathematically realized. The difference between the outflow and inflow *a*PID controllers is that the feedback action *u_D_*:= *g*(*z*_3_, *x_L_*) which approximates the derivative of *x_L_* enters through a degradation and production reaction of the additional controller species **Z_3_**, respectively. In contrast, the auto-catalytic *a*PID controller is based on an auto-catalytic integrator [25] where the additional control species **Z_3_** produces itself. It is shown that for this component to properly function as a differentiator, the initial concentration of **Z_3_** has to be non-zero and *g* has to be designed such that *g*(0, *x_L_*) = 0 (see Section S6 in the SI).

## Discussion

This paper proposes a library of PID controllers that can be realized via biochemical reaction networks. The proposed PID designs are introduced as a hierarchy of controllers ranging from simple to more complex designs. This hierarchical approach that we adopt offers the designer a rich library of controllers that gives rise to a natural compromise between simplicity and achievable performance. At the lower end of the hierarchy, we introduce simple PID controllers that are mathematically realized with a small number of biomolecular species and reactions making them easier to implement biologically. As we move up in the hierarchy, more biomolecular species and/or reactions are introduced to push the limit on the achievable performance. More precisely, higher order PID controllers cover a wider range of PID gains that can be tuned to further enhance performance. Of course, this comes at the price of more complex designs making the controllers more difficult to implement biologically.

In this work, we start by introducing a library of PI controllers based on the antithetic integral motif [26] and an appended feedback control action where the input species is directly actuated by the output species. This is similar in spirit to previous works in [34] and [36] where the proportional control action enters the dynamics additively via a separate repressive production reaction. While this mechanism succeeds in enhancing the overall performance, we introduce other biologically-relevant mechanisms, for the P component, that are capable of achieving even higher performance without risking instability and further reducing the stationary variance (see Figure 4). However, it is shown rigorously and through simulations (see Figure 6) that a PI controller alone is limited, while adding a D component adds more flexibility. Interestingly, it is shown that the performance of a gene expression network can be arbitrarily enhanced with full PID controllers: the PID can be tuned to achieve an arbitrarily fast response without triggering any oscillations or overshoots. This example highlights the power of full PID control. Another nice feature of PID control is the availability of various systematic tuning methods in the literature (see [50] for example). Well-known design tools in control theory (such as the pole placement performed in Figure 6) can be exploited to perform the tuning in the PID parameter space instead of the biomolecular parameter space. Then the obtained PID parameters (PID gains and cutoff frequency) can be mapped by the formulas we derived (see Section S4) to the actual biomolecular parameters. This novel approach considerably facilitates the biomolecular tuning process. It is worth mentioning that the biomolecular tuning is the easiest for the fourth order *a*PID due to the separability of its components which allows tuning each PID gain separately with a different biomolecular parameter. In contrast, the lower order *a*PID controllers mix the various P, I, and D components and render them inseparable (see Figure 1 (e)) which results in each biomolecular parameter tuning multiple gains simultaneously. This is the price one has to pay for obtaining simpler designs. However, this can also be leveraged in some cases. For example, a single biomolecular parameter can tune both the integral and proportional gains simultaneously to enhance the dynamics and variance without risking instability (see the multiplicative *a*PI in Figure 4). This inseparability also offers a nice advantage where the proportional gains can be tuned to be negative without having to switch topologies from N-Type to P-Type. For certain plants, achieving negative gains is critical to achieve a high performance (see Figure 7).

We would like to point out that the proposed control structures are all designed based on linear perturbation analysis (see Section S1 in the SI). This is motivated by the rich set of existing tools to design and analyze linear control systems; whereas nonlinear control design and analysis is challenging and is often treated on a case-by-case basis. In the linearization, the PID structures are verified and hence the dynamics behave exactly like what is expected from classical PID control. However, full non-linear simulations are always carried out to back up the theoretical analyses and implications. Of course, the dynamical behavior of the nonlinear PID controllers may deviate from their linear counterparts when the dynamics are (initially) far from the fixed point. This is a limitation that we believe can serve as a good future research direction where small signal analysis should be extended to large signal analysis as well. Another possible future direction is to analyze the effects of dilution on the full *a*PID controllers in a similar fashion to the analysis carried out for I-controllers only in [10] and [28]. In fact, the simulations in Figure 10(b) show promising results on the roles of P and D controls in reducing the steady-state error incurred by dilution. Furthermore, in our work we lay down a general mathematical framework for biomolecular feedback control systems which can be used to pave the way for other possible controllers in the future. We believe that research along these directions helps building high performance controllers that are capable of reliably manipulating genetic circuits for various applications in synthetic biology and bio-medicine in the same way that PID controllers revolutionized other engineering disciplines such as navigation, telephony, aerospace, etc.

## Methods

### Yeast strain

Strain DBY96 from [59] was used for the cyberloop experiments in this study. All plasmids, strains and related details are summarized in the Key Resources Table in [59].

### Culture media and initialization

Yeast cell cultures were started from a −80°C glycerol stock at least 24 hours prior to the experiment, and were grown in an incubator (Innova 42R, New Brunswick) at 30°C in SD dropout medium (2% glucose, low fluorescence yeast nitrogen base (ForMedium), 5 g/L ammonium sulphate, 8 mg/L methionine, pH 5.8). The cell density was maintained at OD_600_< 0.2 in the incubator (30°C) for the last 12 hour leading to the experiment. Approximately 400 *μ*L of cell culture was centrifuged at 3000 RCF for 6 minutes, and then sufficient volume of supernatant was removed to get a concentrated culture with OD_600_ ~ 4.

### Microfluidic chip loading protocol

The microfluidic chip proposed in [59] was used in the cyberloop experiments in this study. As mentioned in [59], this chip is a single layer poly(dimethylsiloxane) (PDMS, Sylgard 184, Dow Corning, USA) device, attached to a cover glass (thickness: 150 mm, size: 24 mm x 60 mm, Menzel-Glaser, Germany). Before loading, the PDMS device and cover glass were rinsed with acetone, isopropanol, deionized water and dried using an air gun. The chip loading protocol in [59]was followed: Using a conventional pipette, 0.4 *μ*L of the concentrated cell solution (as described before) was loaded into each chamber of the clean and dried microfluidic chip. The cover glass was placed on top of the PDMS device and pressed down very gently, creating an electrostatic bond between the glass and the PDMS. The loaded microfluidic chip was placed onto a custom-built microscope holder. A syringe pump (Model no. 300, New Era Pump Systems, Inc.) was used to maintain 30 *μ*L/min of media flow through the loaded microfluidic chip. Cells were allowed to settle in the new conditions for 2 hours prior to the start of any experiment.

### Imaging and light delivery system

All image acquisitions were performed as described in [59]. Briefly, images were taken under an automated Nikon Ti-Eclipse inverted microscope (Nikon Instruments), equipped with a 40X oil-immersion objective (MRH01401, Nikon AG, Egg, Switzerland) and CMOS camera ORCA-Flash4.0 (Hamamatsu Photonic, Solothurn, Switzerland). Bright-field imaging was done using LED 100 (Märzhäuser Wetzlar GmbH& Co. KG) with diffuser and green interference filter placed in the light path. Fluorescence (mRuby3) imaging was done using Spectra X Light Engine fluorescence excitation light source (Lumencor, Beaverton, USA) with 550/15 nm LED line from the light source, 561/4 nm excitation filter, HC-BS573 beam splitter, 605/40 nm emission filter (filters and beam splitter acquired from AHF Analysetechnik AG, Tubingen, Germany). The microscope sample temperature was maintained at 30°C by enclosing the microscope with an opaque environment box set-up (Life Imaging Systems, Switzerland), which also shielded the cell sample from external light.

To achieve optogenetic stimulation with single-cell resolution under the microscope, a DMD (Digital Micromirror Device) based projection hardware developed in [59] was used. An additional neutral density filter (ND 1.3, 25 mm absorptive filter from Thorlabs) was placed in the light stimulation pathway to reduce blue light intensity reaching the cells. The microscope and DMD projector was operated using an open-source microscope control software YouScope [64].

### Image analysis

In this study, each of the cyberloop experiments was run for 4 hours duration with imaging/sampling done every 2 minutes. At every imaging step, 2 brightfield images above and below the focal plane (+/− 5 a.u. Nikon Perfect Focus System) were acquired, with an exposure of 100 ms each. These images were used for cell segmentation and tracking over the course of the experiment. For nascent RNA count quantification, 5 fluorescence images (Z stacks with step size ~ 0.5μm) were also captured, with an exposure of 300 ms each. The software tools developed in [59] and [33] were employed for cell segmentation, tracking and (nascent RNA) quantification. These image analysis software routines were run on MATLAB (MathWorks) environment.

### Stochastic simulation of proposed controllers

The *in silico* simulations of the proposed biomolecular controllers were run on MATLAB (MathWorks) environment as explained in [33]. At every sampling time, after running the cell quantification routine, the quantified cellular readout (nascent RNA count) was used to compute and update controller reaction network propensities for every tracked cell. Gillespie’s Stochastic Simulation Algorithm (SSA, [60]) was then employed to obtain the controller species abundance. These abundance values for individually tracked cells were used to compute blue light intensities which the corresponding cells were stimulated with via our custom-built DMD projector. The reader is referred to [33] for further technical details.

## Acknowledgments

This project has received funding from the European Research Council (ERC) under the European Union’s Horizon 2020 research and innovation programme (CyberGenetics; grant agreement 743269), and from the European Union’s Horizon 2020 research and innovation programme (COSY-BIO; grant agreement 766840).

## Competing Interests

ETH Zürich has filed a patent application on behalf of the inventors T.F., C.H.C., M.F. and M.K. that includes the designs described (application no. EP21187316.1).

## Notes

### Competing Interest Statement

ETH Zurich has filed a patent application on behalf of the inventors T.F., C.H.C., M.F. and M.K. that includes the designs described (application no. EP21187316.1).

### Summary of Updates

Firstly, we have proposed genetic designs in E.coli with specific biological parts that are capable of realizing two of our PID controllers. The biological parts detail the various choices of transcription factors, promoters and sequestration mechanisms that are capable of realizing the reaction-network control topologies which we propose in the manuscript. Furthermore, we carried out a careful literature review to extract biologically realistic values of the various parameters in the designed circuits. This allowed us to perform numerical simulations that demonstrate the effectiveness of our controllers in more realistic settings. Secondly, we have demonstrated the effectiveness of one of our more complex PID controllers in the Cyberloop, a hybrid experimental setup that we have recently developed. The Cyberloop serves as an ideal rapid prototyping platform for testing biomolecular controllers. In this hybrid setup, the controller is implemented in silico by carrying out a stochastic simulation of the controller topology in the computer, while the network to be controlled is a real biological genetic circuit. The in silico controller actuates the in vivo genetic circuit through light (exploiting light-responsive biological parts) and senses it using fluorescence microscopy. In the new version of our manuscript, we included experiments that clearly demonstrate the performance enhancements gained by one of our biomolecular PID controllers.

## References

[1] M. H. Khammash, “Robust steady-state tracking,” IEEE transactions on automatic control, vol. 40, no. 11, pp. 1872–1880, 1995.

[2] T.-M. Yi, Y. Huang, M. I. Simon, and J. Doyle, “Robust perfect adaptation in bacterial chemotaxis through integral feedback control,” Proceedings of the National Academy of Sciences, vol. 97, no. 9, pp. 4649–4653, 2000.

[3] D. Muzzey, C. A. Gómez-Uribe, J. T. Mettetal, and A. van Oudenaarden, “A systems-level analysis of perfect adaptation in yeast osmoregulation,” Cell, vol. 138, no. 1, pp. 160–171, 2009.

[4] H. El-Samad, J. Goff, and M. Khammash, “Calcium homeostasis and parturient hypocalcemia: an integral feedback perspective,” Journal of theoretical biology, vol. 214, no. 1, pp. 17–29, 2002.

[5] M. J. Dunlop, J. D. Keasling, and A. Mukhopadhyay, “A model for improving microbial biofuel production using a synthetic feedback loop,” Systems and synthetic biology, vol. 4, no. 2, pp. 95–104, 2010.

[6] J. A. Stapleton, K. Endo, Y. Fujita, K. Hayashi, M. Takinoue, H. Saito, and T. Inoue, “Feedback control of protein expression in mammalian cells by tunable synthetic translational inhibition,” ACS synthetic biology, vol. 1, no. 3, pp. 83–88, 2012.

[7] G. Lillacci, S. Aoki, D. Schweingruber, and M. Khammash, “A synthetic integral feedback controller for robust tunable regulation in bacteria,” BioRxiv, p. 170951, 2017.

[8] A. H. Ng, T. H. Nguyen, M. Gomez-Schiavon, G. Dods, R. A. Langan, S. E. Boyken, J. A. Samson, L. M. Waldburger, J. E. Dueber, D. Baker, et al., “Modular and tunable biological feedback control using a de novo protein switch,” Nature, vol. 572, no. 7768, pp. 265–269, 2019.

[9] H.-H. Huang, Y. Qian, and D. Del Vecchio, “A quasi-integral controller for adaptation of genetic modules to variable ribosome demand,” Nature communications, vol. 9, no. 1, pp. 1–12, 2018.

[10] S. K. Aoki, G. Lillacci, A. Gupta, A. Baumschlager, D. Schweingruber, and M. Khammash, “A universal biomolecular integral feedback controller for robust perfect adaptation,” Nature, p. 1, 2019.

[11] G. Lillacci, Y. Benenson, and M. Khammash, “Synthetic control systems for high performance gene expression in mammalian cells,” Nucleic acids research, vol. 46, no. 18, pp. 9855–9863, 2018.

[12] V. Hsiao, E. L. De Los Santos, W. R. Whitaker, J. E. Dueber, and R. M. Murray, “Design and implementation of a biomolecular concentration tracker,” ACS synthetic biology, vol. 4, no. 2, pp. 150–161, 2015.

[13] C. L. Kelly, A. W. K. Harris, H. Steel, E. J. Hancock, J. T. Heap, and A. Papachristodoulou, “Synthetic negative feedback circuits using engineered small rnas,” Nucleic acids research, vol. 46, no. 18, pp. 9875–9889, 2018.

[14] D. K. Agrawal, R. Marshall, V. Noireaux, and E. D. Sontag, “In vitro implementation of robust gene regulation in a synthetic biomolecular integral controller,” Nature communications, vol. 10, no. 1, pp. 1–12, 2019.

[15] T. Frei, C.-H. Chang, M. Filo, and M. Khammash, “Genetically engineered integral feedback controllers for robust perfect adaptation in mammalian cells,” bioRxiv, 2020.

[16] B. A. Francis and W. M. Wonham, “The internal model principle of control theory,” Automatica, vol. 12, no. 5, pp. 457–465, 1976.

[17] N. Minorsky, “Directional stability of automatically steered bodies,” Journal of the American Society for Naval Engineers, vol. 34, no. 2, pp. 280–309, 1922.

[18] R. Vilanova and A. Visioli, PID control in the third millennium. Springer, 2012.

[19] T. L. Blevins, “Pid advances in industrial control,” IFAC Proceedings Volumes, vol. 45, no. 3, pp. 23–28, 2012.

[20] J. Li and Y. Li, “Dynamic analysis and pid control for a quadrotor,” in 2011 IEEE International Conference on Mechatronics and Automation, pp. 573–578, IEEE, 2011.

[21] S. Bennett, “A brief history of automatic control,” IEEE Control Systems Magazine, vol. 16, no. 3, pp. 17–25, 1996.

[22] K. J. Åström and R. M. Murray, Feedback systems: an introduction for scientists and engineers. Princeton university press, 2010.

[23] C. Briat, “A biology-inspired approach to the positive integral control of positive systems: The antithetic, exponential, and logistic integral controllers,” SIAM Journal on Applied, Dynamical Systems, vol. 19, no. 1, pp. 619–664, 2020.

[24] K. Oishi and E. Klavins, “Biomolecular implementation of linear i/o systems,” IET systems biology, vol. 5, no. 4, pp. 252–260, 2011.

[25] C. Briat, C. Zechner, and M. Khammash, “Design of a synthetic integral feedback circuit: dynamic analysis and dna implementation,” ACS Synthetic Biology, vol. 5, no. 10, pp. 1108–1116, 2016.

[26] C. Briat, A. Gupta, and M. Khammash, “Antithetic integral feedback ensures robust perfect adaptation in noisy biomolecular networks,” Cell systems, vol. 2, no. 1, pp. 15–26, 2016.

[27] F. Xiao and J. C. Doyle, “Robust perfect adaptation in biomolecular reaction networks,” in 2018 IEEE Conference on Decision and Control (CDC), pp. 4345–4352, IEEE, 2018.

[28] Y. Qian and D. Del Vecchio, “Realizing ‘integral control’in living cells: how to overcome leaky integration due to dilution?,” Journal of The Royal Society Interface, vol. 15, no. 139, p. 20170902, 2018.

[29] C. C. Samaniego and E. Franco, “Ultrasensitive molecular controllers for quasi-integral feedback,” Cell Systems, 2021.

[30] N. Olsman, F. Xiao, and J. C. Doyle, “Architectural principles for characterizing the performance of antithetic integral feedback networks,” Iscience, vol. 14, pp. 277–291, 2019.

[31] N. Olsman, A.-A. Baetica, F. Xiao, Y. P. Leong, R. M. Murray, and J. C. Doyle, “Hard limits and performance tradeoffs in a class of antithetic integral feedback networks,” Cell systems, vol. 9, no. 1, pp. 49–63, 2019.

[32] M. Filo and M. Khammash, “Optimal parameter tuning of feedback controllers with application to biomolecular antithetic integral control,” in 2019 IEEE 58th Conference on Decision and Control (CDC), pp. 951–957, IEEE, 2019.

[33] S. Kumar, M. Rullan, and M. Khammash, “Rapid prototyping and design of cybergenetic single-cell controllers,” Nature communications, vol. 12, no. 1, pp. 1–13, 2021.

[34] C. Briat, A. Gupta, and M. Khammash, “Antithetic proportional-integral feedback for reduced variance and improved control performance of stochastic reaction networks,” Journal of The Royal Society Interface, vol. 15, no. 143, p. 20180079, 2018.

[35] A. Gupta and M. Khammash, “An antithetic integral rein controller for bio-molecular networks,” in 2019 IEEE 58th Conference on Decision and Control (CDC), pp. 2808–2813, IEEE, 2019.

[36] M. Chevalier, M. Gómez-Schiavon, A. H. Ng, and H. El-Samad, “Design and analysis of a proportional-integral-derivative controller with biological molecules,” Cell Systems, vol. 9, no. 4, pp. 338–353, 2019.

[37] S. Modi, S. Dey, and A. Singh, “Proportional and derivative controllers for buffering noisy gene expression,” in 2019 IEEE 58th Conference on Decision and Control (CDC), pp. 2832–2837, IEEE, 2019.

[38] N. M. Paulino, M. Foo, J. Kim, and D. G. Bates, “Pid and state feedback controllers using dna strand displacement reactions,” IEEE Control Systems Letters, vol. 3, no. 4, pp. 805–810, 2019.

[39] M. Whitby, L. Cardelli, M. Kwiatkowska, L. Laurenti, M. Tribastone, and M. Tschaikowski, “Pid control of biochemical reaction networks,” IEEE Transactions on Automatic Control, 2021.

[40] W. Halter, Z. A. Tuza, and F. Allgäwer, “Signal differentiation with genetic networks,” IFAC-PapersOnLine, vol. 50, no. 1, pp. 10938–10943, 2017.

[41] W. Halter, R. M. Murray, and F. Allgöower, “Analysis of primitive genetic interactions for the design of a genetic signal differentiator,” Synthetic Biology, vol. 4, no. 1, p. ysz015, 2019.

[42] C. C. Samaniego, G. Giordano, and E. Franco, “Practical differentiation using ultrasensitive molecular circuits,” in 2019 18th European Control Conference (ECC), pp. 692–697, IEEE, 2019.

[43] C. C. Samaniego, J. Kim, and E. Franco, “Sequestration and delays enable the synthesis of a molecular derivative operator,” in 2020 59th IEEE Conference on Decision and Control (CDC), pp. 5106–5112, IEEE, 2020.

[44] E. Alexis, C. C. Schulte, L. Cardelli, and A. Papachristodoulou, “Biomolecular mechanisms for signal differentiation,” bioRxiv, 2021.

[45] C. Guiver, H. Logemann, R. Rebarber, A. Bill, B. Tenhumberg, D. Hodgson, and S. Townley, “Integral control for population management,” Journal of Mathematical Biology, vol. 70, no. 5, pp. 1015–1063, 2015.

[46] C. Briat and M. Khammash, “Computer control of gene expression: Robust setpoint tracking of protein mean and variance using integral feedback,” in 2012 IEEE 51st IEEE Conference on Decision and Control (CDC), pp. 3582–3588, IEEE, 2012.

[47] C. Briat and M. Khammash, “Integral population control of a quadratic dimerization process,” in 52nd IEEE Conference on Decision and Control, pp. 3367–3372, IEEE, 2013.

[48] A. Milias-Argeitis, M. Rullan, S. K. Aoki, P. Buchmann, and M. Khammash, “Automated optogenetic feedback control for precise and robust regulation of gene expression and cell growth,” Nature communications, vol. 7, no. 1, pp. 1–11, 2016.

[49] J. J. Anagnost and C. A. Desoer, “An elementary proof of the routh-hurwitz stability criterion,” Circuits, systems and signal processing, vol. 10, no. 1, pp. 101–114, 1991.

[50] K. J. Åstrom and T. Hägglund, PID controllers: theory, design, and tuning, vol. 2. Instrument society of America Research Triangle Park, NC, 1995.

[51] M. T. Mc Auley, D. J. Wilkinson, J. J. Jones, and T. B. Kirkwood, “A whole-body mathematical model of cholesterol metabolism and its age-associated dysregulation,” BMC systems biology, vol. 6, no. 1, pp. 1–21, 2012.

[52] J. Luo, H. Yang, and B.-L. Song, “Mechanisms and regulation of cholesterol homeostasis,” Nature reviews molecular cell biology, vol. 21, no. 4, pp. 225–245, 2020.

[53] F. J. Féelix-Redondo, M. Grau, and D. Fernández-Bergées, “Cholesterol and cardiovascular disease in the elderly. facts and gaps,” Aging and disease, vol. 4, no. 3, p. 154, 2013.

[54] S. Modi, S. Dey, and A. Singh, “Noise suppression in stochastic genetic circuits using pid controllers,” PLoS Computational Biology, vol. 17, no. 7, p. e1009249, 2021.

[55] A. K. Brödel, A. Jaramillo, and M. Isalan, “Engineering orthogonal dual transcription factors for multiinput synthetic promoters,” Nature communications, vol. 7, no. 1, pp. 1–9, 2016.

[56] A. Hochschild and M. Lewis, “The bacteriophage *λ* ci protein finds an asymmetric solution,” Current opinion in structural biology, vol. 19, no. 1, pp. 79–86, 2009.

[57] T. Shopera, W. R. Henson, A. Ng, Y. J. Lee, K. Ng, and T. S. Moon, “Robust, tunable genetic memory from protein sequestration combined with positive feedback,” Nucleic acids research, vol. 43, no. 18, pp. 9086–9094, 2015.

[58] K. E. McGinness, T. A. Baker, and R. T. Sauer, “Engineering controllable protein degradation,” Molecular Cell, vol. 22, no. 5, pp. 701–707, 2006.

[59] M. Rullan, D. Benzinger, G. W. Schmidt, A. Milias-Argeitis, and M. Khammash, “An optogenetic platform for real-time, single-cell interrogation of stochastic transcriptional regulation,” Molecular Cell, vol. 70, no. 4, pp. 745–756, 2018.

[60] D. T. Gillespie, “Exact stochastic simulation of coupled chemical reactions,” Journal of Physical Chemistry, vol. 81, no. 25, pp. 2340–2361, 1977.

[61] A. Gupta and M. Khammash, “Frequency spectra and the color of cellular noise,” bioRxiv, 2020.

[62] X. Y. Ni, T. Drengstig, and P. Ruoff, “The control of the controller: molecular mechanisms for robust perfect adaptation and temperature compensation,” Biophysical journal, vol. 97, no. 5, pp. 1244–1253, 2009.

[63] M. H. Khammash, “Perfect adaptation in biology,” Cell Systems, vol. 12, no. 6, pp. 509–521, 2021.

[64] M. Lang, F. Rudolf, and J. Stelling, “Use of youscope to implement systematic microscopy protocols,” Current Protocols in Molecular Biology, vol. 98, no. 1, pp. 14–21, 2012.

