## Supplementary Information for "A Hierarchy of Biomolecular Proportional-Integral-Derivative Feedback Controllers for Robust Perfect Adaptation and Dynamic Performance"

**S2 Plants with Negative Gains: P-Type $a$PID Feedback Controllers** **14**

**S3 Derivation of the Stability Conditions of the $a$PI Controllers** **16**

**S4 Mappings between the PID and Biomolecular Parameter Spaces** **17**

**S8 A Genetic Design of the Second Order $a$PID Controller** **35**

**S9 Numerical Values** **35**

**S10 Useful Covariance Identities** **39**

**S11 Useful Expectation and Covariance Approximations** **40**

### S1 Linear Perturbation Analysis

In this section, we verify analytically that all the proposed controllers indeed involve Proportional (P), Integral (I) and Derivative (D) control actions. The analysis is carried out using linear perturbation theory where the linearized closed-loop dynamics are examined around the operating point (fixed point).

#### S1.1 Antithetic Proportional-Integral ($a$PI) Feedback Controllers

Consider an arbitrary network controlled by one of the $a$PI controllers depicted in Figure 3. The deterministic dynamics of the closed-loop systems, for all of these $a$PI controllers, can be compactly written as a set of *Ordinary Differential Equations* (ODEs) given by

$$\begin{cases} \dot{x} = f(x) + h(z_1, z_2, x_1, x_L)e_1 \\ \dot{z}_1 = g(\mu, x_L) - \eta z_1 z_2 \\ \dot{z}_2 = \theta x_L - \eta z_1 z_2, \end{cases} \tag{S1}$$

where $f(x) := S\lambda(x)$ and $e_1 := \begin{bmatrix} 1 & 0 & \cdots & 0 \end{bmatrix}^T \in \mathbb{Z}^L$. Let $\begin{bmatrix} \tilde{x}^T & \tilde{z}_1 & \tilde{z}_2 \end{bmatrix}^T$ denote the perturbation from the fixed point $\begin{bmatrix} \bar{x}^T & \bar{z}_1 & \bar{z}_2 \end{bmatrix}^T$ of (S1). To carry out a linear perturbation analysis we assume that the reference signal $\mu$ is allowed to slightly vary in time around a nominal constant reference $\bar{\mu}$. That is, we have

$$\tilde{x}(t) = x(t) - \bar{x}; \quad \tilde{z}_1(t) = z_1(t) - \bar{z}_1; \quad \tilde{z}_2(t) = z_2(t) - \bar{z}_2; \quad \tilde{\mu}(t) = \mu(t) - \bar{\mu}.$$

The linearized dynamics can thus be written as

$$\begin{cases} \dot{\tilde{x}} = A\tilde{x} + (\sigma_1 \tilde{z}_1 - \sigma_2 \tilde{z}_2 - \sigma_3 \tilde{x}_1 - \sigma_4 \tilde{x}_L)\, e_1 \\ \dot{\tilde{z}}_1 = \sigma_5 \tilde{\mu} - \sigma_6 \tilde{x}_L - \eta \bar{z}_2 \tilde{z}_1 - \eta \bar{z}_1 \tilde{z}_2 \\ \dot{\tilde{z}}_2 = \theta \tilde{x}_L - \eta \bar{z}_2 \tilde{z}_1 - \eta \bar{z}_1 \tilde{z}_2, \end{cases}$$

where the Jacobians are defined as $A := \partial f(\bar{x})$, $\partial h(\bar{z}_1, \bar{z}_2, \bar{x}_1, \bar{x}_L) =: \begin{bmatrix} \sigma_1 & -\sigma_2 & -\sigma_3 & -\sigma_4 \end{bmatrix}$, and $\partial g(\bar{\mu}, \bar{x}_L) =: \begin{bmatrix} \sigma_5 & -\sigma_6 \end{bmatrix}$, such that $\sigma_1, \sigma_5 > 0$ and $\sigma_2, \sigma_3, \sigma_4, \sigma_6 \geq 0$. The underlying control structure is most easily uncovered and visualized by drawing the block diagram of the linearized dynamics (refer to Box 1. A Primer on Block Diagrams). Taking the Laplace transforms yields

$$\begin{aligned} \tilde{x}_L(s) &= e_L^T \left(sI - \bar{A}\right)^{-1} e_1 \tilde{u}(s); \qquad \text{where} \quad \bar{A} := A - \sigma_3 e_1 e_1^T \\ \tilde{u}(s) &= \sigma_1 \tilde{z}_1(s) - \sigma_2 \tilde{z}_2(s) - \sigma_4 \tilde{x}_L(s) \\ \tilde{z}_1(s) &= \frac{\sigma_5 \tilde{\mu}(s) - \sigma_6 \tilde{x}_L(s) - \eta \bar{z}_1 \tilde{z}_2(s)}{s + \eta \bar{z}_2} \\ \tilde{z}_2(s) &= \frac{\theta \tilde{x}_L(s) - \eta \bar{z}_2 \tilde{z}_1(s)}{s + \eta \bar{z}_1}. \end{aligned}$$

Next, we express $\tilde{z}_1(s)$ and $\tilde{z}_2(s)$ in terms of $\tilde{\mu}(s), \tilde{x}_L(s)$ and the error defined as $\tilde{e}(s) := \tilde{\mu}(s) - \left(\frac{\theta + \sigma_6}{\sigma_5}\right) \tilde{x}_L(s)$. We have

$$\begin{aligned} \tilde{z}_1(s) &= \left[\sigma_5 \tilde{\mu}(s) - \sigma_6 \tilde{x}_L(s) + \frac{\eta \bar{z}_1 \sigma_5}{s} \tilde{e}(s)\right] \frac{1}{s + \eta\left(\bar{z}_1 + \bar{z}_2\right)} \\ \tilde{z}_2(s) &= \left[\theta \tilde{x}_L(s) - \frac{\eta \bar{z}_2 \sigma_5}{s} \tilde{e}(s)\right] \frac{1}{s + \eta\left(\bar{z}_1 + \bar{z}_2\right)}. \end{aligned}$$

Substituting for $\tilde{z}_1(s)$ and $\tilde{z}_2(s)$ in $\tilde{u}(s)$ and collecting similar terms yield

$$\tilde{u}(s) = \left[\sigma_1 \tilde{\mu}(s) + \frac{\eta\left(\sigma_1 \bar{z}_1 + \sigma_2 \bar{z}_2\right)}{s} \tilde{e}(s) - \frac{\sigma_2 \theta + \sigma_1 \sigma_6}{\sigma_5} \tilde{x}_L(s)\right] \frac{\sigma_5}{s + \eta\left(\bar{z}_1 + \bar{z}_2\right)} - \sigma_4 \tilde{x}_L(s).$$

#### Box 1. A Primer on Block Diagrams

In classical control theory, block diagrams are used to visually represent (deterministic) dynamical systems with inputs and outputs. In a fairly general setting, a dynamical system $\mathcal{M}$ can be written as a set of differential equations coupled with another set of algebraic equations given by

$$\mathcal{M} : \begin{cases} \dot{x} = f(x, u); \qquad x(0) = x_0 \\ y = g(x, u), \end{cases}$$

where $x$ is called the state variable with initial condition $x_0$, $u$ is the input, and $y$ is the output. Note that, for simplicity, $x, u$ and $y$ are all considered to be scalar functions of time (scalar signals); however, the extension to vector-valued signals is straightforward. One can think of $\mathcal{M}$ as a dynamical mapping that maps the input signal $u$ to the output signal $y$. In the rest of this box, we set the initial condition to zero for simplicity (and without loss of generality). This dynamical system $\mathcal{M}$ can be represented as a block diagram depicted in Panel a. This block takes $u$ as an input indicated by the inward arrow, and yields $y$ as the output indicated by the outward arrow. Note that inputs to a block are not affected by the block itself, only outputs are affected. Outputs can serve as inputs to other blocks, and inputs can be incoming as feedback from the output of other blocks (see Figure 1(a) for example). As a result, one of the nice features of a block diagram is to decompose the overall dynamics into multiple modularized sub-dynamical systems, each having a specialized operation.

Block diagrams are especially useful for linear dynamical systems such as the system to the left in Panel b. In this system, the input-output relationship, in the time domain, is given by a linear differential (and algebraic) equation, where $\omega_c$ is a constant. This relationship can be equivalently expressed in the Laplace domain, by taking the Laplace transforms. With slight abuse of notation, let $x(s), u(s)$ and $y(s)$ denote the Laplace transforms of $x(t), u(t)$ and $y(t)$, respectively, with $s$ being the Laplace variable. Note that we drop $t$ and $s$ when the considered domain (time/Laplace) is clear. Then it is straightforward to show that the input-output relationship in the Laplace domain reduces to a multiplication operation $y(s) = M(s)u(s)$, where $M(s) := \frac{\omega_c}{s+\omega_c}$ is called the transfer function of the block. Hence, for linear dynamical systems, the output of a block in the Laplace domain is simply the product between the block's transfer function and its input. This example block operates as a low pass filter that filters out high frequencies, particularly those higher than the cutoff frequency $\omega_c$ [1, Figure 8.15]. Note that in the limit, as $\omega_c \to \infty$, this block becomes the identity operator: $y = u$.

Panel c shows other commonly used blocks representing four linear dynamical systems: (1) a summation junction that sums (and/or subtracts) its inputs, (2) an integral block which integrates the input in time and is equivalent to dividing by $s$ in the Laplace domain, (3) a proportional block which multiplies its input by a constant, and (4) a derivative block which differentiates its input in time and is equivalent to multiplying by $s$ in the Laplace domain. The transfer functions of the integral, proportional and derivative blocks are thus $K_I/s, K_P$ and $K_D s$, respectively, as depicted in Panel c.

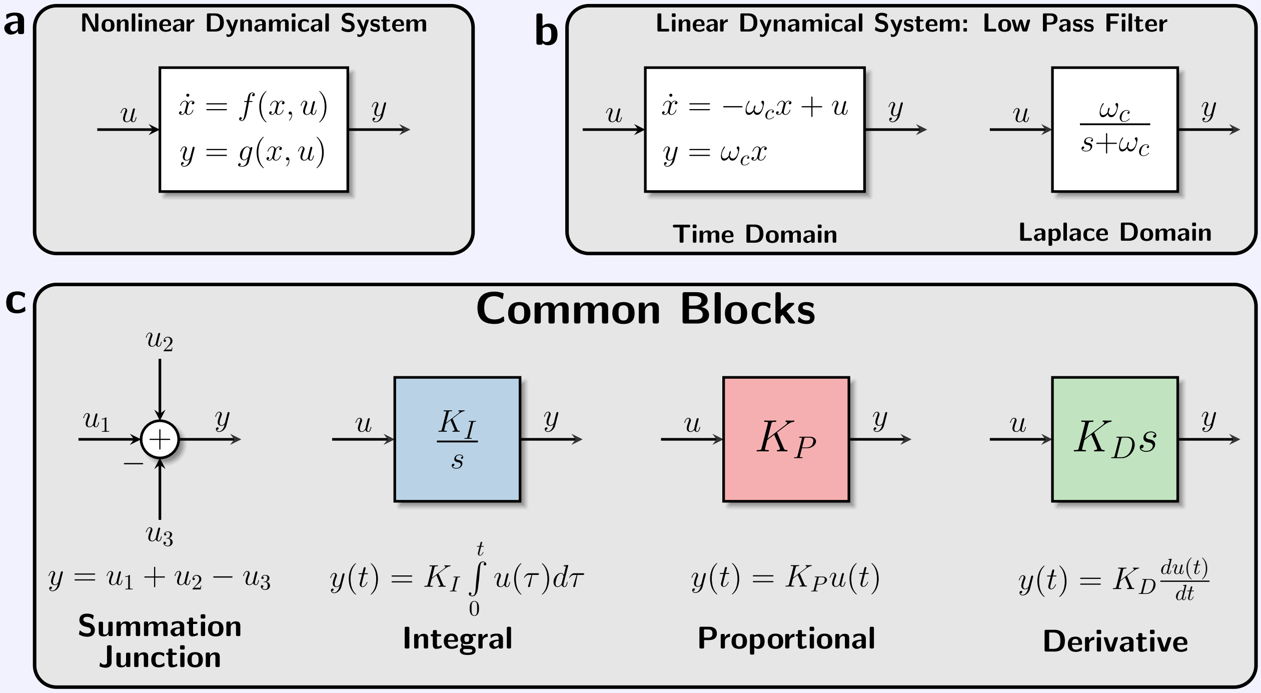

This is the controller transfer function that relates the control action $\tilde{u}$ to the reference signal $\tilde{\mu}$, the error signal $\tilde{e}$ and the plant output $\tilde{x}_L$ in the Laplace domain. Therefore, equipped with the controller and plant transfer functions given by

$$\textbf{Controller:} \quad \tilde{u}(s) = \left[K_F\tilde{\mu}(s) + \frac{K_I}{s}\tilde{e}(s) - K_{P_2}\tilde{x}_L(s)\right]\frac{\omega_c}{s+\omega_c} - K_{P_1}\tilde{x}_L(s); \qquad \tilde{e}(s) := \tilde{\mu}(s) - \left(\frac{\theta+\sigma_6}{\sigma_5}\right)\tilde{x}_L(s);$$

$$\textbf{Plant:} \quad \tilde{x}_L(s) = P(s)\tilde{u}(s);$$

$$\text{where:} \quad \begin{cases} K_F = \dfrac{\sigma_1\sigma_5}{\eta(\bar{z}_1+\bar{z}_2)}, & K_I = \sigma_5\dfrac{\sigma_1\bar{z}_1+\sigma_2\bar{z}_2}{\bar{z}_1+\bar{z}_2}, \quad K_S = \dfrac{\theta+\sigma_6}{\sigma_5}, \quad K_{P_1} = \sigma_4, \\ K_{P_2} = \dfrac{\sigma_2\theta+\sigma_1\sigma_6}{\eta(\bar{z}_1+\bar{z}_2)}, & \omega_c = \eta(\bar{z}_1+\bar{z}_2), \quad P(s) = e_L^T\left(sI-\bar{A}\right)^{-1}e_1, \end{cases} \tag{S2}$$

we can now draw the block diagram shown in Figure S1 which compactly encompasses all of the proposed $a$PI architectures. In particular, for the standalone $a$I controller, both proportional gains $K_{P_1}$ and $K_{P_2}$ are set to zero. For the $a$PI controller of Class 1 (resp. Class 2 & 3), the proportional gain $K_{P_2}$ (resp. $K_{P_1}$) is set to zero. The remaining gains $K_I, K_F$, and $K_S$ are obtained by calculating the partial derivatives $\sigma_i$s for the various $a$PI controllers (see Section S3), and the results are shown in the table of Figure S1. Observe that, for all the proposed architectures, there is an Integral (I) and a Proportional (P) control action. In fact, since the controller acts on both the error signal $\tilde{e}$ and the output signal $\tilde{x}_L$, then the PI architecture (of the linearized dynamics) resembles the setting given in Figure 1(d). The main differences are two additional blocks:

- **Feedforward Block:** This block is a consequence of the positivity of the nonlinear dynamics. The reference signal $\tilde{\mu}$ "lifts" the dynamics towards the positive orthant, by adding the feedforward term to the integrated error.
- **Low Pass Filter:** This block is a dynamical system that filters fast signals with frequencies higher than the cutoff frequency $\omega_c = \eta(\bar{z}_1+\bar{z}_2)$. This block is a consequence of the time dynamics of the nonlinear sequestration reaction.

As demonstrated in Figure S1, for the $a$PI controllers of Class 1 ($K_{P_1} \neq 0, K_{P_2} = 0$), the Proportional control action $K_{P_1}\tilde{x}_L$ is instantaneous since it is fed back to the plant as is and without any filtering (that involves time dynamics). In contrast, for the $a$PI controllers of Class 2 and 3 ($K_{P_1} = 0, K_{P_2} \neq 0$), the Proportional control action $K_{P_2}\tilde{x}_L$ is not instantaneous since it is passed through a low pass filter before it is fed back to the plant. This low pass filtering step arises because the output species does not actuate the input species immediately like the $a$PI controllers of Class 1; instead, the output actuates the input via an intermediate controller species: $\mathbf{Z_2}$ (for Class 2) and $\mathbf{Z_1}$ (for Class 3). This low pass filter typically delays the Proportional control action, and as a result – depending on the performance objective and particular plant at hand – it can have a negative or positive effect on the performance of the closed-loop dynamics.

Note that, if the sequestration reaction is fast enough ($\eta$ is large), the effects of the feedforward block and low pass filter become negligible. This can be observed by examining the asymptotic limit, as $\eta \to \infty$, that yields

$$\lim_{\eta\to\infty} K_F = 0 \qquad \text{and} \qquad \lim_{\eta\to\infty}\left|\frac{\omega_c}{s+\omega_c}\right| = 1.$$

Consequently, as $\eta \to \infty$, the PI architecture of the linearized dynamics becomes exactly the same as that given in Figure 1(d). Lastly, observe using the table of Figure S1 that the various PI gains may depend mutually on the same biomolecular controller parameters. As an example, for Class 1 with multiplicative inhibition, the biomolecular controller parameter $\kappa$ can tune both the proportional gain $K_{P_1}$ and the integral gain $K_I$ simultaneously. This is a consequence of the inseparability of the original nonlinear PI architecture. In contrast, for Class 1 with additive inhibition, the biomolecular controller parameter $\alpha$ can tune the proportional gain $K_{P_1}$ only. It is shown, that this simultaneous tuning of the PID gains, with a single controller parameter, yields better stability properties and performance (see Figure 4 in the main text).

#### S1.2 Antithetic Proportional-Integral Derivative ($a$PID) Feedback Controllers

Once again, to analytically verify the various PID architectures, we carry out a linear perturbation analysis similar to that carried out for the $a$PI controllers. The analysis is carried out for each PID controller (with different orders) separately.

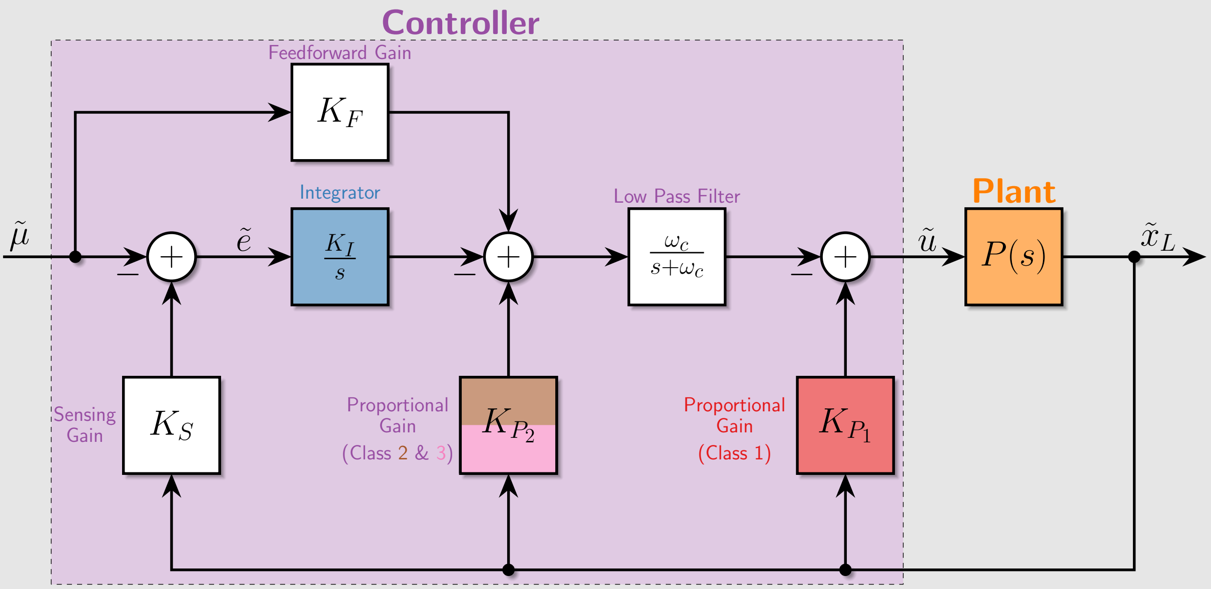

| Class | Inhibition | $K_I$ | $K_S$ | $K_{P_1}$ | $K_{P_2}$ | $K_F$ |
|---|---|---|---|---|---|---|
| **AIF** | None | $k\frac{\bar{z}_1}{\bar{z}_1+\bar{z}_2}$ | $\theta$ | 0 | 0 | $\frac{k}{\eta(\bar{z}_1+\bar{z}_2)}$ |
| **APIF Class 1** | Additive | $k\frac{\bar{z}_1}{\bar{z}_1+\bar{z}_2}$ | $\theta$ | $\frac{\alpha}{\kappa}\frac{n(\bar{x}_L/\kappa)^{n-1}}{[1+(\bar{x}_L/\kappa)^n]^2}$ | 0 | $\frac{k}{\eta(\bar{z}_1+\bar{z}_2)}$ |
| | Multiplicative | $\frac{k}{1+(\bar{x}_L/\kappa)^n}\frac{\bar{z}_1}{\bar{z}_1+\bar{z}_2}$ | $\theta$ | $\frac{k\bar{z}_1}{\kappa}\frac{n(\bar{x}_L/\kappa)^{n-1}}{[1+(\bar{x}_L/\kappa)^n]^2}$ | 0 | $\frac{1}{1+(\bar{x}_L/\kappa)^n}\frac{k}{\eta(\bar{z}_1+\bar{z}_2)}$ |
| | Degradation | $k\frac{\bar{z}_1}{\bar{z}_1+\bar{z}_2}$ | $\theta$ | $\delta\frac{\bar{x}_1}{\bar{x}_1+\kappa_1}$ | 0 | $\frac{k}{\eta(\bar{z}_1+\bar{z}_2)}$ |
| **APIF Class 2** | Additive | $\left(k\bar{z}_1+\alpha\frac{(\bar{z}_2/\kappa)^n}{(1+(\bar{z}_2/\kappa)^n)^2}\right)\frac{1}{\bar{z}_1+\bar{z}_2}$ | $\theta$ | 0 | $\frac{\alpha\theta}{\kappa\eta(\bar{z}_1+\bar{z}_2)}\frac{n(\bar{z}_2/\kappa)^{n-1}}{[1+(\bar{z}_2/\kappa)^n]^2}$ | $\frac{k}{\eta(\bar{z}_1+\bar{z}_2)}$ |
| | Multiplicative | $k\frac{\bar{z}_1}{\bar{z}_1+\bar{z}_2}\frac{1+(n+1)(\bar{z}_2/\kappa)^n}{[1+(\bar{z}_2/\kappa)^n]^2}$ | $\theta$ | 0 | $k\frac{\bar{z}_1}{\bar{z}_1+\bar{z}_2}\frac{\theta}{\kappa\eta}\frac{n(\bar{z}_2/\kappa)^{n-1}}{[1+(\bar{z}_2/\kappa)^n]^2}$ | $\frac{1}{1+(\bar{z}_2/\kappa)^n}\frac{k}{\eta(\bar{z}_1+\bar{z}_2)}$ |
| | Degradation | $\frac{k\bar{z}_1+\delta\bar{x}_1\bar{z}_2}{\bar{z}_1+\bar{z}_2}$ | $\theta$ | 0 | $\frac{\delta\theta}{\eta(\bar{z}_1+\bar{z}_2)}\frac{\bar{x}_1}{\bar{x}_1+\kappa_1}$ | $\frac{k}{\eta(\bar{z}_1+\bar{z}_2)}$ |
| **APIF Class 3** | Additive | $k\frac{\bar{z}_1}{\bar{z}_1+\bar{z}_2}$ | $\theta+\frac{\alpha}{\kappa}\frac{n(\bar{x}_L/\kappa)^{n-1}}{[1+(\bar{x}_L/\kappa)^n]^2}$ | 0 | $\frac{k\alpha}{\kappa\eta(\bar{z}_1+\bar{z}_2)}\frac{n(\bar{x}_L/\kappa)^{n-1}}{[1+(\bar{x}_L/\kappa)^n]^2}$ | $\frac{k}{\eta(\bar{z}_1+\bar{z}_2)}$ |
| | Multiplicative | $\frac{k}{1+(\bar{x}_L/\kappa)^n}\frac{\bar{z}_1}{\bar{z}_1+\bar{z}_2}$ | $\theta\left[1+(\bar{x}_L/\kappa)^n\right]+\bar{\mu}/\kappa\frac{n(\bar{x}_L/\kappa)^{n-1}}{[1+(\bar{x}_L/\kappa)^n]^2}$ | 0 | $\frac{k\bar{\mu}}{\kappa\eta(\bar{z}_1+\bar{z}_2)}\frac{n(\bar{x}_L/\kappa)^{n-1}}{[1+(\bar{x}_L/\kappa)^n]^2}$ | $\frac{1}{1+(\bar{x}_L/\kappa)^n}\frac{k}{\eta(\bar{z}_1+\bar{z}_2)}$ |

Figure S1: **Block Diagram of the $a$PI Feedback Controllers.** This block diagram compactly represents the dynamics (in the Laplace domain) of the linearized closed-loop systems obtained by the various $a$PI controllers proposed in Figure 3 . Particularly, $a$PI controllers of Class 1 (resp. 2 and 3) give rise to the proportional gain $K_{P_1}$ (resp. $K_{P_2}$). The low pass filter in between demonstrates the instantaneous (resp. filtered) proportional control action of the $a$PI controllers of Class 1 (resp. 2 and 3). The table shows the PI gains as a function of the various biomolecular parameters.

##### S1.2.1 Simple Second-Order $a$PID Feedback Controller

Consider an arbitrary network controlled by the second order $a$PID controller depicted in Figure S2(a). The deterministic dynamics of the closed-loop system are given by

$$\begin{cases} \dot{x} = f(x) + h(z_1, x_1, x_L)e_1 \\ \dot{z}_1 = \mu + \beta x_L - \eta z_1 z_2 \\ \dot{z}_2 = \theta x_L - \eta z_1 z_2, \end{cases} \tag{S3}$$

where $f(x) = S\lambda(x)$. Similar to the $a$PI controllers, the actuation propensity $h$ can take different forms as illustrated in the table of Figure S2(a). Once again, let $\begin{bmatrix}\tilde{x}^T & \tilde{z}_1 & \tilde{z}_2\end{bmatrix}^T$ denote the perturbation from the fixed point $\begin{bmatrix}\bar{x}^T & \bar{z}_1 & \bar{z}_2\end{bmatrix}^T$ of (S3). We also assume that the reference signal $\mu$ is allowed to slightly vary in time around a nominal reference $\bar{\mu}$. That is, we have

$$\tilde{x}(t) = x(t) - \bar{x}; \qquad \tilde{z}_1(t) = z_1(t) - \bar{z}_1; \qquad \tilde{z}_2(t) = z_2(t) - \bar{z}_2; \qquad \tilde{\mu}(t) = \mu(t) - \bar{\mu}.$$

**a**

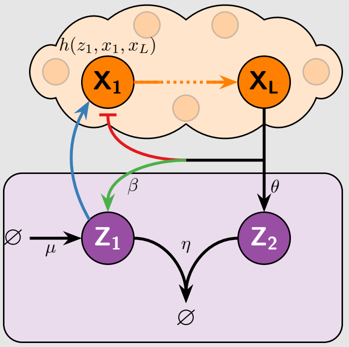

**Total Actuation Propensity**

$$h(z_1, x_1, x_L) := h^+(z_1, x_L) - h^-(x_L)\frac{x_1}{x_1+\kappa_1}$$

$$S_{cl} = \begin{array}{c} \overbrace{\mathcal{R}^1 \ \cdots \ \mathcal{R}^K}^{\mathcal{R}} \ \overbrace{\mathcal{R}_s \ \mathcal{R}_q \ \mathcal{R}_r \ \mathcal{R}_a^+ \ \mathcal{R}_a^-}^{\mathcal{R}_c} \\ \left[\begin{array}{c|ccccc} & 0 & 0 & 0 & 1 & -1 \\ S & & & & & \\ & 0 & 0 & 0 & 0 & 0 \\ \hline 0 & 0 & -1 & 1 & 0 & 0 \\ & 1 & -1 & 0 & 0 & 0 \end{array}\right] \end{array} \begin{array}{l} \left.\begin{array}{c} X_1 \\ \vdots \\ X_L \end{array}\right\} X \\ \left.\begin{array}{c} Z_1 \\ Z_2 \end{array}\right\} Z \end{array}$$

$$\lambda_{cl}(x, z) = \left[\begin{array}{c|ccccc} \overbrace{\mathcal{R}^1 \ \cdots \ \mathcal{R}^K}^{\mathcal{R}} & \mathcal{R}_s & \mathcal{R}_q & \mathcal{R}_r & \mathcal{R}_a^+ & \mathcal{R}_a^- \\ \lambda(x) & \theta x_L & \eta z_1 z_2 & \mu + \beta x_L & h^+(z_1, x_L) & h^-(x_L)\frac{x_1}{x_1+\kappa_1} \end{array}\right]^T$$

| Reference Rxn: $\mathcal{R}_r$ | Sensing Rxn: $\mathcal{R}_s$ | Sequestration Rxn: $\mathcal{R}_q$ |
|---|---|---|
| $\varnothing \xrightarrow{\mu + \beta x_L} Z_1$ | $X_L \xrightarrow{\theta} X_L + Z_2$ | $Z_1 + Z_2 \xrightarrow{\eta} \varnothing$ |

| Inhibition Type | Actuation Rxns | Total Actuation Propensity $h$ |
|---|---|---|
| Additive | $\mathcal{R}_a^+ : \ \varnothing \longrightarrow X_1$ | $kz_1 + \frac{\alpha}{1+(x_L/\kappa)^n}$ |
| Multiplicative | $\mathcal{R}_a^+ : \ \varnothing \longrightarrow X_1$ | $\frac{kz_1}{1+(x_L/\kappa)^n}$ |
| Degradation | $\mathcal{R}_a^+ : \ Z_1 \longrightarrow Z_1 + X_1$<br>$\mathcal{R}_a^- : \ X_1 + X_L \longrightarrow X_L$ | $kz_1 - \delta x_L \frac{x_1}{x_1+\kappa_1}$ |

**b**

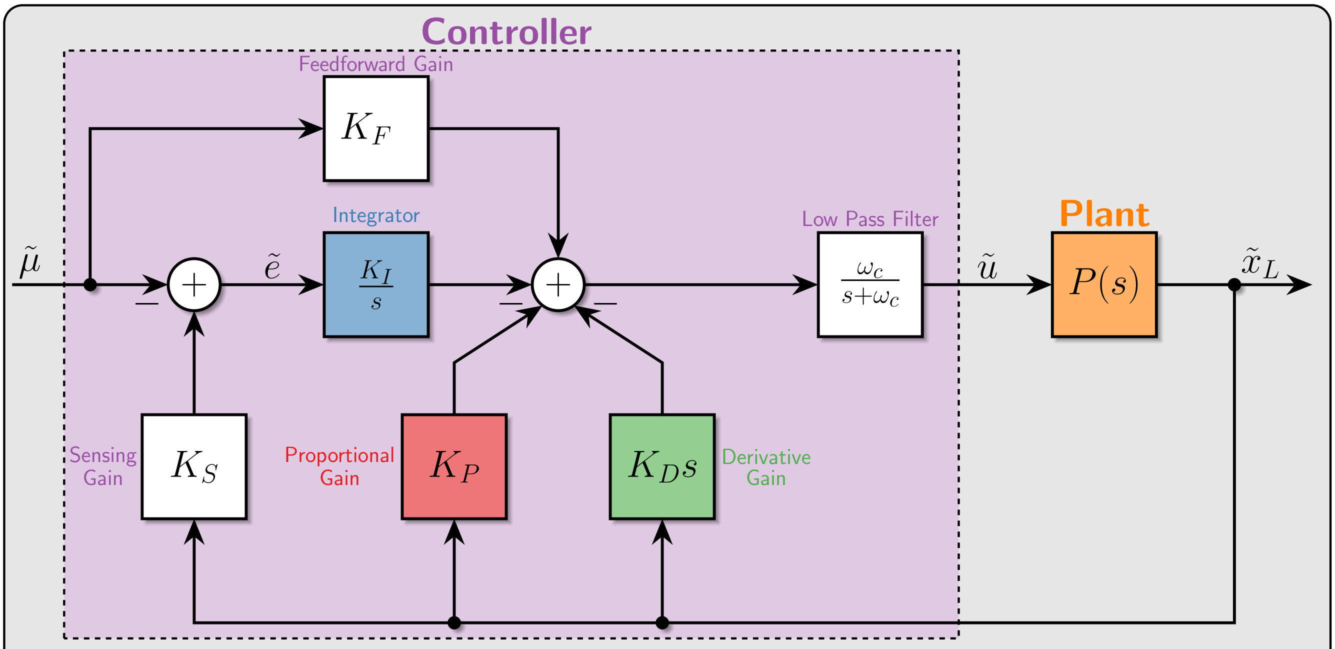

| Inhibition | $K_I$ | $K_P$ | $K_D$ | $K_F$ |
|---|---|---|---|---|
| Additive | $k\frac{\bar{z}_1}{\bar{z}_1+\bar{z}_2}$ | $\frac{\alpha}{\kappa}\frac{n(\bar{x}_L/\kappa)^{n-1}}{[1+(\bar{x}_L/\kappa)^n]^2} - \beta\frac{k}{\eta(\bar{z}_1+\bar{z}_2)}$ | $\frac{\alpha/\kappa}{\eta(\bar{z}_1+\bar{z}_2)}\frac{n(\bar{x}_L/\kappa)^{n-1}}{[1+(\bar{x}_L/\kappa)^n]^2}$ | $\frac{k}{\eta(\bar{z}_1+\bar{z}_2)}$ |
| Multiplicative | $\frac{k}{1+(\bar{x}_L/\kappa)^n}\frac{\bar{z}_1}{\bar{z}_1+\bar{z}_2}$ | $\frac{k}{1+(\bar{x}_L/\kappa)^n}\left[\frac{\bar{z}_1}{\kappa}\frac{n(\bar{x}_L/\kappa)^{n-1}}{1+(\bar{x}_L/\kappa)^n} - \frac{\beta}{\eta(\bar{z}_1+\bar{z}_2)}\right]$ | $\frac{k\bar{z}_1/\kappa}{\eta(\bar{z}_1+\bar{z}_2)}\frac{n(\bar{x}_L/\kappa)^{n-1}}{[1+(\bar{x}_L/\kappa)^n]^2}$ | $\frac{1}{1+(\bar{x}_L/\kappa)^n}\frac{k}{\eta(\bar{z}_1+\bar{z}_2)}$ |
| Degradation | $k\frac{\bar{z}_1}{\bar{z}_1+\bar{z}_2}$ | $\delta\frac{\bar{x}_1}{\bar{x}_1+\kappa} - \frac{\beta k}{\eta(\bar{z}_1+\bar{z}_2)}$ | $\frac{\delta}{\eta(\bar{z}_1+\bar{z}_2)}\frac{\bar{x}_1}{\bar{x}_1+\kappa}$ | $\frac{k}{\eta(\bar{z}_1+\bar{z}_2)}$ |

Figure S2: **Second-Order *a*PID Feedback Controllers. (a) *a*PID Controller Network.** An additional production reaction (marked in green) is appended to the *a*PI controller of class 1 (see Figure 3) to embed an additional derivative control action. Intuitively, this additional reaction introduces an incoherent feedforward pathway from $\mathbf{X_L}$ to $\mathbf{X_1}$ that is cascaded with the standalone *a*I controller. Similar to the previously introduced *a*PI controllers, three different biologically-relevant inhibition mechanisms are considered. **(b) *a*PID Controller Block Diagram.** Similar to the block diagram of the *a*PI controllers depicted in Figure S1, this block diagram represents the dynamics of the linearized closed-loop system obtained by the second-order *a*PID controller with all three inhibition mechanisms. The table shows the PID gains as a function of the various biomolecular parameters.

The linearized dynamics can thus be written as

$$\begin{cases} \dot{\tilde{x}} = A\tilde{x} + (\sigma_1\tilde{z}_1 - \sigma_3\tilde{x}_1 - \sigma_4\tilde{x}_L)\, e_1 \\ \dot{\tilde{z}}_1 = \tilde{\mu} + \beta\tilde{x}_L - \eta\bar{z}_2\tilde{z}_1 - \eta\bar{z}_1\tilde{z}_2 \\ \dot{\tilde{z}}_2 = \theta\tilde{x}_L - \eta\bar{z}_2\tilde{z}_1 - \eta\bar{z}_1\tilde{z}_2, \end{cases}$$

where $A := \partial f(\bar{x})$, $\partial h(\bar{z}_1, \bar{x}_1, \bar{x}_L) =: \begin{bmatrix}\sigma_1 & -\sigma_3 & -\sigma_4\end{bmatrix}$ such that $\sigma_1 > 0$ and $\sigma_3, \sigma_4 \geq 0$. Taking the Laplace transforms yields

$$\begin{aligned} \tilde{x}_L(s) &= e_L^T \left(sI - \bar{A}\right)^{-1} e_1\tilde{u}(s); \qquad \text{where} \quad \bar{A} := A - \sigma_3 e_1 e_1^T \\ \tilde{u}(s) &= \sigma_1\tilde{z}_1(s) - \sigma_4\tilde{x}_L(s) \\ \tilde{z}_1(s) &= \frac{\tilde{\mu}(s) + \beta\tilde{x}_L(s) - \eta\bar{z}_1\tilde{z}_2(s)}{s + \eta\bar{z}_2} \\ \tilde{z}_2(s) &= \frac{\theta\tilde{x}_L(s) - \eta\bar{z}_2\tilde{z}_1(s)}{s + \eta\bar{z}_1}. \end{aligned}$$

Next, we express $\tilde{z}_1(s)$ and $\tilde{z}_2(s)$ in terms of $\tilde{\mu}(s), \tilde{x}_L(s)$ and the error $\tilde{e}(s) := \tilde{\mu}(s) - (\theta - \beta)\,\tilde{x}_L(s)$ as

$$\begin{aligned} \tilde{z}_1(s) &= \left[\tilde{\mu}(s) + \beta\tilde{x}_L(s) + \frac{\eta\bar{z}_1}{s}\tilde{e}(s)\right] \frac{1}{s + \eta\left(\bar{z}_1 + \bar{z}_2\right)} \\ \tilde{z}_2(s) &= \left[\theta\tilde{x}_L(s) - \frac{\eta\bar{z}_2}{s}\tilde{e}(s)\right] \frac{1}{s + \eta\left(\bar{z}_1 + \bar{z}_2\right)}. \end{aligned}$$

The feedback control action $\tilde{u}(s)$ can thus be written as

$$\begin{aligned} \tilde{u}(s) &= \left[\sigma_1\tilde{\mu}(s) + \frac{\eta\sigma_1\bar{z}_1}{s}\tilde{e}(s) + \sigma_1\beta\tilde{x}_L(s)\right] \frac{1}{s + \eta\left(\bar{z}_1 + \bar{z}_2\right)} - \sigma_4\tilde{x}_L(s) \\ &= \left[\sigma_1\tilde{\mu}(s) + \frac{\eta\sigma_1\bar{z}_1}{s}\tilde{e}(s) - \sigma_4 s\tilde{x}_L(s) - \left(\sigma_4\eta(\bar{z}_1 + \bar{z}_2) - \sigma_1\beta\right)\tilde{x}_L(s)\right] \frac{1}{s + \eta\left(\bar{z}_1 + \bar{z}_2\right)}. \end{aligned}$$

This is the controller transfer function that relates the control action $\tilde{u}$ to the reference signal $\tilde{\mu}$, the error signal $\tilde{e}$ and the plant output $\tilde{x}_L$ in the Laplace domain. Therefore, equipped with the controller and plant transfer functions given by

$$\begin{aligned} &\textbf{Controller:} \quad \tilde{u}(s) = \left[K_F\tilde{\mu}(s) + \frac{K_I}{s}\tilde{e}(s) - \Big(K_P + K_D s\Big)\tilde{x}_L(s)\right] \frac{\omega_c}{s + \omega_c}; \qquad \tilde{e}(s) := \tilde{\mu}(s) - (\theta - \beta)\,\tilde{x}_L(s); \\ &\textbf{Plant:} \quad \tilde{x}_L(s) = P(s)\tilde{u}(s); \\ &\text{where:} \begin{cases} K_F = \dfrac{\sigma_1}{\eta(\bar{z}_1 + \bar{z}_2)}, \quad K_I = \sigma_1\dfrac{\bar{z}_1}{\bar{z}_1 + \bar{z}_2}, \quad K_S = \theta - \beta, \quad K_P = \sigma_4 - \dfrac{\sigma_1\beta}{\eta(\bar{z}_1 + \bar{z}_2)}, \\ K_D = \dfrac{\sigma_4}{\eta(\bar{z}_1 + \bar{z}_2)} \quad \omega_c = \eta(\bar{z}_1 + \bar{z}_2), \quad P(s) = e_L^T\left(sI - \bar{A}\right)^{-1} e_1, \end{cases} \end{aligned} \tag{S4}$$

we can now draw the block diagram shown in Figure S2(b) which compactly encompasses the proposed second-order $a$PID architectures with three different inhibition mechanisms. Observe that the various biomolecular controller parameters $(\eta, \theta, \beta, ...)$ appear mutually in the various PID gains and cutoff frequency. This is the consequence of having an inseparable PID controller. Recall that for the $a$PI controllers proposed and analyzed in Section S1.1, the sequestration reaction is assumed to be strong ($\eta$ is large). In contrast, for the proposed second-order $a$PID here, $\eta$ cannot be large because the derivative gain $K_D := \frac{\sigma_4}{\eta(\bar{z}_1 + \bar{z}_2)}$ becomes negligible. As a result, to obtain a complete PID architecture, $\eta$ should play the role of a tuning parameter to control the derivative gain $K_D$. As a result, the obtained control architecture is a filtered PID.

##### S1.2.2 Third-Order $a$PID Feedback Controller

Consider an arbitrary network controlled by the third order $a$PID controller depicted in Figure S3(a). The deterministic dynamics of the closed-loop system are given by

$$\begin{cases} \dot{x} = f(x) + h(z_1, z_3, x_1, x_L)e_1 \\ \dot{z}_1 = \mu - \eta z_1 z_2 \\ \dot{z}_2 = \theta x_L - \eta z_1 z_2 \\ \dot{z}_3 = g(z_3, x_L) - \gamma_0 z_3, \end{cases} \tag{S5}$$

where $f(x) = S\lambda(x)$. The function $h$ is chosen to be monotonically increasing (resp. decreasing) in $z_1$ (resp. in $x_1$ and $x_L$), $g$ is monotonically decreasing in $z_3$, while $g$ and $h$ can be chosen to have any monotonicity in $x_L$ and $z_3$ (see the diamond arrowhead in Figure S3(a)).

Let $\begin{bmatrix}\tilde{x}^T & \tilde{z}_1 & \tilde{z}_2 & \tilde{z}_3\end{bmatrix}^T$ denote the perturbation from the fixed point $\begin{bmatrix}\bar{x}^T & \bar{z}_1 & \bar{z}_2 & \bar{z}_3\end{bmatrix}^T$ of (S5). We also assume that the reference signal $\mu$ is allowed to slightly vary in time around a nominal reference $\bar{\mu}$. That is, we have

$$\tilde{x}(t) = x(t) - \bar{x}; \qquad \tilde{\mu}(t) = \mu(t) - \bar{\mu}; \qquad \tilde{z}_i(t) = z_i(t) - \bar{z}_i; \quad (i = 1, 2, 3).$$

The linearized dynamics can thus be written as

$$\begin{cases} \dot{\tilde{x}} = A\tilde{x} + (\sigma_1\tilde{z}_1 + \sigma_2\tilde{z}_3 - \sigma_3\tilde{x}_1 - \sigma_4\tilde{x}_L)\, e_1 \\ \dot{\tilde{z}}_1 = \tilde{\mu} - \eta\bar{z}_2\tilde{z}_1 - \eta\bar{z}_1\tilde{z}_2 \\ \dot{\tilde{z}}_2 = \theta\tilde{x}_L - \eta\bar{z}_2\tilde{z}_1 - \eta\bar{z}_1\tilde{z}_2 \\ \dot{\tilde{z}}_3 = \sigma_6\tilde{x}_L - (\gamma + \sigma_5)z_3. \end{cases}$$

where $A := \partial f(\bar{x})$, $\partial h(\bar{z}_1, \bar{z}_3, \bar{x}_1, \bar{x}_L) =: \begin{bmatrix}\sigma_1 & \sigma_2 & -\sigma_3 & -\sigma_4\end{bmatrix}$ and $\partial g(\bar{z}_3, \bar{x}_L) =: \begin{bmatrix}-\sigma_5 & \sigma_6\end{bmatrix}$ such that $\sigma_1, \sigma_4 > 0$, $\sigma_3$ and $\sigma_5 \geq 0$. Taking the Laplace transforms yields

$$\begin{aligned} \tilde{x}_L(s) &= e_L^T\left(sI - \bar{A}\right)^{-1} e_1\tilde{u}(s); \qquad \text{where} \quad \bar{A} := A - \sigma_3 e_1 e_1^T \\ \tilde{u}(s) &= \sigma_1\tilde{z}_1(s) + \sigma_2\tilde{z}_3(s) - \sigma_4\tilde{x}_L(s) \\ \tilde{z}_1(s) &= \frac{\tilde{\mu}(s) - \eta\bar{z}_1\tilde{z}_2(s)}{s + \eta\bar{z}_2} \\ \tilde{z}_2(s) &= \frac{\theta\tilde{x}_L(s) - \eta\bar{z}_2\tilde{z}_1(s)}{s + \eta\bar{z}_1} \\ \tilde{z}_3(s) &= \frac{\sigma_6\tilde{x}_L(s)}{s + \gamma_0 + \sigma_5}. \end{aligned}$$

Next, we express $\tilde{z}_1(s)$ and $\tilde{z}_2(s)$ in terms of $\tilde{\mu}(s), \tilde{x}_L(s)$ and the error $\tilde{e}(s) := \tilde{\mu}(s) - \theta\tilde{x}_L(s)$ as

$$\begin{aligned} \tilde{z}_1(s) &= \left[\tilde{\mu}(s) + \frac{\eta\bar{z}_1}{s}\tilde{e}(s)\right]\frac{1}{s + \eta\left(\bar{z}_1 + \bar{z}_2\right)} \\ \tilde{z}_2(s) &= \left[\theta\tilde{x}_L(s) - \frac{\eta\bar{z}_2}{s}\tilde{e}(s)\right]\frac{1}{s + \eta\left(\bar{z}_1 + \bar{z}_2\right)}. \end{aligned}$$

The feedback control action $\tilde{u}(s)$ can thus be written as

$$\begin{aligned} \tilde{u}(s) &= \left[\sigma_1\tilde{\mu}(s) + \frac{\eta\sigma_1\bar{z}_1}{s}\tilde{e}(s)\right]\frac{1}{s + \eta\left(\bar{z}_1 + \bar{z}_2\right)} + \left[\frac{\sigma_2\sigma_6}{s + \gamma_0 + \sigma_5} - \sigma_4\right]\tilde{x}_L(s) \\ &= \left[\sigma_1\tilde{\mu}(s) + \frac{\eta\sigma_1\bar{z}_1}{s}\tilde{e}(s)\right]\frac{1}{s + \eta\left(\bar{z}_1 + \bar{z}_2\right)} - \left[\sigma_4 s + \sigma_4(\gamma_0 + \sigma_5) - \sigma_2\sigma_6\right]\frac{\tilde{x}_L(s)}{s + \gamma_0 + \sigma_5}. \end{aligned}$$

This is the controller transfer function that relates the control action $\tilde{u}$ to the reference signal $\tilde{\mu}$, the error signal $\tilde{e}$ and the plant output $\tilde{x}_L$ in the Laplace domain. Therefore, equipped by the controller and plant transfer functions given by

$$\begin{aligned} &\textbf{Controller:} \quad \tilde{u}(s) = \left[K_F\tilde{\mu}(s) + \frac{K_I}{s}\tilde{e}(s)\right]\frac{\omega_c}{s + \omega_c} - \left[K_P + K_D s\right]\frac{\omega_0}{s + \omega_0}\tilde{x}_L(s); \qquad \tilde{e}(s) := \tilde{\mu}(s) - (\theta - \beta)\,\tilde{x}_L(s) \\ &\textbf{Plant:} \quad \tilde{x}_L(s) = P(s)\tilde{u}(s); \\ &\text{where:} \quad \begin{cases} K_F = \dfrac{\sigma_1}{\eta(\bar{z}_1 + \bar{z}_2)}, \quad K_I = \sigma_1\dfrac{\bar{z}_1}{\bar{z}_1 + \bar{z}_2}, \quad K_S = \theta, \quad K_P = \sigma_4 - \dfrac{\sigma_2\sigma_6}{\gamma_0 + \sigma_5}, \\ K_D = \dfrac{\sigma_4}{\gamma_0 + \sigma_5} \quad \omega_c = \eta(\bar{z}_1 + \bar{z}_2), \quad \omega_0 = \gamma_0 + \sigma_5, \quad P(s) = e_L^T\left(sI - \bar{A}\right)^{-1} e_1, \end{cases} \end{aligned} \tag{S6}$$

we can now draw the block diagram shown in Figure S3(b) which encompasses the proposed third-order $a$PID architectures with the different inhibition mechanisms. The first term in the controller transfer function corresponds to the filtered integral action that simplifies to a pure integral action $\frac{K_I}{s}\tilde{e}(s)$ as $\eta \to \infty$. The second term involves a filtered PD control action with a cutoff frequency $\omega_0$ as demonstrated in the block diagram. Note that, if the cutoff frequency is large enough particularly $\omega_0 > K_P/K_D$ which is always satisfied when $\mathbf{Z_3}$ embeds an incoherent feed-forward loop (i.e. $\sigma_2\sigma_6 > 0$), then a lag compensator is obtained. Otherwise, if $\omega_0 < K_P/K_D$ which is always satisfied when $\mathbf{Z_3}$ embeds a coherent feed-forward loop (i.e. $\sigma_2\sigma_6 < 0$), then a lead compensator is obtained. In conclusion, the overall realized control architecture (as $\eta \to \infty$) is I + filtered PD.

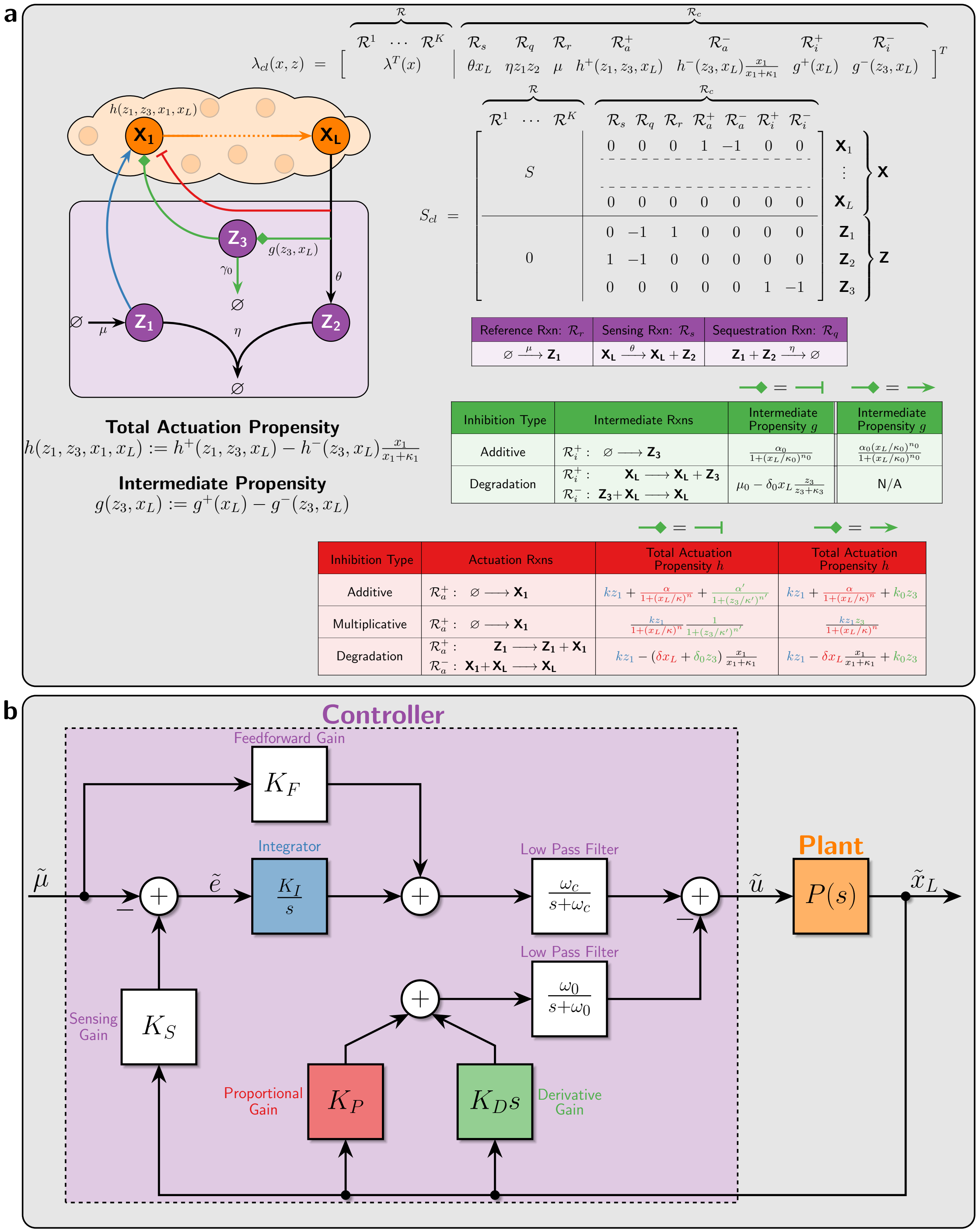

Figure S3: **Third-Order *a*PID Feedback Controllers. (a) *a*PID Controller Network.** An additional pathway (marked in green) is appended to the *a*PI controller of class 1 (see Figure 3) via an intermediate controller species $\mathbf{Z_3}$ to embed an additional derivative control action. Similar to the second-order *a*PID, this additional pathway introduces an incoherent feedforward loop from $\mathbf{X_L}$ to $\mathbf{X_1}$ via the intermediate species $\mathbf{Z_3}$. Note that this can be done by either two inhibitions or two activations as depicted by the green diamond arrowhead. **(b) *a*PID Controller Block Diagram.** This block diagram represents the dynamics of the linearized closed-loop system obtained by the third-order *a*PID controller with different inhibition mechanisms. An additional low pass filter appears (with cutoff frequency $\omega_0$) which indicates that the realized PD control action is filtered before it actuates the plant. The various gains are expressed in terms of the biomolecular parameters and partial derivatives of $h$ and $g$ in (S6).

##### S1.2.3 Fourth-Order *a*PID Feedback Controller (with Additive D)

Consider an arbitrary network controlled by the fourth-order PID controller depicted in Figure S4(a). The deterministic dynamics of the closed-loop system are given by

$$\begin{cases} \dot{x} = f(x) + h(z_1, x_1, x_L, u_D)e_1; \qquad u_D = g(z_3, x_L) \\ \dot{z}_1 = \mu - \eta z_1 z_2 \\ \dot{z}_2 = \theta x_L - \eta z_1 z_2 \\ \dot{z}_3 = \mu_0 - \eta_0 z_3 z_4 \\ \dot{z}_4 = \theta_0 u_D - \eta_0 z_3 z_4, \end{cases} \tag{S7}$$

where $f(x) = S\lambda(x)$. The function $h$ is chosen to be monotonically increasing (resp. decreasing) in $z_1$ and $u_D$ (resp. in $x_1$ and $x_L$), while $g$ is monotonically increasing (resp. decreasing) in $z_3$ (resp. $x_L$) as depicted in the network diagram of Figure S4(a). An example of the function $g$ is

$$u_D = g(z_3, x_L) = \frac{k_0 z_3}{1 + (x_L/\kappa_0)^{n_0}}.$$

Let $\begin{bmatrix}\tilde{x}^T & \tilde{z}_1 & \tilde{z}_2 & \tilde{z}_3 & \tilde{z}_4 & \tilde{u}_D\end{bmatrix}^T$ denote the perturbation from the fixed point $\begin{bmatrix}\bar{x}^T & \bar{z}_1 & \bar{z}_2 & \bar{z}_3 & \bar{z}_4 & \bar{u}_D\end{bmatrix}^T$ of (S7) with $\bar{u}_D := g(\bar{z}_3, \bar{x}_L) = \mu_0/\theta_0$. We also assume that the reference signal $\mu$ is allowed to slightly vary in time around a nominal reference $\bar{\mu}$. That is, we have

$$\tilde{u}_D(t) = u_D(t) - \bar{u}_D; \qquad \tilde{x}(t) = x(t) - \bar{x}; \qquad \tilde{\mu}(t) = \mu(t) - \bar{\mu}; \qquad \tilde{z}_i(t) = z_i(t) - \bar{z}_i; \quad (i = 1, 2, 3, 4).$$

The linearized dynamics can thus be written as

$$\begin{cases} \dot{\tilde{x}} = A\tilde{x} + (\sigma_1 \tilde{z}_1 - \sigma_3 \tilde{x}_1 - \sigma_4 \tilde{x}_L)\, e_1 + \sigma_D \tilde{u}_D e_1; \qquad \tilde{u}_D := \sigma_5 \tilde{z}_3 - \sigma_6 \tilde{x}_L \\ \dot{\tilde{z}}_1 = \tilde{\mu} - \eta \bar{z}_2 \tilde{z}_1 - \eta \bar{z}_1 \tilde{z}_2 \\ \dot{\tilde{z}}_2 = \theta \tilde{x}_L - \eta \bar{z}_2 \tilde{z}_1 - \eta \bar{z}_1 \tilde{z}_2 \\ \dot{\tilde{z}}_3 = -\eta_0 \bar{z}_4 \tilde{z}_3 - \eta_0 \bar{z}_3 \tilde{z}_4 \\ \dot{\tilde{z}}_4 = \theta_0 \tilde{u}_D - \eta_0 \bar{z}_4 \tilde{z}_3 - \eta_0 \bar{z}_3 \tilde{z}_4, \end{cases}$$

where $A := \partial f(\bar{x})$, $\partial h(\bar{z}_1, \bar{x}_1, \bar{x}_L, \bar{u}_D) =: \begin{bmatrix}\sigma_1 & -\sigma_3 & -\sigma_4 & \sigma_D\end{bmatrix}$ and $\partial g(\bar{z}_3, \bar{x}_L) =: \begin{bmatrix}\sigma_5 & -\sigma_6\end{bmatrix}$ such that $\sigma_1, \sigma_4, \sigma_5, \sigma_6, \sigma_D > 0$, and $\sigma_3 \geq 0$. Taking the Laplace transforms yields

$$\begin{aligned} \tilde{x}_L(s) &= e_L^T \left(sI - \bar{A}\right)^{-1} e_1 \tilde{u}(s); \qquad \text{where} \quad \bar{A} := A - \sigma_3 e_1 e_1^T \\ \tilde{u}(s) &= \sigma_1 \tilde{z}_1(s) - \sigma_4 \tilde{x}_L(s) + \sigma_D \tilde{u}_D(s); \qquad \tilde{u}_D(s) = \sigma_5 \tilde{z}_3(s) - \sigma_6 \tilde{x}_L(s) \\ \tilde{z}_1(s) &= \frac{\tilde{\mu}(s) - \eta \bar{z}_1 \tilde{z}_2(s)}{s + \eta \bar{z}_2} \\ \tilde{z}_2(s) &= \frac{\theta \tilde{x}_L(s) - \eta \bar{z}_2 \tilde{z}_1(s)}{s + \eta \bar{z}_1} \\ \tilde{z}_3(s) &= -\frac{\eta_0 \bar{z}_3 \tilde{z}_4(s)}{s + \eta_0 \bar{z}_4} \\ \tilde{z}_4(s) &= \frac{\theta_0 \tilde{u}_D(s) - \eta_0 \bar{z}_4 \tilde{z}_3(s)}{s + \eta_0 \bar{z}_3}. \end{aligned}$$

Next, we express $\tilde{z}_1(s)$ and $\tilde{u}_D(s)$ in terms of $\tilde{\mu}(s), \tilde{x}_L(s)$ and the error $\tilde{e}(s) := \tilde{\mu}(s) - \theta \tilde{x}_L(s)$ as

$$\begin{aligned} \tilde{z}_1(s) &= \left[\tilde{\mu}(s) + \frac{\eta \bar{z}_1}{s} \tilde{e}(s)\right] \frac{1}{s + \eta\left(\bar{z}_1 + \bar{z}_2\right)} \\ \tilde{u}_D(s) &= -\frac{\sigma_6 s}{s + \theta_0 \sigma_5 \frac{\eta_0 \bar{z}_3}{s + \eta_0 (\bar{z}_3 + \bar{z}_4)}} \tilde{x}_L(s). \end{aligned}$$

The feedback control action $\tilde{u}(s)$ can thus be written as

$$\tilde{u}(s) = \left[\sigma_1 \tilde{\mu}(s) + \frac{\eta \sigma_1 \bar{z}_1}{s} \tilde{e}(s)\right] \frac{1}{s + \eta\left(\bar{z}_1 + \bar{z}_2\right)} - \sigma_4 \tilde{x}_L(s) - \frac{\sigma_D \sigma_6 s}{s + \theta_0 \sigma_5 F_{\eta_0}(s)} \tilde{x}_L(s),$$

where $F_{\eta_0}(s) := \frac{\eta_0 \bar{z}_3}{s + \eta_0(\bar{z}_3 + \bar{z}_4)}$. This is the controller transfer function that relates the control action $\tilde{u}$ to the reference signal $\tilde{\mu}$, the error signal $\tilde{e}$ and the plant output $\tilde{x}_L$ in the Laplace domain.

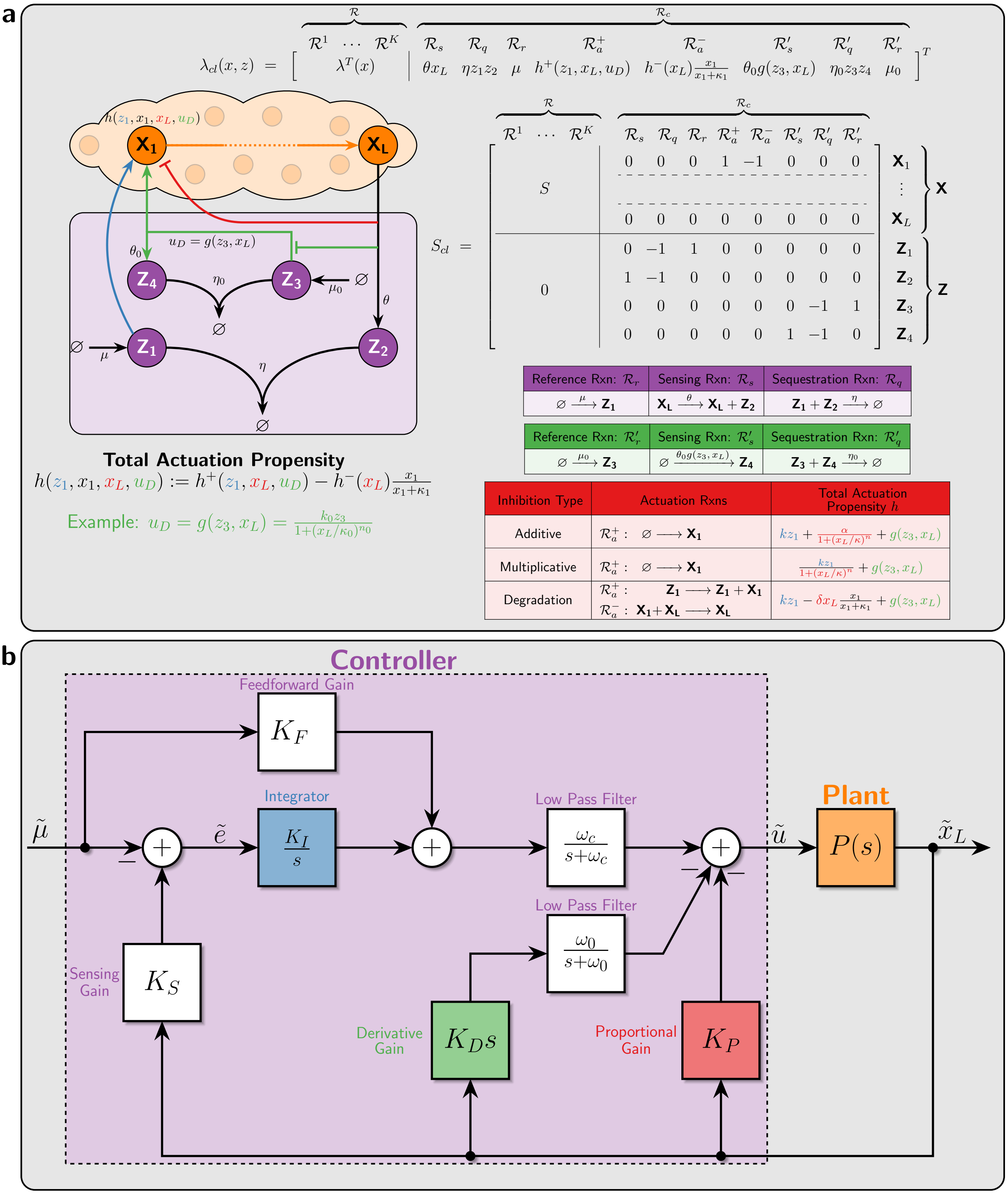

Figure S4: **Fourth-Order *a*PID Feedback Controllers (with Additive D). (a) *a*PID Controller Network.** An additional pathway (marked in green) is appended to the *a*PI controller of class 1 (see Figure 3) via two intermediate controller species $\mathbf{Z_3}$ and $\mathbf{Z_4}$ to embed an additional derivative control action. The obtained derivative mechanism here is fundamentally different from the previous incoherent feedforward mechanisms. Here the derivative operation is achieved by connecting an antithetic integral motif in feedback with itself ($\mathbf{Z_3}$ feeds back into $\mathbf{Z_4}$). **(b) *a*PID Controller Block Diagram.** This block diagram represents the dynamics of the linearized closed-loop system obtained by the fourth-order *a*PID controller with different inhibition mechanisms. An additional low pass filter appears (with cutoff frequency $\omega_0$) which indicates that only the realized D control action is filtered before it actuates the plant. The various gains are expressed in terms of the biomolecular parameters and partial derivatives of $h$ and $g$ in (S8).

**Lemma 1.** *As $\eta_0 \to \infty$, $F_{\eta_0}(s) \to 1$.*

*Proof.* At steady state we have

$$\bar{x}_L = \frac{\mu}{\theta}, \qquad g\left(\bar{z}_3, \frac{\mu}{\theta}\right) = \frac{\mu_0}{\theta_0}, \qquad \bar{z}_3\bar{z}_4 = \frac{\mu_0}{\eta_0}.$$

Hence $\bar{z}_3\bar{z}_4 \to 0$ as $\eta_0 \to \infty$. But $\bar{z}_3$ is independent of $\eta_0$, then $\bar{z}_4 \to 0$ as $\eta_0 \to \infty$. This implies that

$$F_{\eta_0}(s) \underset{\eta_0 \to \infty}{\longrightarrow} \frac{\eta_0 \bar{z}_3}{s + \eta_0 \bar{z}_3} \to 1.$$

□

Therefore, equipped with the controller and plant transfer functions given by (in the limit as $\eta_0 \to \infty$)

$$\begin{aligned} &\textbf{Controller:} \quad \tilde{u}(s) = \left[K_F \tilde{\mu}(s) + \frac{K_I}{s}\tilde{e}(s)\right]\frac{\omega_c}{s+\omega_c} - \left[K_P + K_D s \frac{\omega_0}{s+\omega_0}\right]\tilde{x}_L(s) \\ &\textbf{Plant:} \quad \tilde{x}_L(s) = P(s)\tilde{u}(s); \\ &\text{where:} \quad \begin{cases} K_F = \dfrac{\sigma_1}{\eta(\bar{z}_1+\bar{z}_2)}, \quad K_I = \sigma_1 \dfrac{\bar{z}_1}{\bar{z}_1+\bar{z}_2}, \quad K_S = \theta, \quad K_P = \sigma_4, \\ K_D = \dfrac{\sigma_D \sigma_6}{\theta_0 \sigma_5}, \quad \omega_c = \eta(\bar{z}_1+\bar{z}_2), \quad \omega_0 = \theta_0\sigma_5, \quad P(s) = e_L^T\left(sI - \bar{A}\right)^{-1} e_1, \end{cases} \end{aligned} \tag{S8}$$

we can now draw the block diagram shown in Figure S4(b) which encompasses the proposed fourth-order $a$PID architectures with the different inhibition mechanisms for the proportional component. The first term in the controller transfer function corresponds to the filtered integral action that simplifies to a pure integral action $\frac{K_I}{s}\tilde{e}(s)$ as $\eta \to \infty$. The second term involves a separate proportional and filtered derivative control actions with a cutoff frequency $\omega_0$ as illustrated in the block diagram. In conclusion, the realized control architecture, as $\eta, \eta_0 \to \infty$, is a PI + filtered D.

##### S1.2.4 Fourth-Order $a$PID Feedback Controller (with D Entering as a Degradation)

Consider an arbitrary network controlled by the fourth-order $a$PID controller depicted in Figure S5. The deterministic dynamics of the closed-loop system are given by

$$\begin{cases} \dot{x} = f(x) + h(z_1, x_1, x_L, u_D)e_1; \qquad u_D := g(z_3, x_L) \\ \dot{z}_1 = \mu - \eta z_1 z_2 \\ \dot{z}_2 = \theta x_L - \eta z_1 z_2 \\ \dot{z}_3 = \mu_0 - \eta_0 z_3 z_4 \\ \dot{z}_4 = \theta_0 u_D - \eta_0 z_3 z_4, \end{cases} \tag{S9}$$

where $f(x) = S\lambda(x)$. The function $h$ is chosen to be monotonically increasing (resp. decreasing) in $z_1$ (resp. in $x_1, x_L$ and $u_D$), while $g$ is monotonically increasing $z_3$ and $x_L$ as depicted in the network diagram of Figure S5. An example of the function $g$ is given by

$$g(z_3, x_L) = \alpha_1 z_3 + \alpha_2 x_L.$$

Let $\begin{bmatrix}\tilde{x}^T & \tilde{z}_1 & \tilde{z}_2 & \tilde{z}_3 & \tilde{z}_4 & \tilde{u}_D\end{bmatrix}^T$ denote the perturbation from the fixed point $\begin{bmatrix}\bar{x}^T & \bar{z}_1 & \bar{z}_2 & \bar{z}_3 & \bar{z}_4 & \tilde{u}_D\end{bmatrix}^T$ of (S9) with $\bar{u}_D = g(\bar{z}_3, \bar{x}_L) = \mu_0/\theta_0$. We also assume that the reference signal $\mu$ is allowed to slightly vary in time around a nominal reference $\bar{\mu}$. That is, we have

$$\tilde{u}_D(t) = u_D(t) - \bar{u}_D; \qquad \tilde{x}(t) = x(t) - \bar{x}; \qquad \tilde{\mu}(t) = \mu(t) - \bar{\mu}; \qquad \tilde{z}_i(t) = z_i(t) - \bar{z}_i; \quad (i = 1,2,3,4).$$

The linearized dynamics can thus be written as

$$\begin{cases} \dot{\tilde{x}} = A\tilde{x} + (\sigma_1\tilde{z}_1 - \sigma_3\tilde{x}_1 - \sigma_4\tilde{x}_L)\,e_1 - \sigma_D\tilde{u}_D e_1; \qquad \tilde{u}_D = \sigma_5\tilde{z}_3 + \sigma_6\tilde{x}_L \\ \dot{\tilde{z}}_1 = \tilde{\mu} - \eta\bar{z}_2\tilde{z}_1 - \eta\bar{z}_1\tilde{z}_2 \\ \dot{\tilde{z}}_2 = \theta\tilde{x}_L - \eta\bar{z}_2\tilde{z}_1 - \eta\bar{z}_1\tilde{z}_2 \\ \dot{\tilde{z}}_3 = -\eta_0\bar{z}_4\tilde{z}_3 - \eta_0\bar{z}_3\tilde{z}_4 \\ \dot{\tilde{z}}_4 = \theta_0\tilde{u}_D - \eta_0\bar{z}_4\tilde{z}_3 - \eta_0\bar{z}_3\tilde{z}_4, \end{cases}$$

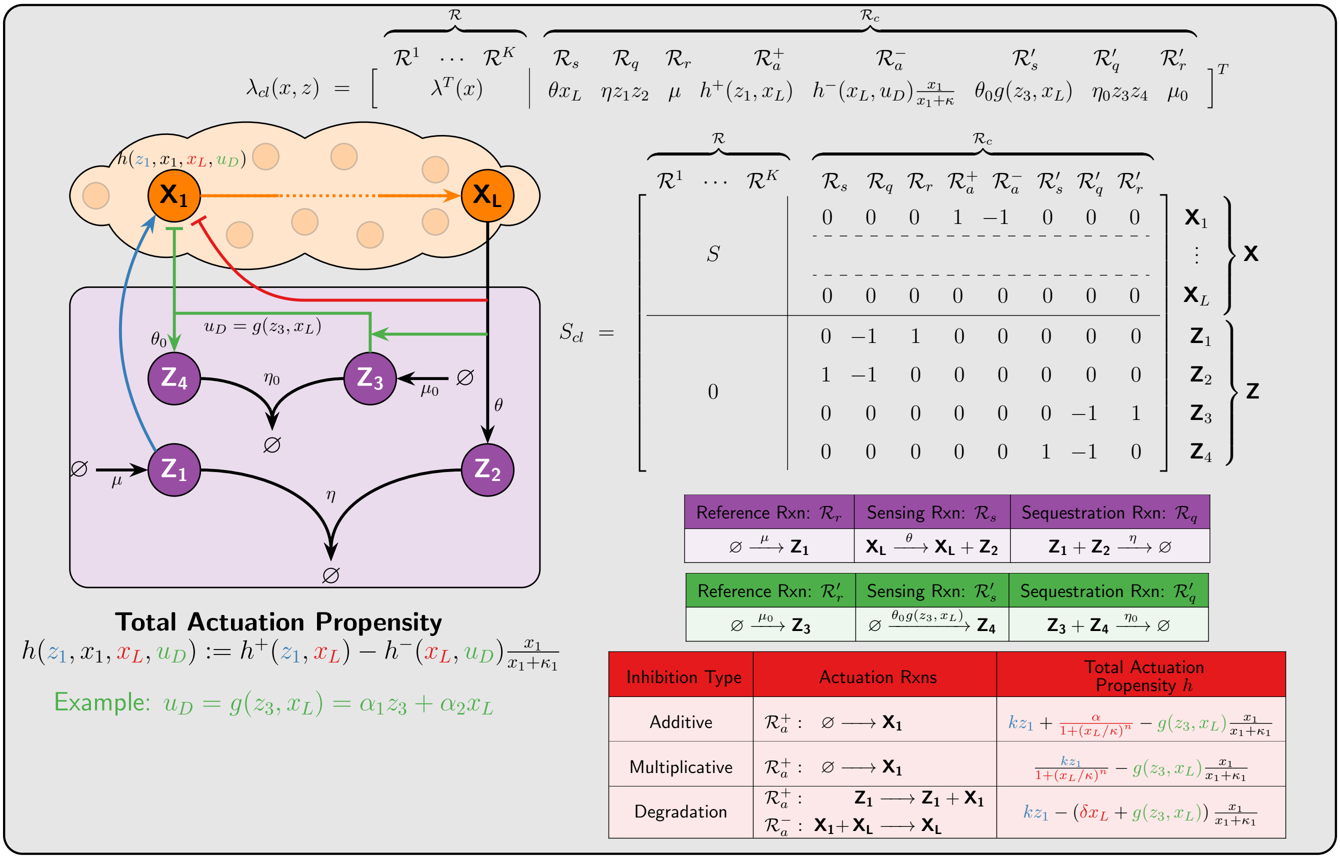

Figure S5: **Fourth-Order *a*PID Feedback Controllers (with the Derivative Entering as a Degradation). (a) *a*PID Controller Network.** There are two differences between this controller and the one proposed in Figure S4. First, the actuation by the D component is carried out as a degradation reaction. More precisely, the input species $\mathbf{X_1}$ is degraded at a rate $g(z_3, x_L)$. Furthermore, $g$ has to be a monotonically increasing function of $z_3$ and $x_L$. The resulting block diagram of the linearized dynamics is exactly the same as that given in Figure S4(b).

where $A := \partial f(\bar{x})$, $\partial h(\bar{z}_1, \bar{x}_1, \bar{x}_L, \bar{u}_D) =: \begin{bmatrix} \sigma_1 & -\sigma_3 & -\sigma_4 & -\sigma_D \end{bmatrix}$ and $\partial g(\bar{z}_3, \bar{x}_L) =: \begin{bmatrix} \sigma_5 & \sigma_6 \end{bmatrix}$ such that $\sigma_1, \sigma_4, \sigma_5, \sigma_6, \sigma_D > 0$, and $\sigma_3 \geq 0$. Taking the Laplace transforms yields

$$
\begin{aligned}
\tilde{x}_L(s) &= e_L^T \left(sI - \bar{A}\right)^{-1} e_1 \tilde{u}(s); \qquad \text{where} \quad \bar{A} := A - \sigma_3 e_1 e_1^T \\
\tilde{u}(s) &= \sigma_1 \tilde{z}_1(s) - \sigma_4 \tilde{x}_L(s) - \sigma_D \tilde{u}_D(s); \qquad \tilde{u}_D(s) = \sigma_5 \tilde{z}_3(s) + \sigma_6 \tilde{x}_L(s) \\
\tilde{z}_1(s) &= \frac{\tilde{\mu}(s) - \eta \bar{z}_1 \tilde{z}_2(s)}{s + \eta \bar{z}_2} \\
\tilde{z}_2(s) &= \frac{\theta \tilde{x}_L(s) - \eta \bar{z}_2 \tilde{z}_1(s)}{s + \eta \bar{z}_1} \\
\tilde{z}_3(s) &= -\frac{\eta_0 \bar{z}_3 \tilde{z}_4(s)}{s + \eta_0 \bar{z}_4} \\
\tilde{z}_4(s) &= \frac{\theta_0 \tilde{u}_D(s) - \eta_0 \bar{z}_4 \tilde{z}_3(s)}{s + \eta_0 \bar{z}_3}.
\end{aligned}
$$

Next, we express $\tilde{z}_1(s)$ and $\tilde{u}_D(s)$ in terms of $\tilde{\mu}(s), \tilde{x}_L(s)$ and the error $\tilde{e}(s) := \tilde{\mu}(s) - \theta \tilde{x}_L(s)$ as

$$
\begin{aligned}
\tilde{z}_1(s) &= \left[\tilde{\mu}(s) + \frac{\eta \bar{z}_1}{s} \tilde{e}(s)\right] \frac{1}{s + \eta \left(\bar{z}_1 + \bar{z}_2\right)} \\
\tilde{u}_D(s) &= \frac{\sigma_6 s}{s + \theta_0 \sigma_5 \frac{\eta_0 \bar{z}_3}{s + \eta_0 (\bar{z}_3 + \bar{z}_4)}} \tilde{x}_L(s).
\end{aligned}
$$

The feedback control action $\tilde{u}(s)$ is thus the same as that in additive derivative design. Therefore, the block diagram and PID gains are also the same and are given in Figure S4(b) and (S8), respectively. The only differences are in the signs of the partial derivatives of $g$, and the partial derivative of $h$ with respect to $u_D$.

### S2 Plants with Negative Gains: P-Type *a*PID Feedback Controllers

There are two controller implementation types to be considered: N-Type and P-Type. N-Type (Negative feedback) controllers are suitable for positive gain plants, while P-Type (Positive feedback) controllers are suitable for negative gain plants. This ensures the overall control loop implements negative feedback. The various *a*PID control architectures proposed and analyzed in Section S1 consider only N-Type controller implementations. In this section, we provide their P-Type counterparts, depicted in Figure S6, that are usually suitable for plants with negative gains.

The PID structures of the various P-Type controllers proposed in Figure S6 can be revealed once again using a linear perturbation analysis. The same derivations as those in Section S1.2 are carried out to yield the following results.

**Simple Second-Order *a*PID:** The deterministic dynamics of the closed-loop system are given by

$$\begin{cases} \dot{x} = f(x) + h(z_2, x_L)e_1 \\ \dot{z}_1 = \mu + \beta x_L - \eta z_1 z_2 \\ \dot{z}_2 = \theta x_L - \eta z_1 z_2, \end{cases} \tag{S10}$$

where $f(x) = S\lambda(x)$, and $h$ is a monotonically increasing function in $z_2$ and $x_L$. The transfer functions are given by

$$\begin{aligned} &\textbf{Controller:} \quad \tilde{u}(s) = \left[\frac{K_I}{s}\tilde{e}(s) + \Big(K_P + K_D s\Big)\tilde{x}_L(s)\right]\frac{\omega_c}{s+\omega_c}; \qquad \tilde{e}(s) := K_S\tilde{x}_L(s) - \tilde{\mu}(s) \\ &\textbf{Plant:} \quad \tilde{x}_L(s) = P(s)\tilde{u}(s); \\ &\text{where:} \quad \begin{cases} K_I = \sigma_1 \dfrac{\bar{z}_2}{\bar{z}_1 + \bar{z}_2}, \quad K_S = \theta - \beta, \qquad K_P = \sigma_4 + \dfrac{\sigma_1\theta}{\eta(\bar{z}_1 + \bar{z}_2)}, \\ K_D = \dfrac{\sigma_4}{\eta(\bar{z}_1 + \bar{z}_2)} \quad \omega_c = \eta(\bar{z}_1 + \bar{z}_2), \quad P(s) = e_L^T\left(sI - A\right)^{-1}e_1, \end{cases} \end{aligned} \tag{S11}$$

where $A := \partial f(\bar{x})$, $\partial h(\bar{z}_2, \bar{x}_L) =: \begin{bmatrix}\sigma_1 & \sigma_4\end{bmatrix}$ such that $\sigma_1 > 0$ and $\sigma_4 \geq 0$.

**Third-Order *a*PID:** The deterministic dynamics of the closed-loop system are given by

$$\begin{cases} \dot{x} = f(x) + h(z_2, z_3, x_1, x_L)e_1 \\ \dot{z}_1 = \mu - \eta z_1 z_2 \\ \dot{z}_2 = \theta x_L - \eta z_1 z_2 \\ \dot{z}_3 = g(z_3, x_L) - \gamma z_3, \end{cases} \tag{S12}$$

where $f(x) = S\lambda(x)$. The function $h$ is chosen to be monotonically increasing (resp. decreasing) in $z_2$ and $x_L$ (resp. in $x_1$), $g$ is monotonically decreasing in $z_3$, while $g$ and $h$ are chosen to have the opposite monotonicity in $x_L$ and $z_3$ respectively (see the diamond arrowhead in Figure S6). The transfer functions are given by

$$\begin{aligned} &\textbf{Controller:} \quad \tilde{u}(s) = \left[K_0\tilde{x}_L(s) + \frac{K_I}{s}\tilde{e}(s)\right]\frac{\omega_c}{s+\omega_c} + \left[K_P + K_D s\right]\frac{\omega_0}{s+\omega_0}\tilde{x}_L(s); \qquad \tilde{e}(s) := K_S\tilde{x}(s) - \tilde{\mu}(s) \\ &\textbf{Plant:} \quad \tilde{x}_L(s) = P(s)\tilde{u}(s); \\ &\text{where:} \quad \begin{cases} K_F = \dfrac{\sigma_1\theta}{\eta(\bar{z}_1 + \bar{z}_2)}, \quad K_I = \sigma_1\dfrac{\bar{z}_2}{\bar{z}_1 + \bar{z}_2}, \quad K_S = \theta, \qquad K_P = \sigma_4 + \dfrac{\sigma_2\sigma_6}{\gamma + \sigma_5}, \\ K_D = \dfrac{\sigma_4}{\gamma + \sigma_5} \qquad \omega_c = \eta(\bar{z}_1 + \bar{z}_2), \quad \omega_0 = \gamma + \sigma_5, \quad P(s) = e_L^T\left(sI - \bar{A}\right)^{-1}e_1, \end{cases} \end{aligned} \tag{S13}$$

where $A := \partial f(\bar{x})$, $\partial h(\bar{z}_2, \bar{z}_3, \bar{x}_1, \bar{x}_L) =: \begin{bmatrix}\sigma_1 & \sigma_2 & -\sigma_3 & \sigma_4\end{bmatrix}$, $\bar{A} := A - \sigma_3 e_1 e_1^T$ and $\partial g(\bar{z}_3, \bar{x}_L) =: \begin{bmatrix}-\sigma_5 & \sigma_6\end{bmatrix}$ such that $\sigma_1, \sigma_4 > 0$, $\sigma_3, \sigma_5 \geq 0$ and $\sigma_2\sigma_6 < 0$.

**Fourth-Order *a*PID:** The deterministic dynamics of the closed-loop system are given by

$$\begin{cases} \dot{x} = f(x) + h(z_2, x_L, u_D)e_1; \qquad u_D = g(z_3, x_L) \\ \dot{z}_1 = \mu - \eta z_1 z_2 \\ \dot{z}_2 = \theta x_L - \eta z_1 z_2 \\ \dot{z}_3 = \mu_0 - \eta_0 z_3 z_4 \\ \dot{z}_4 = \theta_0 u_D - \eta_0 z_3 z_4, \end{cases} \tag{S14}$$

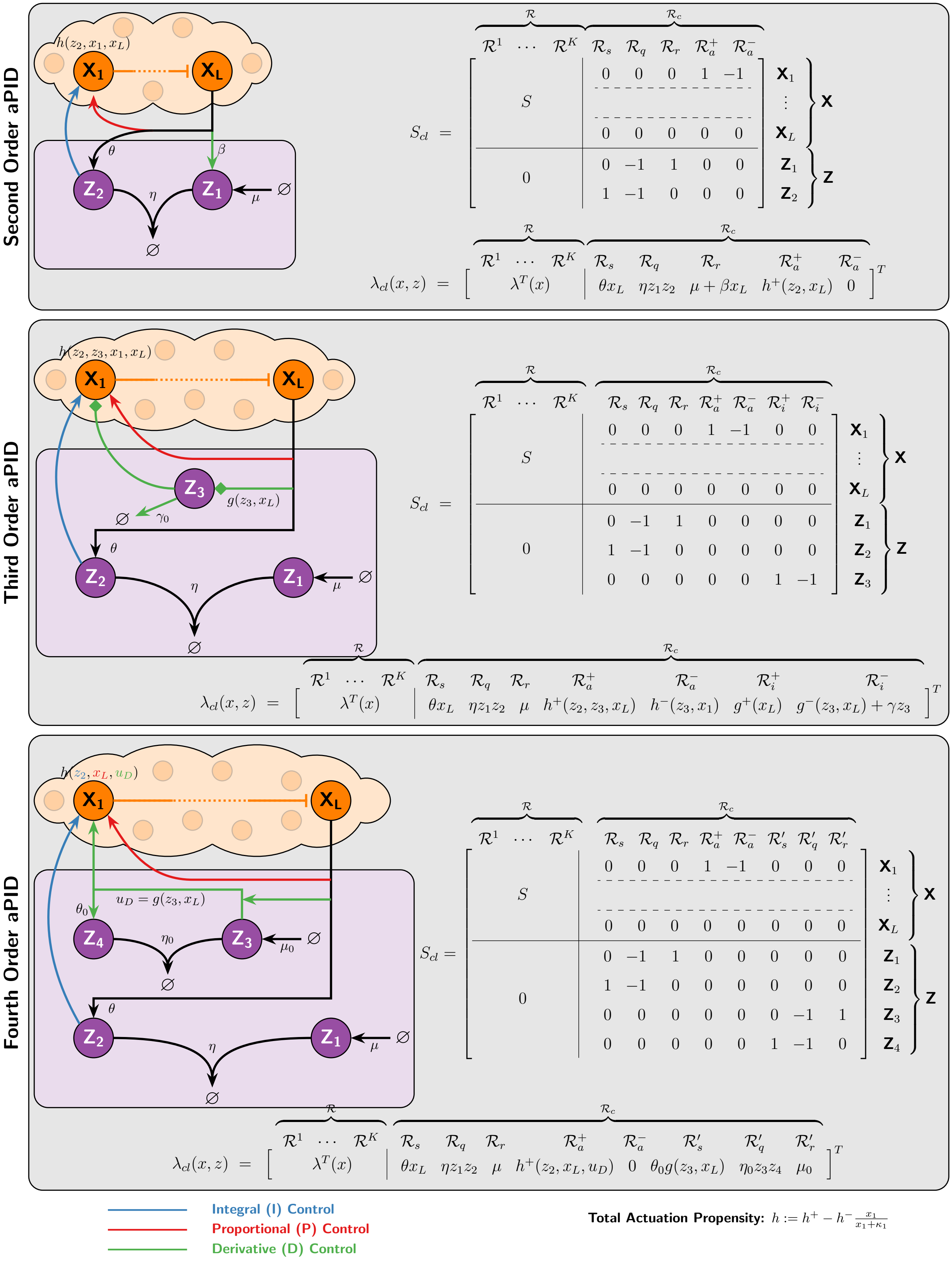

Figure S6: **P-Type *a*PID Controllers.**

where $f(x) = S\lambda(x)$. The function $h$ is chosen to be monotonically increasing in $z_1, x_L$ and $u_D$, while $g$ is monotonically increasing in $z_3$ and $x_L$ as depicted in the network diagram of Figure S6. The transfer functions are given by

$$\begin{aligned}
&\textbf{Controller:} \quad \tilde{u}(s) = \left[K_0\tilde{\mu}(s) + \frac{K_I}{s}\tilde{e}(s)\right]\frac{\omega_c}{s+\omega_c} + \left[K_P + K_D s\frac{\omega_0}{s+\omega_0}\right]\tilde{x}_L(s); \qquad \tilde{e}(s) := K_S\tilde{x}_L(s) - \tilde{\mu}(s)\\
&\textbf{Plant:} \quad \tilde{x}_L(s) = P(s)\tilde{u}(s);\\
&\text{where:} \quad \begin{cases} K_0 = \dfrac{\sigma_1\theta}{\eta(\bar{z}_1+\bar{z}_2)}, & K_I = \sigma_1\dfrac{\bar{z}_2}{\bar{z}_1+\bar{z}_2}, \quad K_S = \theta, \quad K_P = \sigma_4,\\ K_D = \dfrac{\sigma_D\sigma_6}{\theta_0\sigma_5}, & \omega_c = \eta(\bar{z}_1+\bar{z}_2), \quad \omega_0 = \theta_0\sigma_5, \quad P(s) = e_L^T(sI-A)^{-1}e_1, \end{cases}
\end{aligned} \tag{S15}$$

where $A := \partial f(\bar{x})$, $\partial h(\bar{z}_1, \bar{x}_L, \bar{u}_D) =: \begin{bmatrix}\sigma_1 & \sigma_4 & \sigma_D\end{bmatrix}$ and $\partial g(\bar{z}_3, \bar{x}_L) =: \begin{bmatrix}\sigma_5 & \sigma_6\end{bmatrix}$ such that $\sigma_1, \sigma_4, \sigma_5, \sigma_6, \sigma_D > 0$.

#### S3 Derivation of the Stability Conditions of the *a*PI Controllers

Consider the closed-loop network in Figure 4(a) where the *a*PI controller is in feedback with a particular plant comprised of two species $\mathbf{X_1}$ and $\mathbf{X_2}$. We now exploit the Routh-Hurwitz Criterion to derive a necessary and sufficient stability condition for all of the proposed *a*PI controllers. Using the closed-loop stoichiometry matrix and propensity function, shown in Figure 4, one can write the deterministic closed-loop dynamics as

$$\begin{cases} \dot{x}_1 = h(z_1, z_2, x_1, x_2) - \gamma_1 x_1\\ \dot{x}_2 = k_1 x_1 - \gamma_2 x_2\\ \dot{z}_1 = g(\mu, x_2) - \eta z_1 z_2\\ \dot{z}_2 = \theta x_2 - \eta z_1 z_2. \end{cases} \tag{S16}$$

Hence, the Jacobian $J$, evaluated at the fixed point $\begin{bmatrix}\bar{x}_1 & \bar{x}_2 & \bar{z}_1 & \bar{z}_2\end{bmatrix}^T$, of the right hand side is calculated to be

$$J = \begin{bmatrix} -(\gamma_1+\sigma_3) & -\sigma_4 & \sigma_1 & -\sigma_2\\ k_1 & -\gamma_2 & 0 & 0\\ 0 & -\sigma_6 & -\eta\bar{z}_2 & -\eta\bar{z}_1\\ 0 & \theta & -\eta\bar{z}_2 & -\eta\bar{z}_1 \end{bmatrix},$$

where $\partial h(\bar{z}_1, \bar{z}_2, \bar{x}_1, \bar{x}_2) =: \begin{bmatrix}\sigma_1 & -\sigma_2 & -\sigma_3 & -\sigma_4\end{bmatrix}$, and $\partial g(\mu, \bar{x}_2) =: \begin{bmatrix}\sigma_5 & -\sigma_6\end{bmatrix}$. Note that $\sigma_i \geq 0$. The characteristic polynomial $p(s) := \det(sI - J)$ is thus calculated to be

$$\begin{aligned} p(s) = s^4 + (\eta\bar{z} + \bar{\gamma}_1)s^3 + (\eta\bar{z}\bar{\gamma}_1 + \bar{\gamma}_2)s^2 + [\eta\bar{z}\bar{\gamma}_2 + k_1(\theta\sigma_2 + \sigma_1\sigma_6)]s + \eta k_1(\theta+\sigma_6)(\sigma_1\bar{z}_1 + \sigma_2\bar{z}_2)\\ \text{where} \quad \bar{z} := \bar{z}_1 + \bar{z}_2, \quad \bar{\gamma}_1 := \gamma_1 + \gamma_2 + \sigma_3, \quad \text{and} \quad \bar{\gamma}_2 := (\gamma_1+\sigma_3)\gamma_2 + k_1\sigma_4. \end{aligned}$$

Using the characteristic polynomial we construct the Routh-Hurwitz table:

| | | | |
|---|---|---|---|
| $s^4$ | $1$ | $\eta\bar{z}\bar{\gamma}_1 + \bar{\gamma}_2$ | $\eta k_1(\theta+\sigma_6)(\sigma_1\bar{z}_1+\sigma_2\bar{z}_2)$ |
| $s^3$ | $\eta\bar{z}+\bar{\gamma}_1$ | $\eta\bar{z}\bar{\gamma}_2 + k_1(\theta\sigma_2+\sigma_1\sigma_6)$ | $0$ |
| $s^2$ | $\frac{\eta\bar{z}\bar{\gamma}_1(\eta\bar{z}+\bar{\gamma}_1)+\bar{\gamma}_1\bar{\gamma}_2-k_1(\theta\sigma_2+\sigma_1\sigma_6)}{\eta\bar{z}+\bar{\gamma}_1}$ | $\eta k_1(\theta+\sigma_6)(\sigma_1\bar{z}_1+\sigma_2\bar{z}_2)$ | $0$ |
| $s^1$ | $\eta\bar{z}\bar{\gamma}_2 + k_1(\theta\sigma_2+\sigma_1\sigma_6) - \frac{\eta k_1(\theta+\sigma_6)(\sigma_1\bar{z}_1+\sigma_2\bar{z}_2)(\eta\bar{z}+\bar{\gamma}_1)^2}{\eta\bar{\gamma}_1\bar{z}(\eta\bar{z}+\bar{\gamma}_1)+\bar{\gamma}_1\bar{\gamma}_2-k_1(\theta\sigma_2+\sigma_1\sigma_6)}$ | $0$ | $0$ |
| $s^0$ | $\eta k_1(\theta+\sigma_6)(\sigma_1\bar{z}_1+\sigma_2\bar{z}_2)$ | $0$ | $0$ |

The Routh-Hurwitz criterion states that the necessary and sufficient condition for local stability of the fixed point is obtained by forcing all the entries of the first column to be positive. Therefore the exact necessary and sufficient conditions for local stability of the fixed point are

$$\eta\bar{z}\bar{\gamma}_1(\eta\bar{z}+\bar{\gamma}_1)+\bar{\gamma}_1\bar{\gamma}_2 > k_1(\theta\sigma_2+\sigma_1\sigma_6) \quad \text{and} \quad \eta\bar{z}\bar{\gamma}_2+k_1(\theta\sigma_2+\sigma_1\sigma_6) > \frac{\eta k_1(\theta+\sigma_6)(\sigma_1\bar{z}_1+\sigma_2\bar{z}_2)(\eta\bar{z}+\bar{\gamma}_1)^2}{\eta\bar{\gamma}_1\bar{z}(\eta\bar{z}+\bar{\gamma}_1)+\bar{\gamma}_1\bar{\gamma}_2-k_1(\theta\sigma_2+\sigma_1\sigma_6)}. \tag{S17}$$

These stability conditions are cumbersome and not easy to compare for different *a*PI controllers (that is, different $\sigma_i$'s and $\bar{z}$). However, one can obtain a simpler but approximate stability condition for the case of strong sequestration (asymptotic limit as $\eta \to \infty$). In fact, the first condition in (S17) is always satisfied when $\eta \to \infty$ assuming that $\sigma_1, \sigma_2$

and $\sigma_6$ are finite (which can be easily checked to be a valid assumption for the proposed $a$PI controllers). To simplify the second condition in (S17), we first examine the asymptotic behavior of $\bar{z}_1$ and $\bar{z}_2$ as $\eta \to \infty$. For any $\eta > 0$, we have that $\bar{z}_1\bar{z}_2 = \theta\bar{x}_2/\eta$. Thus, as long as $\bar{x}_2$ is finite, we have that $\bar{z}_1\bar{z}_2 = 0$ in the limit as $\eta \to \infty$. This means that we have two possibilities in the asymptotic limit as $\eta \to \infty$: if $\bar{z}_1 > 0$ then $\bar{z}_2 = 0$, and if $\bar{z}_2 > 0$ then $\bar{z}_1 = 0$. As a result, as $\eta \to \infty$, the second condition in (S17) takes one of the two forms

$$\begin{cases} \sigma_1 k_1(\theta + \sigma_6) < \bar{\gamma}_1\bar{\gamma}_2 & \text{and} \quad (\bar{z}_1 > 0, \bar{z}_2 = 0) \\ \sigma_2 k_1(\theta + \sigma_6) < \bar{\gamma}_1\bar{\gamma}_2 & \text{and} \quad (\bar{z}_1 = 0, \bar{z}_2 > 0). \end{cases}$$

This can be rewritten as

$$\begin{cases} \sigma_1 k_1(\theta + \sigma_6) < (\gamma_1 + \gamma_2 + \sigma_3)(\gamma_1\gamma_2 + \sigma_3\gamma_2 + k_1\sigma_4) & \text{and} \quad (\bar{z}_1 > 0, \bar{z}_2 = 0) \\ \sigma_2 k_1(\theta + \sigma_6) < (\gamma_1 + \gamma_2 + \sigma_3)(\gamma_1\gamma_2 + \sigma_3\gamma_2 + k_1\sigma_4) & \text{and} \quad (\bar{z}_1 = 0, \bar{z}_2 > 0). \end{cases} \tag{S18}$$

Note that the conditions $z_1 > 0$ and $z_2 > 0$ prevent instabilities that arise due to negative fixed points. Equations (S18) provide the approximate stability conditions compactly for all the proposed $a$PI controllers. To this end, the stability conditions for each $a$PI controller are obtained by simply computing $\sigma_i$'s, $\bar{z}_1$, and $\bar{z}_2$ for each particular controller architecture, and then take the limit as $\eta \to \infty$. The calculation results are given in Table S1, where $\bar{x}_L = r$ is the desired set-point and is equal to $\mu/\theta$ for $a$PI controllers of Class 1 and 2. For the $a$PI controllers of Class 3, the set-point $\bar{x}_L = r$ solves a polynomial equation of degree $n+1$ given by

$$\begin{aligned} &\text{Additive Inhibition:} && \bar{x}_L^{n+1} - \frac{\mu}{\theta}\bar{x}_L^n + \kappa^n\bar{x}_L - \kappa^n\frac{\mu+\alpha}{\theta} = 0 \\ &\text{Multiplicative Inhibition:} && \bar{x}_L^{n+1} + \kappa^n\bar{x}_L - \kappa^n\frac{\mu}{\theta} = 0. \end{aligned} \tag{S19}$$

For example, for a hill coefficient $n = 1$, the unique non-negative steady state (assuming closed-loop stability) of the output concentration is given by

$$\begin{aligned} &\text{Additive Inhibition:} && \bar{x}_L = \frac{\frac{\mu}{\theta} - \kappa + \sqrt{\left(\kappa + \frac{\mu}{\theta}\right)^2 + 4\kappa\frac{\mu}{\theta}}}{2} =: r \\ &\text{Multiplicative Inhibition:} && \bar{x}_L = \frac{\kappa}{2}\left(\sqrt{1 + \frac{4}{\kappa}\frac{\mu}{\theta}} - 1\right) =: r. \end{aligned} \tag{S20}$$

Based on the table and (S18), one can write the stability conditions for each $a$PI controller. The results are tabulated in Figure 4(b).

### S4 Mappings between the PID and Biomolecular Parameter Spaces

In this section, we unravel the mappings between the PID parameters $(K_P, K_I, K_D, \omega_c, \omega_0)$ and the various biomolecular parameters $(k, \eta, \theta, \beta, \delta, ...)$. We first start with the analysis problem: Given the biomolecular parameters, what are the PID gains and cutoff frequency? Then we move to the design problem: What are the biomolecular parameters that achieve some desired PID gains and cutoff frequency?

Throughout the subsequent analysis, we will make an assumption about the plant to be controlled as detailed next. Let the plant be comprised of $L$ species $\mathbf{X} = \{\mathbf{X_1}, \mathbf{X_2}, ..., \mathbf{X_L}\}$ interacting with each other via a biochemical reaction network described by a general stoichiometry matrix $S$ and a propensity function $\lambda$ as depicted in Figure 2(a). Let $u$ denote the actuation input of the plant that enters the plant dynamics as the total actuation propensity function affecting the production and/or degradation of the input species $\mathbf{X_1}$. Furthermore, let $\mathbf{X_L}$ denote the output of interest. Hence the deterministic dynamics of the plant, regardless of what the controller is, are governed by the following ODE:

$$\dot{x} = S\lambda(x) + ue_1, \tag{S21}$$

where $e_i$ is a vector whose entries are all zeros except the $i^{\text{th}}$ entry being equal to one. Let $F_i$ $(i = 1, 2, ..., L)$ denote the steady-state maps of the plant, that is, if $u$ is a constant then

$$\begin{cases} S\lambda(x) + ue_1 = 0 \\ \qquad x_i = e_i^T x. \end{cases} \implies \qquad x_i = F(u). \tag{S22}$$

| Class | Inhibition | $\sigma_i$'s | $\bar{z}_1(\eta \to \infty)$ | $\bar{z}_2(\eta \to \infty)$ |
|---|---|---|---|---|
| **AIF** | None | $\sigma_1 = k, \quad \sigma_5 = 1$<br>$\sigma_2 = \sigma_3 = \sigma_4 = \sigma_6 = 0$ | $\frac{\gamma_1\gamma_2 r}{kk_1}$ | 0 |
| **APIF Class 1** | Additive | $\sigma_1 = k, \quad \sigma_5 = 1$<br>$\sigma_4 = \frac{\alpha}{\kappa}\frac{n(r/\kappa)^{n-1}}{[1+(r/\kappa)^n]^2}$<br>$\sigma_2 = \sigma_3 = \sigma_6 = 0$ | $\frac{1}{k}\left[\frac{\gamma_1\gamma_2 r}{k_1} - \frac{\alpha}{1+(r/\kappa)^n}\right]$ | 0 |
| | Multiplicative | $\sigma_1 = \frac{k}{1+(r/\kappa)^n}, \quad \sigma_5 = 1$<br>$\sigma_4 = \frac{\gamma_1\gamma_2}{k_1}\frac{n(r/\kappa)^n}{1+(r/\kappa)^n}$<br>$\sigma_2 = \sigma_3 = \sigma_6 = 0$ | $\frac{\gamma_1\gamma_2 r}{kk_1}\left[1+(r/\kappa)^n\right]$ | 0 |
| | Degradation | $\sigma_1 = k, \quad \sigma_5 = 1, \ \sigma_2 = \sigma_6 = 0$<br>$\sigma_4 = r\frac{\delta\gamma_2}{\gamma_2 r + \kappa_1 k_1}$<br>$\sigma_3 = \frac{\delta r \kappa_1 k_1^2}{(\gamma_2 r + \kappa_1 k_1)^2}$ | $\frac{\gamma_2 r}{kk_1}\left(\gamma_1 + \frac{\delta r}{\kappa_1 + \gamma_2 r/k_1}\right)$ | 0 |
| **APIF Class 2** | Additive | $\sigma_1 = k, \quad \sigma_5 = 1$<br>$\sigma_2 = \frac{\alpha}{\kappa}\frac{n(\bar{z}_2/\kappa)^n}{1+(\bar{z}_2/\kappa)^n}$<br>$\sigma_3 = \sigma_4 = \sigma_6 = 0$ | $\begin{cases} \frac{1}{k}\left(\frac{\gamma_1\gamma_2 r}{k_1} - \alpha\right) & \text{if} \quad \alpha < \frac{\gamma_1\gamma_2 r}{k_1} \\ 0 & \text{if} \quad \alpha > \frac{\gamma_1\gamma_2 r}{k_1} \end{cases}$ | $\begin{cases} 0 & \text{if} \quad \alpha < \frac{\gamma_1\gamma_2 r}{k_1} \\ \kappa\sqrt[n]{\frac{\alpha k_1}{\gamma_1\gamma_2 r} - 1} & \text{if} \quad \alpha > \frac{\gamma_1\gamma_2 r}{k_1} \end{cases}$ |
| | Multiplicative | $\sigma_1 = k, \quad \sigma_5 = 1$<br>$\sigma_2 = \frac{k\bar{z}_1}{\kappa}\frac{n(\bar{z}_2/\kappa)^{n-1}}{[1+(\bar{z}_2/\kappa)^n]^2}$<br>$\sigma_3 = \sigma_4 = \sigma_6 = 0$ | $\frac{\gamma_1\gamma_2 r}{kk_1}$ | 0 |
| | Degradation | $\sigma_1 = k, \quad \sigma_5 = 1$<br>$\sigma_2 = \delta\frac{\bar{x}_1}{\bar{x}_1 + \kappa_1},$<br>$\sigma_4 = \sigma_6 = 0, \quad \sigma_3 = \delta\bar{z}_2\frac{\kappa_1}{(\bar{x}_1+\kappa_1)^2}$ | $\frac{\gamma_1\gamma_2 r}{kk_1}$ | 0 |
| **APIF Class 3** | Additive | $\sigma_1 = k, \quad \sigma_5 = 1$<br>$\sigma_6 = \frac{\alpha}{\kappa}\frac{n(r/\kappa)^{n-1}}{[1+(r/\kappa)^n]^2}$<br>$\sigma_2 = \sigma_3 = \sigma_4 = 0$ | $\frac{\gamma_1\gamma_2 r}{kk_1}$ | 0 |
| | Multiplicative | $\sigma_1 = k, \quad \sigma_5 = \frac{1}{1+(r/\kappa)^n}$<br>$\sigma_6 = \frac{\mu}{\kappa}\frac{n(r/\kappa)^{n-1}}{[1+(r/\kappa)^n]^2}$<br>$\sigma_2 = \sigma_3 = \sigma_4 = 0$ | $\frac{\gamma_1\gamma_2 r}{kk_1}$ | 0 |

Table S1: Calculation results of $\bar{z}_1$, $\bar{z}_2$ and the various partial derivatives $\sigma_i$'s of the functions $h$ and $g$ evaluated at the fixed point. The calculations are carried out for the particular plant given in Figure 4(a) and for the all the proposed $a$PI controllers with different inhibition mechanisms.

**Assumption 1.** *Assume that for the desired steady-state output $\bar{x}_L = r$, there exists a supporting input, denoted by $\bar{u}$, that achieves the desired output. More precisely, for $r > 0, \exists \bar{u} \in \mathcal{U} \subset \mathbb{R}$ such that $F_L(\bar{u}) = r$, where $\mathcal{U}$ represents the set of admissible inputs.*

**Assumption 2.** *Assume that for the supporting input $\bar{u}$, the steady-state concentration of the input species, denoted by $\bar{x}_1$, exists and is non-negative. More precisely, for $r > 0, \exists \bar{u} \in \mathcal{U} \subset \mathbb{R}$ and $\bar{x}_1 \geq 0$ such that $F_L(\bar{u}) = r$ and $\bar{x}_1 = F_1(\bar{u})$.*

We emphasize that these assumptions do not depend on the type of controller. They only depend on the plant and the particular choice of actuated input species. These assumptions has to be satisfied, otherwise the choice of the input species is simply inadequate. That is, if these assumptions are not satisfied then any type of controller can not achieve the desired output without changing the choice of the input species. In the subsequent analysis, we let $\mathcal{U} = \mathbb{R}_+$ for simplicity. However, this can be relaxed to include negative inputs as well, but with a lower bound that depends on the particular actuation mechanism.

Next, we treat the analysis and design problems for each *a*PID controller separately. Due to the superiority of degradation inhibition demonstrated in Figure 4, we consider degradation inhibitions whenever possible. The other inhibition mechanisms can be similarly treated as well.

#### S4.1 Second Order *a*PID Feedback Controller

Consider an arbitrary plant satisfying Assumptions 1 and 2 controlled with a second order *a*PID controller with degradation inhibition (see Figure S2 (a)). The closed loop dynamics are thus given by

$$\begin{cases} \dot{x} = S\lambda(x) + ue_1; \qquad u = kz_1 - \delta x_L \dfrac{x_1}{x_1 + \kappa_1} \\ \dot{z}_1 = \mu + \beta x_L - \eta z_1 z_2 \\ \dot{z}_2 = \theta x_L - \eta z_1 z_2. \end{cases} \tag{S23}$$

The set-point is given by $\bar{x}_L := r = \mu/(\theta - \beta)$ with $\theta > \beta$. For a given plant and set-point $r$, the supporting input $\bar{u}$ and input species concentration $\bar{x}_1$ are both fixed (see Assumptions 1 and 2). We first treat the analysis problem, then move on to the design problem.

**Analysis:** The controller coordinates $(\bar{z}_1, \bar{z}_2)$ of the fixed point are given by

$$\begin{cases} \bar{z}_1 = \dfrac{1}{k}\left(\bar{u} + \delta r \dfrac{\bar{x}_1}{\bar{x}_1 + \kappa_1}\right) \\ \bar{z}_2 = \dfrac{\mu + \beta r}{\eta \bar{z}_1}. \end{cases} \tag{S24}$$

Clearly, $\bar{z}_1, \bar{z}_2 \geq 0$ when $\bar{u} \in \mathcal{U} = \mathbb{R}_+$. Hence, using the formulas derived in (S4), one can write the PID gains $(K_P, K_I, K_D)$ and cutoff frequency $\omega_c$ in terms of the various biochemical parameters as

$$K_I = k\frac{\bar{z}_1}{\bar{z}_1 + \bar{z}_2}, \quad K_P = \delta T(\bar{x}_1) - \frac{k\beta}{\eta(\bar{z}_1 + \bar{z}_2)}, \quad K_D = \frac{\delta T(\bar{x}_1)}{\eta(\bar{z}_1 + \bar{z}_2)}, \quad \omega_c = \eta(\bar{z}_1 + \bar{z}_2), \tag{S25}$$

where $T(\bar{x}_1) := \frac{\bar{x}_1}{\bar{x}_1 + \kappa_1} \approx 1$ (by choosing $\kappa_1$ to be small enough), and $(\bar{z}_1, \bar{z}_2)$ are given in (S24). Clearly for this *a*PID controller, the gains and cutoff frequency depend on the plant via the supporting input $\bar{u}$.

**Design:** By fixing $\kappa_1, \mu$ and $r$ (and thus $\bar{u}$ and $\bar{x}_1$), one can solve the nonlinear algebraic equations given in (S25) and (S24) for the biomolecular parameters $\delta, \eta, k, \beta$ and $\theta$ in terms of the PID gains and cutoff frequency. The derivations are slightly tedious, but the results are provided here

$$\begin{cases} \delta = K_D\omega_c\left(1 + \kappa_1/\bar{x}_1\right), \qquad \eta = \dfrac{K_I\omega_c}{\bar{u} + rK_D\omega_c}, \\ k^{\pm} = \dfrac{\omega_c}{2\mu}\left(\bar{u} + rK_P \pm \sqrt{(\bar{u} + rK_P)^2 - 4\mu\dfrac{K_I}{\omega_c}(\bar{u} + K_D\omega_c r)}\right), \\ \beta^{\pm} = \dfrac{1}{k^{\pm}}(K_D\omega_c - K_P)\omega_c, \qquad \theta^{\pm} = \dfrac{\mu}{r} + \beta^{\pm}. \end{cases} \tag{S26}$$

Several observations can be made here. First, observe that two values of $k, \beta$ and $\theta$ can achieve the same PID gains and cutoff frequency. Furthermore, observe that the designed controller parameters depend on the plant at hand (via the supporting input $\bar{u}$). This is the price one has to pay for the simplicity in the design of the second order *a*PID controller. Finally, since the biomolecular parameters $k$ and $\beta$ cannot be negative nor complex numbers, the achievable PID gains and cutoff frequency are constrained.

**PID Coverage:** Constraining $k$ and $\beta$ to be real and non-negative yields the following achievable PID gains and cutoff frequency.

$$\mathcal{S}_2 = \left\{ (K_P, K_I, K_D, \omega_c) \in \mathbb{R}^4 : \; K_P \leq K_D\omega_c, \; 0 \leq K_I \leq \frac{\omega_c}{4\mu} \frac{(\bar{u} + rK_P)^2}{\bar{u} + rK_D\omega_c}, \; K_D, \omega_c \geq 0 \right\}. \tag{S27}$$

This means that the second order *a*PID controller cannot achieve a given $(K_P, K_I, K_D, \omega_c) \notin \mathcal{S}_2$ which, potentially, may put a limitation on the overall achievable performance. Once again, this is the price of the simplicity and minimality of the controller design.

#### S4.2 Third Order *a*PID Feedback Controller

Consider an arbitrary plant satisfying Assumptions 1 and 2 controlled with a third order *a*PID controller with degradation inhibition of $\mathbf{X_1}$ and additive inhibition of $\mathbf{Z_3}$ (see Figure S3(a)). The closed loop dynamics are thus given by

$$\begin{cases} \dot{x} = S\lambda(x) + ue_1; \qquad u = kz_1 - (\delta x_L + \delta_0 z_3)\dfrac{x_1}{x_1 + \kappa_1} \\ \dot{z}_1 = \mu - \eta z_1 z_2 \\ \dot{z}_2 = \theta x_L - \eta z_1 z_2 \\ \dot{z}_3 = \dfrac{\alpha_0}{1 + (x_L/\kappa_0)^{n_0}} - \gamma_0 z_3. \end{cases} \tag{S28}$$

The set-point is given by $\bar{x}_L := r = \mu/\theta$. For a given plant and set-point $r$, the supporting input $\bar{u}$ and input species concentration $\bar{x}_1$ are both fixed (see Assumptions 1 and 2). We first treat the analysis problem, then move on to the design problem. In what follows, we consider large $\eta$ so that the low pass filter with cutoff frequency $\omega_c \to \infty$ in Figure S3(b) becomes the identity map and thus can be ignored. As a result, only one low pass filter with cutoff frequency $\omega_0$ remains.

**Analysis:** The controller coordinates $(\bar{z}_1, \bar{z}_2, \bar{z}_3)$ of the fixed point are given by

$$\begin{cases} \bar{z}_1 = \dfrac{1}{k}\left[\bar{u} + \left(\delta r + \dfrac{\delta_0}{\gamma_0}\dfrac{\alpha_0}{1 + (r/\kappa_0)^{n_0}}\right) T(\bar{x}_1)\right] \\ \bar{z}_2 = \dfrac{\mu}{\eta \bar{z}_1} \underset{\eta \to \infty}{\longrightarrow} 0 \\ \bar{z}_3 = \dfrac{1}{\gamma_0}\dfrac{\alpha_0}{1 + (r/\kappa_0)^{n_0}}, \end{cases} \tag{S29}$$

where $T(\bar{x}_1) := \frac{\bar{x}_1}{\bar{x}_1 + \kappa_1} \approx 1$ (by choosing $\kappa_1$ to be small enough). Clearly, $\bar{z}_1, \bar{z}_2, \bar{z}_3 \geq 0$ when $\bar{u} \in \mathcal{U} = \mathbb{R}_+$. Hence, using the formulas derived in (S6), one can write the PID gains $(K_P, K_I, K_D)$ and cutoff frequency $\omega_0$ in terms of the various biomolecular parameters as

$$K_I = k\frac{\bar{z}_1}{\bar{z}_1 + \bar{z}_2} \underset{\eta \to \infty}{\longrightarrow} k, \quad K_P = \left(\delta - \frac{\delta_0 \alpha_0}{\gamma_0}\frac{n_0}{r}\frac{(r/\kappa_0)^{n_0}}{[1 + (r/\kappa_0)^{n_0}]^2}\right) T(\bar{x}_1), \quad K_D = \frac{\delta}{\gamma_0} T(\bar{x}_1), \quad \omega_0 = \gamma_0. \tag{S30}$$

Clearly for this *a*PID controller, the gains and cutoff frequency are independent of the plant for large $\eta$ and small $\kappa_1$. This is a clear advantage over the second-order *a*PID controller, but it comes at the price of more complexity (one additional controller species). However, observe that $\delta$ simultaneously tunes both $K_P$ and $K_D$ gains. This is a consequence of the incoherent feedforward loop that realizes an inseparable PD component.

**Design:** By fixing $\kappa_1, \kappa_0, n_0, \alpha_0, \mu$ and $r$ (and thus $\bar{u}$ and $\bar{x}_1$) and setting $\eta$ to be large enough, one can solve the nonlinear algebraic equations given in (S30) for the biomolecular parameters $\delta, k, \delta_0, \gamma_0$ and $\theta$ in terms of the PID gains and cutoff frequency. The results are provided here

$$\begin{cases} \delta = K_D\omega_0\left(1+\kappa_1/\bar{x}_1\right), \qquad k = K_I, \qquad \gamma_0 = \omega_0 \\ \delta_0 = \omega_0(K_D\omega_0 - K_P)\dfrac{r}{n_0\alpha_0}\left(1+\kappa_1/\bar{x}_1\right)\dfrac{\left[1+(r/\kappa_0)^{n_0}\right]^2}{(r/\kappa_0)^{n_0}}, \qquad \theta = \dfrac{\mu}{r}. \end{cases} \tag{S31}$$

Two observations can be made here. First, observe that the designed biomolecular controller parameters are independent of the plant at hand (once $\kappa_1$ is designed to be small enough). Furthermore, since the biochemical parameter $\delta_0$ cannot be negative, the achievable PID gains and cutoff frequency are constrained.

**PID Coverage:** Constraining $\delta_0$ to be non-negative yields the following achievable PID gains and cutoff frequency.

$$\mathcal{S}_3 = \left\{(K_P, K_I, K_D, \omega_0) \in \mathbb{R}^4 : \ K_P \leq K_D\omega_0, \ K_I, K_D, \omega_0 \geq 0\right\}. \tag{S32}$$

This means that the third order $a$PID controller cannot achieve a given $(K_P, K_I, K_D, \omega_0) \notin \mathcal{S}_3$. Furthermore, it is immediate to see that $\mathcal{S}_2 \subset \mathcal{S}_3$. As a result, one can reasonably expect that the third order $a$PID controller is capable of achieving better performance than the second order counterpart.

##### S4.3 Fourth Order *a*PID Feedback Controller

Consider an arbitrary plant satisfying Assumptions 1 and 2 controlled with a fourth order $a$PID controller with the derivative component entering the dynamics as a degradation reaction(see Figure S5). The closed-loop dynamics are thus given by

$$\begin{cases} \dot{x} = S\lambda(x) + ue_1; \qquad u = kz_1 - (\delta x_L + u_D)\dfrac{x_1}{x_1+\kappa_1}; \qquad u_D = \alpha_1 z_3 + \alpha_2 x_L \\ \dot{z}_1 = \mu - \eta z_1 z_2 \\ \dot{z}_2 = \theta x_L - \eta z_1 z_2 \\ \dot{z}_3 = \mu_0 - \eta_0 z_3 z_4 \\ \dot{z}_4 = \theta_0 u_D - \eta_0 z_3 z_4. \end{cases} \tag{S33}$$

The set-point is given by $\bar{x}_L := r = \mu/\theta$. For a given plant and set-point $r$, the supporting input $\bar{u}$ and input species concentration $\bar{x}_1$ are both fixed (see Assumptions 1 and 2). We first treat the analysis problem, then move on to the design problem. In what follows, we consider large $\eta$ and $\eta_0$ so that the low pass filter with cutoff frequency $\omega_c \to \infty$ in Figure S4(b) becomes the identity map and thus can be ignored. As a result, only one low pass filter with cutoff frequency $\omega_0$ remains.

**Analysis:** The controller coordinates $(\bar{z}_1, \bar{z}_2, \bar{z}_3, \bar{z}_4)$ of the fixed point are given by

$$\begin{cases} \bar{z}_1 = \dfrac{1}{k}\left[\bar{u} + (\delta r + \mu_0)\,\sigma_D\right] \\ \bar{z}_2 = \dfrac{\mu}{\eta\bar{z}_1} \underset{\eta\to\infty}{\longrightarrow} 0 \\ \bar{z}_3 = \dfrac{\mu_0 - \alpha_2 r}{\alpha_1} \\ \bar{z}_4 = \dfrac{\mu_0}{\eta_0\bar{z}_3} \underset{\eta_0\to\infty}{\longrightarrow} 0, \end{cases} \tag{S34}$$

where $\sigma_D := T(x_1) = \frac{\bar{x}_1}{\bar{x}_1+\kappa_1} \approx 1$ (by choosing $\kappa_1$ to be small enough). In what follows, $\mu_0$ is thought to be as design parameter that freely places the $z_3$-coordinate of the fixed point. That is, given $\alpha_1, \alpha_2, r$ and a desired $\bar{z}_3 > 0$, $\mu_0$ is designed to be

$$\mu_0 = \alpha_1\bar{z}_3 + \alpha_2 r.$$

As a result, $\bar{z}_1, \bar{z}_2, \bar{z}_3, \bar{z}_4 \geq 0$ when $\bar{u} \in \mathcal{U} = \mathbb{R}_+$. Hence, using the formulas derived in (S8), one can write the PID gains $(K_P, K_I, K_D)$ and cutoff frequency $\omega_0$ in terms of the various biomolecular parameters as

$$K_I = k\frac{\bar{z}_1}{\bar{z}_1+\bar{z}_2} \underset{\eta\to\infty}{\longrightarrow} k, \quad K_P = \delta T(\bar{x}_1), \quad K_D = \frac{T(\bar{x}_1)}{\theta_0}\frac{\alpha_2}{\alpha_1}, \quad \omega_0 = \theta_0\alpha_1. \tag{S35}$$

Clearly for this $a$PID controller, the gains and cutoff frequency are independent of the plant for large $\eta$ and small $\kappa_1$.

**Design:** By fixing $\theta_0, \kappa_1, \bar{z}_3$ and $r$ (and thus $\bar{u}$ and $\bar{x}_1$) and setting $\eta$ and $\eta_0$ to be large enough, one can solve the nonlinear algebraic equations given in (S35) for the biomolecular parameters $\delta, k, \alpha_1, \alpha_2, \mu_0$ and $\theta$ in terms of the PID gains and cutoff frequency. The results are provided here

$$\delta = \frac{K_P}{T(\bar{x}_1)}, \quad k = K_I, \quad \alpha_1 = \frac{\omega_0}{\theta_0}, \quad \alpha_2 = \frac{K_D \omega_0}{T(\bar{x}_1)}, \quad \theta = \frac{\mu}{r}. \tag{S36}$$

Two observations can be made here. First, observe that the designed controller biochemical parameters are independent of the plant at hand (once $\kappa_1$ is designed to be small enough then $T(\bar{x}_1) \approx 1$). Furthermore, observe that the three PID gains can be tuned separately with different biomolecular controller parameters. This is a consequence of separable P, I, and D components.

**PID Coverage:** The achievable PID gains and cutoff frequency are given by

$$\mathcal{S}_4 = \left\{ (K_P, K_I, K_D, \omega_0) \in \mathbb{R}^4 : \; K_P, K_I, K_D, \omega_0 \geq 0 \right\}. \tag{S37}$$

This means that the fourth order $a$PID controller can achieve any positive $(K_P, K_I, K_D, \omega_0)$. Furthermore, it is immediate to see that, over the positive orthant $\mathbb{R}^4_+$, $\mathcal{S}_4 \subset \mathcal{S}_3 \subset \mathcal{S}_2$. However, the second and third order $a$PID controllers can achieve negative $K_P$ gains while this fourth order $a$PID controller cannot. Nevertheless, this can be easily remedied by using a hybrid design where a P-Type proportional controller is mixed with an N-Type Integral and Derivative controllers (see for example Figure 7(d) in the main text).

### S5 $a$PID Control of Gene Expression

Consider a plant comprised of a simple gene expression network depicted in Figure 6(a). The deterministic dynamics of the plant are governed by the following two linear ordinary differential equations

$$\begin{cases} \dot{x}_1 = u - \gamma_1 x_1 \\ \dot{x}_2 = k_1 x_1 - \gamma_2 x_2, \end{cases} \tag{S38}$$

where the input $u$ is the total actuation propensity, $\mathbf{X_1}$ is the input species, and $\mathbf{X_2}$ is the output species. The transfer function of the plant is thus given by

$$P(s) = \frac{k_1}{(s + \gamma_1)(s + \gamma_2)}. \tag{S39}$$

We first show that a PI controller alone exhibits a fundamental limitation on the performance enhancement. Then we show that adding the derivative component adds more flexibility in the case of a second order $a$PID controller; whereas, it adds arbitrary flexibility in the case of a third and fourth order $a$PID controllers.

#### S5.1 Limitation of an $a$PI Controller

Consider the gene expression network controlled by an $a$PI controller of Class 1 with degradation inhibition depicted in Figure S7(a). To obtain a pure PI controller, we consider a large $\eta \to \infty$ which will make $\omega_c \to \infty$ and reduce the low pass filter in Figure S1 to the identity map and $K_F \approx 0$ as depicted in Figure S7(b). Hence it is straightforward to write down the closed-loop transfer function from $\tilde{\mu}$ to $\tilde{x}_L$ given by

$$H_1(s) := \frac{\tilde{x}_L(s)}{\tilde{\mu}(s)} = \frac{K_I P(s)}{s + [K_I K_S + K_P s] \, P(s)}. \tag{S40}$$

To do a root locus analysis in $K_P$, we rewrite $H_1(s)$ in a root-locus standard form given by

$$H_1(s) = \frac{K_I P(s) / \left(s + K_I K_S P(s)\right)}{1 + K_P \frac{s P(s)}{s + K_I K_S P(s)}}. \tag{S41}$$

By substituting the plant transfer function given in (S39), the denominator of $H_1(s)$ can be written as

$$D(s) = 1 + K_P G(s); \qquad \text{with} \qquad G(s) := \frac{k_1 s}{s^3 + (\gamma_1 + \gamma_2)s^2 + \gamma_1 \gamma_2 s + k_1 K_I K_S}. \tag{S42}$$

Note that when $K_P = 0$, the closed-loop characteristic polynomial is given by

$$D_I(s) = s^3 + (\gamma_1 + \gamma_2)s^2 + \gamma_1 \gamma_2 s + k_1 K_I K_S. \tag{S43}$$

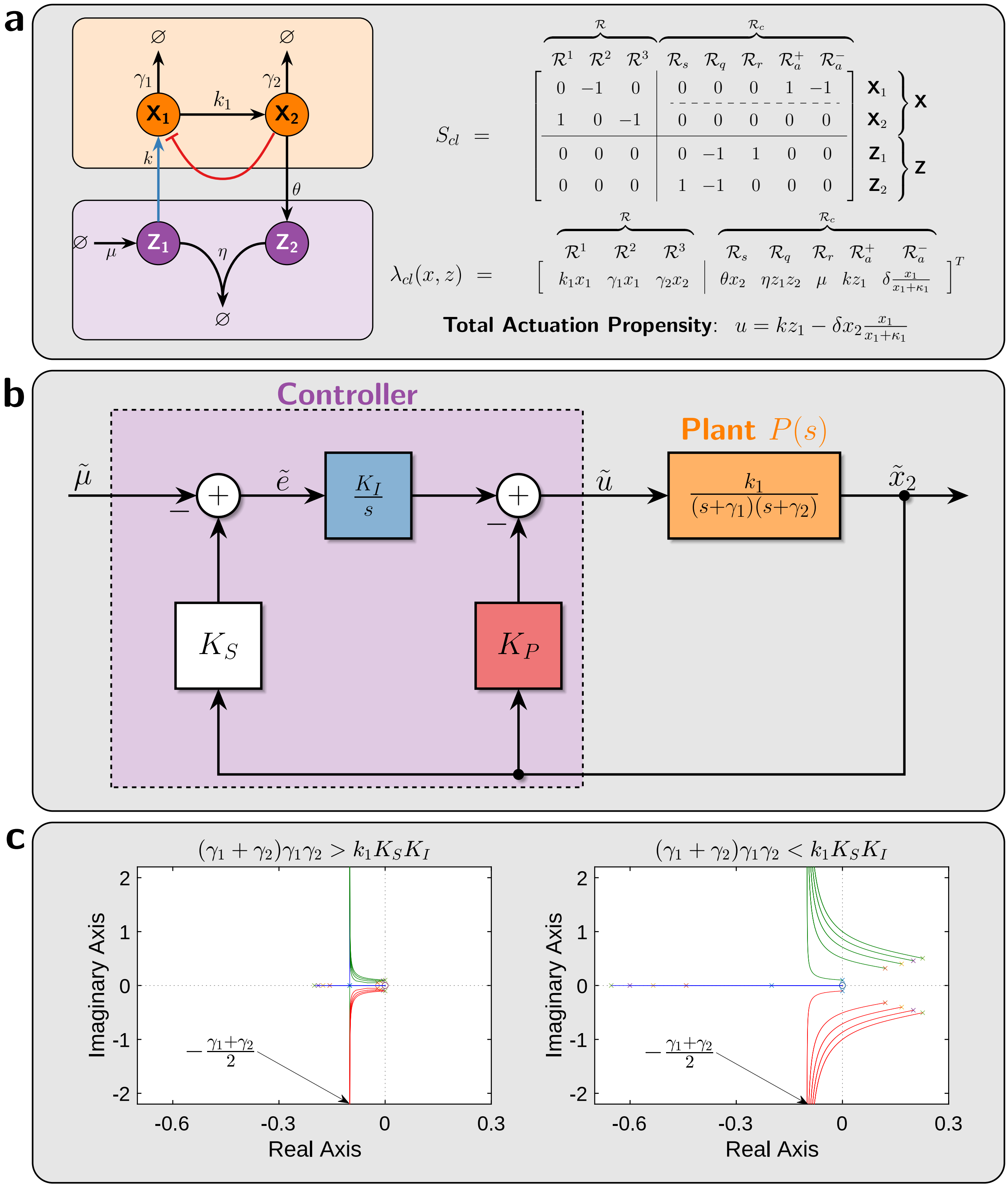

Figure S7: **Root Locus in $K_P$ for Different Integral Gains $K_I$. (a) Gene Expression Network.** The plant input, denoted by $u$, is the total actuation propensity that takes into account both positive and negative actuation. **(b)** Block Diagram of the Linearized Dynamics. This block diagram is a particular instance of the general block diagram given in Figure S1 when the plant is gene expression, $\eta$ is large enough and $\kappa_1$ is small enough. **(c) Root Locus in $K_P$ for multiple values of the integral gain $K_I$.**

This polynomial has all roots lying in the left-half complex plane iff $(\gamma_1 + \gamma_2)\gamma_1\gamma_2 < k_1 K_S K_I$. For the root locus analysis, two cases can be considered.

In the first case, we have $(\gamma_1 + \gamma_2)\gamma_1\gamma_2 \leq k_1 K_S K_I$. Hence for $K_P = 0$, there are two complex poles in the right-half plane (Descartes rule of signs rules out positive real poles), and one negative pole on the real axis. As $K_P$ increases a simple root locus argument shows that the negative real pole approaches the zero at the origin, while the complex unstable poles cross the imaginary axis and diverge vertically to + infinity and - infinity at an angle of $+\pi/2$ and $-\pi/2$, respectively. It is fairly straightforward to compute the abscissa of the vertical asymptote $\sigma_A = -(\gamma_1+\gamma_2)/2$ which turns out to be independent of the integral gain $K_I$. The resulting root locus is illustrated in the left plot of Figure S7(c) for multiple values of $K_I$.

For the second case, we have $(\gamma_1+\gamma_2)\gamma_1\gamma_2 > k_1 K_S K_I$. Hence for $K_P = 0$, there are either three negative real poles on the real axis, or there are two complex poles and one real pole in the left half of the complex plane. Either way, a simple root locus argument shows that, as $K_P$ increases, one real pole approaches the zero at the origin, while the other two poles approach the vertical asymptote, at $\sigma_A = -(\gamma_1+\gamma_2)/2$ and diverge to + infinity and - infinity at an angle of $+\pi/2$ and $-\pi/2$, respectively. Once again, this vertical asymptote is independent of the integral gain $K_I$. The resulting root locus is illustrated in the right plot of Figure S7(c) for multiple values of $K_I$.

In conclusion, for any integral and proportional gains $K_I$ and $K_P$, we cannot place all the poles to the left of the vertical asymptote at $\sigma_A = -(\gamma_1+\gamma_2)/2$. Hence the growth/decay rate of the response cannot be made faster than the average of the two degradation rates of $\mathbf{X_1}$ and $\mathbf{X_2}$ without giving rise to oscillations. As a result, if $\gamma_1$ and $\gamma_2$ are small, the performance of the PI controller can be slow and unsatisfactory. This analytical result is demonstrated by the simulations shown in Figure 6(b).

#### S5.2 Flexibility of the *a*PID Controllers

Consider the gene expression network controlled by the second, third and fourth order $a$PID controllers. Using the block diagrams in Figures S2(b), S3(b) and S4(b), one can write down the closed-loop transfer functions $H_2(s), H_3(s)$ and $H_4(s)$ of the second, third and fourth order $a$PID controllers, respectively, as

$$\begin{aligned}
H_2(s) &= \frac{K_F k_1 \omega s + K_I k_1 \omega}{s^4 + (\gamma_1+\gamma_2+\omega)s^3 + \left[\omega(K_D k_1 + \gamma_1 + \gamma_2) + \gamma_1\gamma_2\right]s^2 + \omega(K_P k_1 + \gamma_1\gamma_2)s + k_1 K_I K_S \omega} \\
H_3(s) &= \frac{K_I k_1 s + K_I k_1 \omega}{s^4 + (\gamma_1+\gamma_2+\omega)s^3 + \left[\omega(K_D k_1 + \gamma_1 + \gamma_2) + \gamma_1\gamma_2\right]s^2 + \left[K_I K_S k_1 + \omega(K_P k_1 + \gamma_1\gamma_2)\right]s + k_1 K_I K_S \omega} \\
H_4(s) &= \frac{K_I k_1 s + K_I k_1 \omega}{s^4 + (\gamma_1+\gamma_2+\omega)s^3 + \left[\omega(K_D k_1 + \gamma_1 + \gamma_2) + K_P k_1 + \gamma_1\gamma_2\right]s^2 + \left[K_I K_S k_1 + \omega(K_P k_1 + \gamma_1\gamma_2)\right]s + k_1 K_I K_S \omega},
\end{aligned} \tag{S44}$$

where the indeces of $\omega$ are dropped for notational convenience, $\eta_0$ is chosen to be large and $\eta$ is chosen to be large only for the third and fourth order $a$PID controllers. Consider the pole placement design problem, where one aims at designing the values of the PID gains and the cutoff frequency to place the four closed-loop poles at $s = -a$. Ideally, we aim at placing the poles in the left-half complex plane to guarantee stability and on the real axis to avoid oscillations. Furthermore, to obtain a fast response, the poles should be placed far to the left in the left-half complex plane. These criteria can all be achieved if we make $a > 0$ and large enough. Next, we study if this is possible to achieve with the three $a$PID controllers. Since the four closed-loop poles are place at $s = -a$, then the closed-loop characteristic polynomial is given by

$$p(s) = (s+a)^4 = s^4 + 4as^3 + 6a^2s^2 + 4a^3s + a^4. \tag{S45}$$

Equating $p(s)$ to the denominators of $H_2(s), H_3(s)$ and $H_4(s)$ allows us to express the designed PID gains and cutoff frequency in terms of the plant parameters $(\gamma_1, \gamma_2, k_1)$, the sensing gain $K_S$ and the placed pole $-a$ as

$$\begin{aligned}
&\textbf{Second Order:} \begin{cases} K_P = \dfrac{4a^3}{k_1[4a-(\gamma_1+\gamma_2)]} - \dfrac{\gamma_1\gamma_2}{k_1}, & K_I = \dfrac{a^4}{K_S k_1[4a-(\gamma_1+\gamma_2)]}, \\ K_D = \dfrac{6a^2-\gamma_1\gamma_2}{k_1[4a-(\gamma_1+\gamma_2)]} - \dfrac{\gamma_1+\gamma_2}{k_1}, & \omega = 4a-(\gamma_1+\gamma_2) \end{cases} \\
&\textbf{Third Order:} \begin{cases} K_P = \dfrac{a^3[15a-4(\gamma_1+\gamma_2)]}{k_1[4a-(\gamma_1+\gamma_2)]^2} - \dfrac{\gamma_1\gamma_2}{k_1}, & K_I = \dfrac{a^4}{K_S k_1[4a-(\gamma_1+\gamma_2)]}, \\ K_D = \dfrac{6a^2-\gamma_1\gamma_2}{k_1[4a-(\gamma_1+\gamma_2)]} - \dfrac{\gamma_1+\gamma_2}{k_1}, & \omega = 4a-(\gamma_1+\gamma_2) \end{cases} \\
&\textbf{Fourth Order:} \begin{cases} K_P = \dfrac{a^3[15a-4(\gamma_1+\gamma_2)]}{k_1[4a-(\gamma_1+\gamma_2)]^2} - \dfrac{\gamma_1\gamma_2}{k_1}, & K_I = \dfrac{a^4}{K_S k_1[4a-(\gamma_1+\gamma_2)]}, \\ K_D = \dfrac{(\gamma_1+\gamma_2-3a)^4}{k_1[4a-(\gamma_1+\gamma_2)]^3}, & \omega = 4a-(\gamma_1+\gamma_2). \end{cases}
\end{aligned} \tag{S46}$$

Recall, that the sets of achievable PID gains and cutoff frequencies for the second, third and fourth order $a$PID controllers are given by $\mathcal{S}_2, \mathcal{S}_3$ and $\mathcal{S}_4$ in (S27), (S32) and (S37), respectively. These sets constrain the achievable poles

$s = -a$ to the following regions on the real axis.

$$\begin{aligned}
&\textbf{Second Order:} && (K_P, K_I, K_D, \omega) \in \mathcal{S}_2 && \implies && \frac{\gamma_1+\gamma_2}{2} \le a \le (2+\sqrt{2})\left(\frac{\gamma_1+\gamma_2}{2}\right) \\
&\textbf{Third Order:} && (K_P, K_I, K_D, \omega) \in \mathcal{S}_3 && \implies && \frac{\gamma_1+\gamma_2}{4} \le a \\
&\textbf{FourthOrder:} && (K_P, K_I, K_D, \omega) \in \mathcal{S}_4 && \implies && \frac{\gamma_1+\gamma_2}{4} \le a.
\end{aligned} \tag{S47}$$

As a result, for the second order $a$PID controller, the fastest poles achievable are for $a = (2+\sqrt{2})\left(\frac{\gamma_1+\gamma_2}{2}\right)$; whereas, for the third and fourth order $a$PID controllers, there is no theoretical limit.

In conclusion, this case study shows that the second order $a$PID greatly improves the performance compared to only $a$PI controllers, but it is limited in the sense that the response cannot be made arbitrarily fast without giving rise to overshoots and/or oscillations. However, the third and fourth order $a$PID controllers are capable of making the response as fast as desired without giving rise to overshoots nor oscillations. These analytical results are demonstrated by the simulations shown in Figure 6(c).

### S6 Alternative Differentiators

The derivative operations of the second and third order $a$PID controllers are realized via incoherent feedforward loops. As for the fourth order $a$PID controller, the derivative operator that we refer to as *Antithetic Differentiator* is fundamentally different. It is realized by placing the antithetic integral motif in feedback with itself. This is a trick in control theory for implementing differentiators using integrators. Of course, the resulting differentiator is low pass filtered since a pure derivative cannot be realized physically: a pure derivative requires accessing future inputs. In this section, we show that this trick can be used to construct other differentiators by exploiting different integrators (other than the antithetic integrator).

The basic underlying idea is demonstrated in the block diagrams of Figure S8(a). A straightforward algebraic calculation shows that the filtered derivative operation acting on the input $u$ in the left block diagram of Figure S8(a) can be exactly realized by placing an integrator (blue block) in the feedback path as depicted in the right block diagram. In fact, for the right block diagram, we have

$$y(s) = \omega_0 K_D \left(u(s) - \frac{y(s)}{K_D s}\right) \quad \implies \quad \left(1 + \frac{\omega_0}{s}\right) y(s) = \omega_0 K_D u(s) \quad \implies \quad \frac{y(s)}{u(s)} = K_D s \frac{\omega_0}{s+\omega_0} = H(s).$$

As a result, the input/output relationships for both block diagrams are exactly the same. This particular idea motivated the design of the antithetic differentiator using the antithetic integral motif. Nevertheless, one can use other integrators such as the inflow/outflow zero-order integrators depicted in Figure S8(b) and auto-catalytic integrator depicted in Figure S8(c) to design other differentiators as well.

#### S6.1 Outflow Zero-Order Differentiator

Consider the outflow zero-order differentiator depicted in Figure S8(b), where the dynamics are shown as a single differential equation and an output algebraic equation. Let $(\tilde{u}, \tilde{z}, \tilde{y})$ denote the perturbation from the operating point $(\bar{u}, \bar{z}, \bar{y})$ with

$$\mu_0 - \theta_0 \bar{y} \frac{\bar{z}}{\bar{z} + \kappa_0} = 0 \qquad \text{and} \qquad \bar{y} = g(\bar{z}, \bar{u}),$$

where $g$ is a monotonically increasing function in $z$ and $u$.

**Requirement:** For this circuit to function as a differentiator, $\kappa_0$ needs to be small $\kappa_0 << z$. Otherwise, it can be shown that this circuit will realize a filtered PD instead of a filtered D which can be leveraged if need be.

Under this condition, we have $\frac{z}{z+\kappa_0} \approx 1$ and thus the dynamics can be approximated as

$$\begin{cases} \dot{z} = \mu_0 - \theta_0 y \\ y = g(z, u), \end{cases}$$

and $\bar{y} = \mu_0/\theta_0$ is independent of the input $\bar{u}$ at steady-state. This represents the offset value of the computed derivative, i.e. the computed derivative is lifted by a positive constant equal to $\mu_0/\theta_0$. To carry out a linear perturbation analysis we consider the dynamics of the perturbation variables given by

$$\tilde{u}(t) = u(t) - \bar{u}; \quad \tilde{z}(t) = z(t) - \bar{z}; \quad \tilde{y}(t) = y(t) - \bar{y}.$$

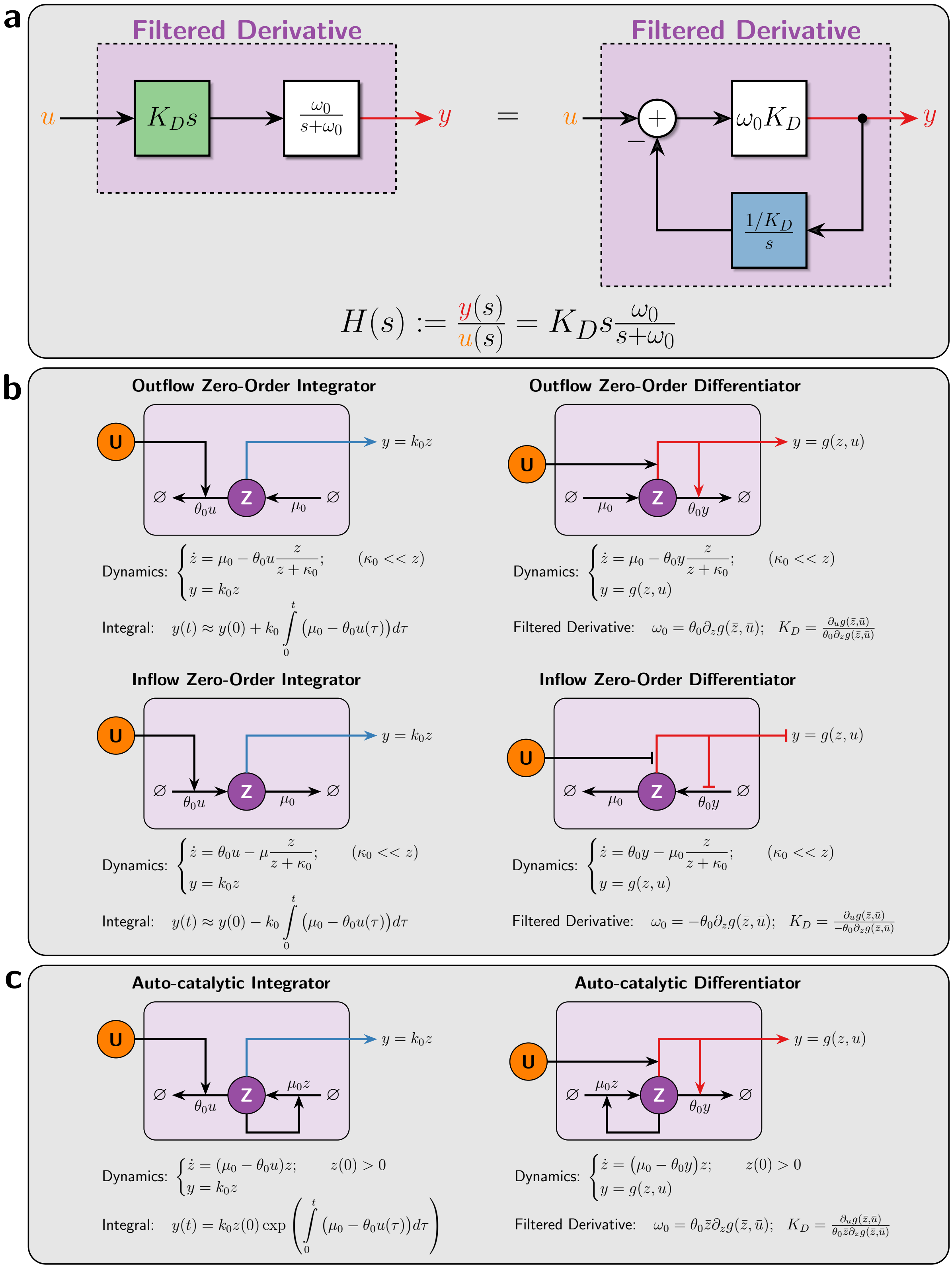

Figure S8: **Alternative Differentiators. (a) Implementation of Differentiators using Integrators.** The block diagram to the left depicts the basic filtered derivative operation, where $u$ is the input and $y$ is the output. The derivative gain is denoted by $K_D$ and the cutoff frequency of the low pass filter is denoted by $\omega_0$. This operation can be realized by placing an integrator in the feedback path from the output $y$. A straight forward computation shows that both block diagrams have the same exact transfer function denoted by $H(s)$. **(b) and (c) Design of Differentiators using Zero-Order and Auto-Catalytic Integrators.** The input is the concentration of **U**, denoted by lowercase $u$; whereas, the output, denoted by $y$, is a rate. By exploiting the idea of **(a)**, one can use the various integrators to design differentators with gain $K_D$ and cutoff frequency $\omega_0$.

The approximate linearized dynamics can thus be written as

$$\begin{cases} \dot{\tilde{z}} = -\theta_0 \tilde{y} \\ \tilde{y} = \sigma_z \tilde{z} + \sigma_u \tilde{u} \end{cases}$$

where $\partial g(\bar{z}, \bar{u}) =: \begin{bmatrix} \sigma_z & \sigma_u \end{bmatrix}$ such that $\sigma_z, \sigma_u > 0$. Taking the Laplace transforms yields

$$\begin{cases} \tilde{z}(s) = -\theta_0 \dfrac{\tilde{y}(s)}{s} \\ \tilde{y}(s) = \sigma_z \tilde{z}(s) + \sigma_u \tilde{u}(s) \end{cases} \implies \frac{\tilde{y}(s)}{\tilde{u}(s)} = K_D s \frac{\omega_0}{s + \omega_0} \quad \text{with} \quad \begin{cases} K_D := \dfrac{1}{\theta_0} \dfrac{\sigma_u}{\sigma_z} \\ \omega_0 := \theta_0 \sigma_z. \end{cases}$$

This analysis shows indeed that this circuit mathematically realizes a P-Type (positive) filtered derivative with gain $K_D$ and cutoff frequency $\omega_0$ as depicted in Figure S8(b). Note that an N-Type version (negative derivative) of this circuit can be simply achieved by requiring $g$ to be a monotonically decreasing function in $u$ instead as depicted in Figure 12.

#### S6.2 Inflow Zero-Order Differentiator

Consider the inflow zero-order differentiator depicted in Figure S8(b), where the dynamics are shown as a single differential equation and an output algebraic equation. Let $(\tilde{u}, \tilde{z}, \tilde{y})$ denote the perturbation from the operating point $(\bar{u}, \bar{z}, \bar{y})$ with

$$\theta_0 \bar{y} - \mu_0 \frac{\bar{z}}{\bar{z} + \kappa_0} = 0 \qquad \text{and} \qquad \bar{y} = g(\bar{z}, \bar{u}),$$

where $g$ is a monotonically increasing (resp. decreasing) function in $u$ (resp. $z$).

**Requirement:** For this circuit to function as a differentiator, $\kappa_0$ needs to be small $\kappa_0 << z$. Otherwise, it can be shown that this circuit will realize a filtered PD instead of a filtered D which can be leveraged if need be.

Under this condition, we have $\frac{z}{z+\kappa_0} \approx 1$ and thus the dynamics can be approximated as

$$\begin{cases} \dot{z} = \theta_0 y - \mu_0 \\ y = g(z, u), \end{cases}$$

and $\bar{y} = \mu_0/\theta_0$ is independent of the input $\bar{u}$ at steady-state. This represents the offset value of the computed derivative, i.e. the computed derivative is lifted by a positive constant equal to $\mu_0/\theta_0$. To carry out a linear perturbation analysis we consider the dynamics of the perturbation variables given by

$$\tilde{u}(t) = u(t) - \bar{u}; \quad \tilde{z}(t) = z(t) - \bar{z}; \quad \tilde{y}(t) = y(t) - \bar{y}.$$

The approximate linearized dynamics can thus be written as

$$\begin{cases} \dot{\tilde{z}} = \theta_0 \tilde{y} \\ \tilde{y} = -\sigma_z \tilde{z} + \sigma_u \tilde{u} \end{cases}$$

where $\partial g(\bar{z}, \bar{u}) =: \begin{bmatrix} -\sigma_z & \sigma_u \end{bmatrix}$ such that $\sigma_z, \sigma_u > 0$. Taking the Laplace transforms yields

$$\begin{cases} \tilde{z}(s) = \theta_0 \dfrac{\tilde{y}(s)}{s} \\ \tilde{y}(s) = -\sigma_z \tilde{z}(s) + \sigma_u \tilde{u}(s) \end{cases} \implies \frac{\tilde{y}(s)}{\tilde{u}(s)} = K_D s \frac{\omega_0}{s + \omega_0} \quad \text{with} \quad \begin{cases} K_D := \dfrac{1}{\theta_0} \dfrac{\sigma_u}{\sigma_z} \\ \omega_0 := \theta_0 \sigma_z. \end{cases}$$

This analysis shows indeed that this circuit mathematically realizes a P-Type (positive) filtered derivative with gain $K_D$ and cutoff frequency $\omega_0$ as depicted in Figure S8(b). Note that an N-Type version (negative derivative) of this circuit can be simply achieved by requiring $g$ to be a monotonically decreasing function in $u$ instead as depicted in Figure 12.

#### S6.3 Auto-Catalytic Differentiator

Consider the auto-catalytic differentiator depicted in Figure S8(c), where the dynamics are shown as a single differential equation and an output algebraic equation. Let $(\tilde{u}, \tilde{z}, \tilde{y})$ denote the perturbation from the operating point $(\bar{u}, \bar{z}, \bar{y})$ with

$$(\mu_0 - \theta_0 \bar{y})\, \bar{z} = 0 \qquad \text{and} \qquad \bar{y} = g(\bar{z}, \bar{u}),$$

where $g$ is a monotonically increasing function in $u$ and $z$. The dynamics have two fixed points given by

$$\bar{z}_1 = 0 \qquad \text{and} \qquad \bar{z}_2 \text{ satisfying } g(\bar{z}_2, \bar{u}) = \mu_0/\theta_0,$$

for a given steady-state value $\bar{u} > 0$ of the input. The corresponding outputs are given by

$$\bar{y}_1 = g(0, \bar{u}) \qquad \text{and} \qquad \bar{y}_2 = \mu_0/\theta_0.$$

$\bar{y}_1$ and $\bar{y}_2$ represent the offset value of the computed derivative which is required to be strictly positive; otherwise, the differentiator cannot compute the derivatives of decreasing inputs since the output $y$ is a rate that cannot be negative. This requirement is automatically satisfied if the only stable fixed point is $\bar{z}_2 > 0$, satisfying $g(\bar{z}_2, \bar{u}) = \mu_0/\theta_0$, which implies that the offset is given by $\bar{y}_2 = \mu_0/\theta_0 > 0$. In contrast, if the origin is the only locally stable fixed point, then the offset is given by $\bar{y}_1 = g(0, \bar{u})$. This imposes the requirement that $g$ should be chosen such that $g(0, \bar{u}) > 0$.

Let's analyze the local stability around the two possible fixed points. Let $f(z) := [\mu_0 - \theta_0 g(z, u)]z$ denote the right-hand side of the differential equation. Hence the derivative of $f$ is given by $f'(z) = \mu_0 - \theta_0 g(z, u) - \theta_0 \partial_z g(z, u) z$. Evaluating the derivative at the two fixed points $\bar{z}_1$ and $\bar{z}_2$ for a given $\bar{u} > 0$ yields

$$f'(\bar{z}_1) = \mu_0 - \theta_0 g(0, \bar{u}) \qquad \text{and} \qquad f'(\bar{z}_2) = -\theta_0 \partial_z g(\bar{z}_2, \bar{u}) \bar{z}_2.$$

Therefore, as long as the fixed point $\bar{z}_2$ is strictly positive, it is locally stable. The other fixed point $\bar{z}_1$ at the origin is locally stable only if $g(0, \bar{u}) > \mu_0/\theta_0$.

**Example 1:** If $g(z, u) = \eta_0 z u$ which is monotonically increasing in both $z$ and $u$ as required, then $\bar{z} = 0$ is an unstable fixed point while $\bar{z} = \frac{\mu_0}{\theta_0} \frac{1}{\eta_0 \bar{u}} > 0$ is a locally stable fixed point for any $\bar{u} > 0$. This means that the derivative offset is given by $\bar{y} = \mu_0/\theta_0 > 0$.

**Example 2:** If $g(z, u) = \alpha z + \beta u$ which is also monotonically increasing in both $z$ and $u$ as required, then $\bar{z} = 0$ is the only locally stable fixed point if $\bar{u} > \mu_0/\beta\alpha_0$ yielding a derivative offset of $\bar{y} = \beta \bar{u}$; whereas, $\bar{z} = \frac{1}{\alpha}\left(\frac{\mu_0}{\theta_0} - \beta \bar{u}\right)$ is the only locally stable fixed point if $\bar{u} < \mu_0/\beta\alpha_0$ yielding a derivative offset of $\bar{y} = \mu_0/\theta_0$.

Before we proceed, we point out that $g$ functions like Example 1 have a nice property: the offset is unconditionally strictly positive and is given by $\mu_0/\theta_0$ which is independent of the applied input $u$. In fact, we show next that $g$ functions like Example 2 have the tendency to fail in realizing a differentiator if $\bar{z} = 0$ is a locally stable fixed point. To carry out a linear perturbation analysis, we consider the dynamics of the perturbation variables given by

$$\tilde{u}(t) = u(t) - \bar{u}; \quad \tilde{z}(t) = z(t) - \bar{z}; \quad \tilde{y}(t) = y(t) - \bar{y}.$$

The approximate linearized dynamics can thus be written as

$$\begin{cases} \dot{\tilde{z}} = (\mu_0 - \theta_0 \bar{y})\tilde{z} - \theta_0 \bar{z} \tilde{y} \\ \tilde{y} = \sigma_z \tilde{z} + \sigma_u \tilde{u} \end{cases}$$

where $\partial g(\bar{z}, \bar{u}) =: \begin{bmatrix} \sigma_z & \sigma_u \end{bmatrix}$ such that $\sigma_z, \sigma_u > 0$. Observe that operating around the origin, i.e. $\bar{z} = 0$, destroys the feedback from $\tilde{y}$ to $\tilde{z}$, and results in a failure to realize a derivative operation. This imposes the requirement that the fixed point at the origin has to be unstable. This requirement is automatically and unconditionally satisfied for $g$ functions like Example 1; however, for $g$ functions like Example 2 this requirement is only satisfied for a certain range of inputs. As a result, we ask the following condition to be satisfied.

**Requirement:** The function $g$ must be designed such that the fixed point at the origin is unstable while the other fixed point $\bar{z}$ satisfying $g(\bar{z}, \bar{u}) = \mu_0/\theta_0$ is strictly positive for all inputs $\bar{u} > 0$. Example 1 provides a function $g$ that satisfies this requirement.

When this requirement is met, the output at steady state is given by $\bar{y} = \mu_0/\theta_0$. Taking the Laplace transforms of the linearized dynamics and substituting $\bar{y} = \mu_0/\theta_0$ yields

$$\begin{cases} \tilde{z}(s) = -\theta_0 \bar{z} \dfrac{\tilde{y}(s)}{s} \\ \tilde{y}(s) = \sigma_z \tilde{z}(s) + \sigma_u \tilde{u}(s) \end{cases} \implies \quad \frac{\tilde{y}(s)}{\tilde{u}(s)} = K_D s \frac{\omega_0}{s + \omega_0} \quad \text{with} \quad \begin{cases} K_D := \dfrac{1}{\theta_0 \bar{z}} \dfrac{\sigma_u}{\sigma_z} \\ \omega_0 := \theta_0 \sigma_z \bar{z}, \end{cases}$$

where $\bar{z}$ satisfies $g(\bar{z}, \bar{u}) = \mu_0/\theta_0$. This analysis shows indeed that this circuit mathematically realizes a P-Type (positive) filtered derivative with gain $K_D$ and cutoff frequency $\omega_0$ as depicted in Figure S8(c). Note that an N-Type version (negative derivative) of this circuit can be simply achieved by requiring $g$ to be a monotonically decreasing function in $u$ instead as depicted in Figure 12.

##### S6.4 Alternative PID Controllers

By appending the differentiators, introduced in this section, to the antithetic integral motif and the proportional control action, alternative PID controllers can be constructed as demonstrated in Figure 12. The inflow and outflow $a$PID controllers require a zeroth-order degradation reaction that can be realized via a saturated degradation mechanism $\kappa_0 << z_3$. The auto-catalytic $a$PID controller does not require such saturation; however, it requires that the fixed point with $\bar{z}_3 = 0$ to be unstable so that the dynamics converge to the other fixed point with $\bar{u}_D = g(\bar{z}_3, \bar{x}_L) = \mu_0/\theta_0$. We show next that the instability of $\bar{z}_3 = 0$ can be guaranteed by a suitable constraint on the function $g$.

For convenience, we rewrite the dynamics of the (N-Type) auto-catalytic $a$PID controller here (from Figure 12).

$$\begin{cases} \dot{x} = S\lambda(x) + h(z_1, x_1, x_L, u_D)e_1; \qquad u_D = g(z_3, x_L) \\ \dot{z}_1 = \mu - \eta z_1 z_2 \\ \dot{z}_2 = \theta x_L - \eta z_1 z_2 \\ \dot{z}_3 = (\mu_0 - \theta_0 u_D) z_3, \end{cases}$$

The jacobian of the dynamics around a fixed point $(\bar{x}, \bar{z}_1, \bar{z}_2, \bar{z}_3)$ is given by

$$\left[\begin{array}{cccc|ccc} & \vdots & & & \vdots & \vdots & \vdots \\ & S\partial\lambda(\bar{x}) - \sigma_3 e_1 e_1^T - (\sigma_4 + \sigma_D\sigma_6)e_1 e_L^T & & & \sigma_1 e_1 & 0 & \sigma_5 e_1 \\ & \vdots & & & \vdots & \vdots & \vdots \\ \hline 0 & \cdots & 0 & 0 & -\eta\bar{z}_2 & -\bar{z}_1 & 0 \\ 0 & \cdots & 0 & \theta & -\eta\bar{z}_2 & -\bar{z}_1 & 0 \\ 0 & \cdots & 0 & \theta_0\bar{z}_3\sigma_6 & 0 & 0 & (\mu_0 - \theta_0\bar{u}_D) - \theta_0\bar{z}_3\sigma_5 \end{array}\right],$$

where $\partial h(\bar{z}_1, \bar{x}_1, \bar{x}_L, \bar{u}_D) =: \begin{bmatrix}\sigma_1 & -\sigma_3 & \sigma_4 & \sigma_D\end{bmatrix}$ and $\partial g(\bar{z}_3, \bar{x}_L) =: \begin{bmatrix}\sigma_5 & -\sigma_6\end{bmatrix}$ with $\sigma_1, \sigma_3, \sigma_4, \sigma_6, \sigma_6, \sigma_D \geq 0$. Observe that if $\bar{z}_3 = 0$, then the last row of the Jacobian matrix becomes $\begin{bmatrix}0 & \cdots & 0 & 0 & 0 & 0 & (\mu_0 - \theta_0\bar{u}_D)\end{bmatrix}$ with $\bar{u}_D = g(0, \bar{x}_L)$. This implies that $\mu_0 - \theta_0 g(0, \bar{x}_L)$ is an eigenvalue of the Jacobian matrix. Clearly this eigenvalue is nonnegative for any set-point $\bar{x}_L$ if $g(0, x_L) = 0$. Therefore, the design requirement that $g(0, x_L) = 0$ guarantees that the origin is an unstable fixed point for any set-point $x_L$. One can show that this condition also is required for the P-Type auto-catalytic $a$PID controller as well.

Finally, before we close this section, we would like to point out that yet another set of PID controllers can also be obtained using different integral components as depicted in Figure S9.

#### S7 Stationary Variance Approximation for the $a$PI Controllers

Consider the closed-loop network depicted in Figure 4(a) where a gene expression network is connected in feedback with the $a$PI controllers of Class 1. The goal of this section is to derive an approximate formula for the stationary variance of the output species $\mathbf{X_2}$. First, we consider a general plant to write down the evolution equations of the variance. Then, we derive an approximate closed formula for the output stationary variance in the case of the particular gene expression plant given in Figure 4(a).

##### S7.1 Evolution of the Covariances for a General Plant

Let $X_{cl} := \begin{bmatrix} X \\ Z \end{bmatrix}$ denote the closed-loop state variable encrypting the copy numbers of the plant and controller species $\mathbf{X}$ and $\mathbf{Z}$, respectively. Define the instantaneous covariance of the closed-loop state variable as

$$\text{Cov}\left[X_{cl}(t)\right] := \mathbb{E}\left[\Big(X_{cl}(t) - \mathbb{E}\left[X_{cl}(t)\right]\Big)\Big(X_{cl}(t) - \mathbb{E}\left[X_{cl}(t)\right]\Big)^T\right],$$

whose evolution is described by the following differential equation (we drop the time variable for notational convenience)

$$\frac{d}{dt}\text{Cov}\left[X_{cl}\right] = S_{cl}\mathcal{D}\big(\mathbb{E}\left[\lambda_{cl}(X_{cl})\right]\big)S_{cl}^T + S_{cl}\text{Cov}\left[\lambda_{cl}(X_{cl}), X_{cl}\right] + \text{Cov}\left[X_{cl}, \lambda_{cl}(X_{cl})\right]S_{cl}^T, \tag{S48}$$

where $S_{cl}$ and $\lambda_{cl}$, depicted in Figure 3, denote the closed-loop stoichiometry matrix and propensity function, respectively. Note that $\mathcal{D}$ is the diagonal operator such that for any vector $x$, $\mathcal{D}(x)$ is a diagonal matrix whose diagonal

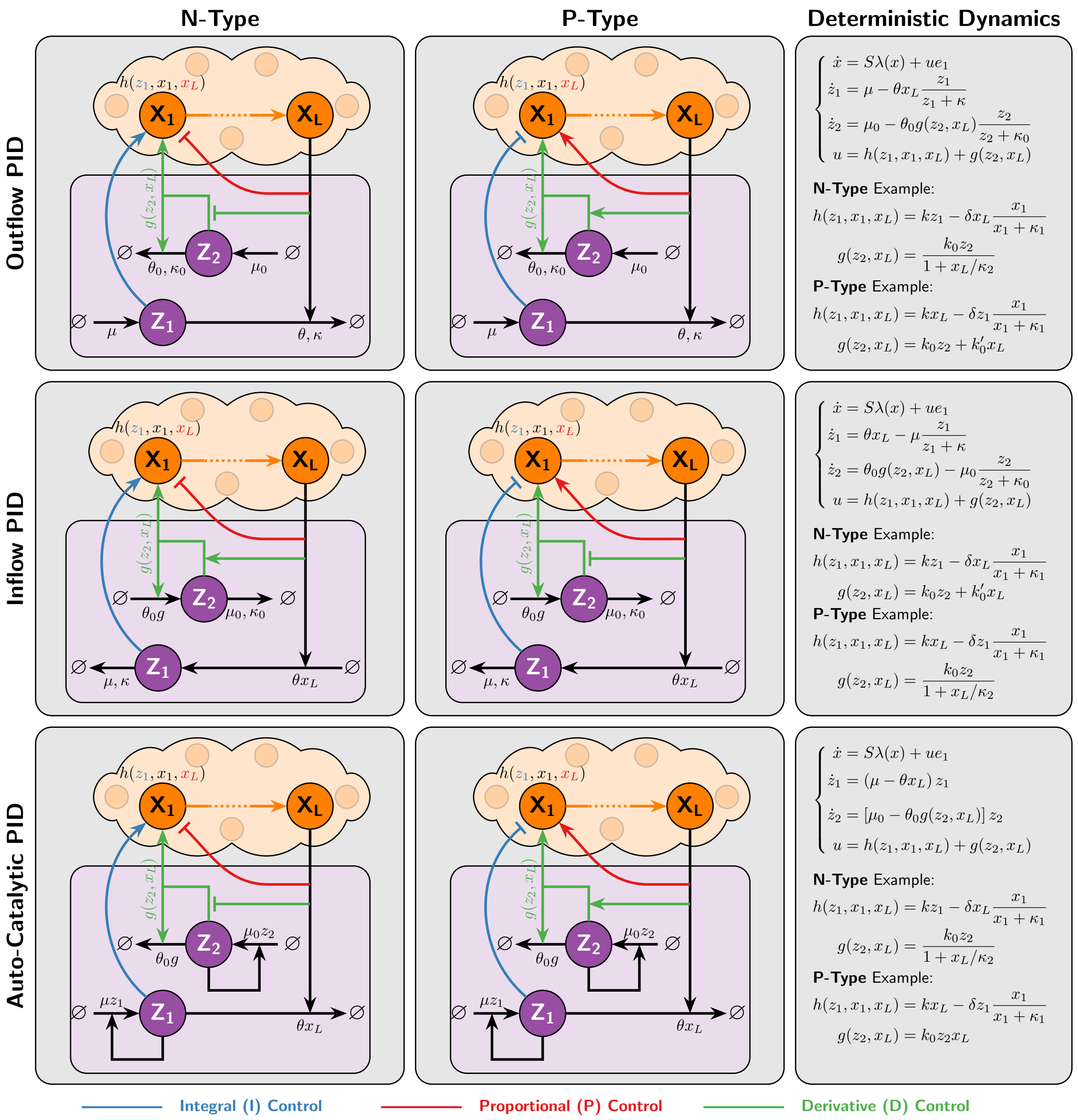

Figure S9: **Alternative PID Controllers.** This collection of controllers demonstrate how one can design various N- and P-Type PID controllers by exploiting integral components other than the antithetic integral motif. The controllers in the first two rows are realized via zeroth-order degradation reactions while the controllers in the last row are realized via auto-catalytic reactions.

entries are $x$. Define the matrices and vector

$$S_m := \begin{bmatrix} 0 & 0 & 0 & 1 & -1 \\ \vdots & \vdots & \vdots & 0 & 0 \\ \vdots & \vdots & \vdots & \vdots & \vdots \\ 0 & 0 & 0 & 0 & 0 \end{bmatrix}_{L\times 5}, \quad S_c := \begin{bmatrix} 0 & -1 & 1 & 0 & 0 \\ 1 & -1 & 0 & 0 & 0 \end{bmatrix}, \quad \lambda_c(x,z) := \begin{bmatrix} \theta x_L \\ \eta z_1 z_2 \\ \mu \\ h^+(z_1, x_L) \\ h^-(x_1, x_L) \end{bmatrix},$$

so that the closed-loop stoichiometry matrix and propensity function can be written as $S_{cl} = \begin{bmatrix} S & S_m \\ 0 & S_c \end{bmatrix}$ and $\lambda_{cl}(x,z) = \begin{bmatrix} \lambda(x) \\ \lambda_c(x,z) \end{bmatrix}$, respectively. Note that $h^+$ and $h^-$ are functions that can take the forms given in Figure 3 depending on the adopted inhibition mechanism. By substituting the plant and controller components of $X_{cl}, S_{cl}$, and $\lambda_{cl}$ in (S48), we proceed as follows

$$\frac{d}{dt}\begin{bmatrix}\mathrm{Cov}\left[X\right] & \mathrm{Cov}\left[X,Z\right]\\ \mathrm{Cov}\left[Z,X\right] & \mathrm{Cov}\left[Z\right]\end{bmatrix} = \begin{bmatrix}S & S_m\\ 0 & S_c\end{bmatrix}\begin{bmatrix}\mathcal{D}\big(\mathbb{E}\left[\lambda(X)\right]\big) & 0\\ 0 & \mathcal{D}\big(\mathbb{E}\left[\lambda_c(X,Z)\right]\big)\end{bmatrix}\begin{bmatrix}S^T & 0\\ S_m^T & S_c^T\end{bmatrix}$$
$$+\begin{bmatrix}S & S_m\\ 0 & S_c\end{bmatrix}\begin{bmatrix}\mathrm{Cov}\left[\lambda(X),X\right] & \mathrm{Cov}\left[\lambda(X),Z\right]\\ \mathrm{Cov}\left[\lambda_c(X,Z),X\right] & \mathrm{Cov}\left[\lambda_c(X,Z),Z\right]\end{bmatrix}$$
$$+\begin{bmatrix}\mathrm{Cov}\left[X,\lambda(X)\right] & \mathrm{Cov}\left[X,\lambda_c(X,Z)\right]\\ \mathrm{Cov}\left[Z,\lambda(X)\right] & \mathrm{Cov}\left[Z,\lambda_c(X,Z)\right]\end{bmatrix}\begin{bmatrix}S^T & 0\\ S_m^T & S_c^T\end{bmatrix}$$

Thus we have

$$\begin{aligned}\frac{d}{dt}\mathrm{Cov}\left[X\right] &= S\mathcal{D}\big(\mathbb{E}\left[\lambda(X)\right]\big)S^T + S_m\mathcal{D}\big(\mathbb{E}\left[\lambda_c(X,Z)\right]\big)S_m^T + S\mathrm{Cov}\left[\lambda(X),X\right] + S_m\mathrm{Cov}\left[\lambda_c(X,Z),X\right]\\ &\quad + \mathrm{Cov}\left[X,\lambda(X)\right]S^T + \mathrm{Cov}\left[X,\lambda_c(X,Z)\right]S_m^T\\ \frac{d}{dt}\mathrm{Cov}\left[X,Z\right] &= S_m\mathcal{D}\big(\mathbb{E}\left[\lambda_c(X,Z)\right]\big)S_c^T + S\mathrm{Cov}\left[\lambda(X),Z\right] + S_m\mathrm{Cov}\left[\lambda_c(X,Z),Z\right] + \mathrm{Cov}\left[X,\lambda_c(X,Z)\right]S_c^T\\ \frac{d}{dt}\mathrm{Cov}\left[Z\right] &= S_c\mathcal{D}\big(\mathbb{E}\left[\lambda_c(X,Z)\right]\big)S_c^T + S_c\mathrm{Cov}\left[\lambda_c(X,Z),Z\right] + \mathrm{Cov}\left[Z,\lambda_c(X,Z)\right]S_c^T.\end{aligned}$$

Next, by substituting for $S_m, S_c$ and $\lambda_c(X,Z)$ and doing some algebraic calculations, we obtain

$$\begin{cases}\frac{d}{dt}\mathrm{Cov}\left[X\right] = S\mathcal{D}\big(\mathbb{E}\left[\lambda(X)\right]\big)S^T + \mathbb{E}\left[h^+\left(Z_1,X_L\right) - h^-\left(X_1,X_L\right)\right]e_1e_1^T + S\mathrm{Cov}\left[\lambda(X),X\right] + e_1\mathrm{Cov}\left[h\left(Z_1,X_1,X_L\right),X\right]\\ \qquad + \mathrm{Cov}\left[X,\lambda(X)\right]S^T + \mathrm{Cov}\left[X,h\left(Z_1,X_1,X_L\right)\right]e_1^T\\ \frac{d}{dt}\mathrm{Cov}\left[X,Z_1\right] = S\mathrm{Cov}\left[\lambda(X),Z_1\right] + \mathrm{Cov}\left[h(Z_1,X_1,X_L),Z_1\right]e_1 - \eta\mathrm{Cov}\left[X,Z_1Z_2\right]\\ \frac{d}{dt}\mathrm{Cov}\left[X,Z_2\right] = S\mathrm{Cov}\left[\lambda(X),Z_2\right] + \mathrm{Cov}\left[h(Z_1,X_1,X_L),Z_2\right]e_1 - \eta\mathrm{Cov}\left[X,Z_1Z_2\right] + \theta\mathrm{Cov}\left[X,X_L\right]\\ \frac{d}{dt}\mathrm{Var}\left[Z_1\right] = \mu + \eta\mathbb{E}\left[Z_1Z_2\right] - 2\eta\mathrm{Cov}\left[Z_1,Z_1Z_2\right]\\ \frac{d}{dt}\mathrm{Var}\left[Z_2\right] = \theta\mathbb{E}\left[X_L\right] + \eta\mathbb{E}\left[Z_1Z_2\right] - 2\eta\mathrm{Cov}\left[Z_2,Z_1Z_2\right] + 2\theta\mathrm{Cov}\left[X_L,Z_2\right]\\ \frac{d}{dt}\mathrm{Cov}\left[Z_1,Z_2\right] = \eta\mathbb{E}\left[Z_1Z_2\right] - \eta\mathrm{Cov}\left[Z_1+Z_2,Z_1Z_2\right] + \theta\mathrm{Cov}\left[X_L,Z_1\right],\end{cases} \tag{S49}$$

where, as in Figure 3, the total actuation propensity function is defined as $h(z_1,x_1,x_L) := h^+(z_1,x_L) - h^-(x_1,x_L)$, and $e_i$ is a vector whose entries are all zeros except the $i^{\text{th}}$ entry is one. Note that $h$ does not depend on $z_2$ in this section because only $a$PI controllers of Class 1 are considered. The set of differential equations in (S49) describe the evolution of the various covariances in the closed-loop network. Observe that it does not involve first and second order moments only (expectations and covariances), but also third order moments like $\mathrm{Cov}\left[X,Z_1Z_2\right]$ and $\mathrm{Cov}\left[Z_1+Z_2,Z_1Z_2\right]$ that have there own differential equations. This, in addition to the nonlinearity of $\lambda$ and $h$ (in general), give rise to the moment closure problem.

#### S7.2 Steady-State (Stationary) Analysis

Let $\mathbb{E}_\pi\left[\cdot\right]$, $\mathrm{Var}_\pi\left[\cdot\right]$, and $\mathrm{Cov}_\pi\left[\cdot,\cdot\right]$ denote the the stationary expectation, variance and covariance, respectively. Assuming that the closed-loop system is ergodic, the various time derivatives in (4) and (S49) are set to zero at stationarity. Particularly, we have the following relationships that hold regardless of what the plant is

$$\begin{aligned}\frac{d}{dt}\left(\mathbb{E}_\pi\left[Z_1\right] - \mathbb{E}_\pi\left[Z_2\right]\right) = 0 &\implies \mathbb{E}_\pi\left[X_L\right] = \frac{\mu}{\theta}\\ \frac{d}{dt}\mathbb{E}_\pi\left[Z_1\right] = 0 &\implies \mathbb{E}_\pi\left[Z_1Z_2\right] = \frac{\mu}{\eta}\\ \frac{d}{dt}\mathrm{Var}_\pi\left[Z_1\right] = 0 &\implies \mathrm{Cov}_\pi\left[Z_1,Z_1Z_2\right] = \frac{\mu}{\eta}\\ \frac{d}{dt}\left(\mathrm{Var}_\pi\left[Z_1\right] + \mathrm{Var}_\pi\left[Z_2\right] - 2\mathrm{Cov}_\pi\left[Z_1,Z_2\right]\right) = 0 &\implies \mathrm{Cov}_\pi\left[X_L,Z_1-Z_2\right] = \frac{\mu}{\theta}\\ \frac{d}{dt}\mathrm{Var}_\pi\left[Z_2\right] = 0 &\implies \mathrm{Cov}_\pi\left[Z_2,Z_1Z_2\right] = \frac{\mu + \theta\mathrm{Cov}_\pi\left[X_L,Z_2\right]}{\eta}.\end{aligned} \tag{S50}$$

These relationships will be useful in what follows, particularly in the asymptotic limit as $\eta \to \infty$.

#### S7.3 Application to Gene Expression

Consider the case where the plant is the gene expression network described in Figure 4(a). The plant stoichiometry matrix and propensity vector are given by

$$S = \begin{bmatrix} 0 & -1 & 0 \\ 1 & 0 & -1 \end{bmatrix} \qquad \text{and} \qquad \lambda(x) = \begin{bmatrix} k_1 x_1 \\ \gamma_1 x_1 \\ \gamma_2 x_2 \end{bmatrix}.$$

Then, by substituting $S$ and $\lambda$ in (4) and (S49), we obtain the the following set of differential equations for the expectations and covariances

$$\begin{cases} \frac{d}{dt}\mathbb{E}[X_1] = \mathbb{E}[h(Z_1, X_1, X_2)] - \gamma_1 \mathbb{E}[X_1] \\ \frac{d}{dt}\mathbb{E}[X_2] = k_1 \mathbb{E}[X_1] - \gamma_2 \mathbb{E}[X_2] \\ \frac{d}{dt}\mathbb{E}[Z_1] = \mu - \eta \mathbb{E}[Z_1 Z_2] \\ \frac{d}{dt}\mathbb{E}[Z_2] = \theta \mathbb{E}[X_2] - \eta \mathbb{E}[Z_1 Z_2], \end{cases}$$

$$\begin{cases} \frac{d}{dt}\mathrm{Var}[X_1] = \gamma_1 \mathbb{E}[X_1] + \mathbb{E}\left[h^+(Z_1, X_2) + h^-(X_1, X_2)\right] - 2\gamma_1 \mathrm{Var}[X_1] + 2\mathrm{Cov}[X_1, h(Z_1, X_1, X_2)] \\ \frac{d}{dt}\mathrm{Var}[X_2] = \gamma_2 \mathbb{E}[X_2] + k_1 \mathbb{E}[X_1] - 2\gamma_2 \mathrm{Var}[X_2] + 2k_1 \mathrm{Cov}[X_1, X_2] \\ \frac{d}{dt}\mathrm{Cov}[X_1, X_2] = k_1 \mathrm{Var}[X_1] - (\gamma_1 + \gamma_2)\mathrm{Cov}[X_1, X_2] + \mathrm{Cov}[X_2, h(Z_1, X_1, X_2)] \\ \frac{d}{dt}\mathrm{Cov}[X_1, Z_1] = -\gamma_1 \mathrm{Cov}[X_1, Z_1] + \mathrm{Cov}[h(Z_1, X_1, X_2), Z_1] - \eta \mathrm{Cov}[X_1, Z_1 Z_2] \\ \frac{d}{dt}\mathrm{Cov}[X_1, Z_2] = -\gamma_1 \mathrm{Cov}[X_1, Z_2] + \mathrm{Cov}[h(Z_1, X_1, X_2), Z_2] - \eta \mathrm{Cov}[X_1, Z_1 Z_2] + \theta \mathrm{Cov}[X_1, X_2] \\ \frac{d}{dt}\mathrm{Cov}[X_2, Z_1] = -\gamma_2 \mathrm{Cov}[X_2, Z_1] + k_1 \mathrm{Cov}[X_1, Z_1] - \eta \mathrm{Cov}[X_2, Z_1 Z_2] \\ \frac{d}{dt}\mathrm{Cov}[X_2, Z_2] = -\gamma_2 \mathrm{Cov}[X_2, Z_2] + k_1 \mathrm{Cov}[X_1, Z_2] - \eta \mathrm{Cov}[X_2, Z_1 Z_2] + \theta \mathrm{Var}[X_2] \\ \frac{d}{dt}\mathrm{Var}[Z_1] = \mu + \eta \mathbb{E}[Z_1 Z_2] - 2\eta \mathrm{Cov}[Z_1 Z_2, Z_1] \\ \frac{d}{dt}\mathrm{Var}[Z_2] = \theta \mathbb{E}[X_2] + \eta \mathbb{E}[Z_1 Z_2] - 2\eta \mathrm{Cov}[Z_1 Z_2, Z_2] + 2\theta \mathrm{Cov}[X_2, Z_2] \\ \frac{d}{dt}\mathrm{Cov}[Z_1, Z_2] = \eta \mathbb{E}[Z_1 Z_2] - \eta \mathrm{Cov}[Z_1 Z_2, Z_1 + Z_2] + \theta \mathrm{Cov}[X_2, Z_1] \end{cases}$$

**Steady-State (Stationary) Analysis:** Assuming that the closed-loop system is ergodic, the time derivatives at stationarity are set to zero. We have

$$\mathbb{E}_\pi[X_2] = \frac{\mu}{\theta}; \qquad \mathbb{E}_\pi[X_1] = \frac{\gamma_2}{k_1}\frac{\mu}{\theta}; \qquad \mathbb{E}_\pi[h(Z_1, X_1, X_2)] = \frac{\gamma_1\gamma_2}{k_1}\frac{\mu}{\theta}; \qquad \mathbb{E}_\pi[Z_1 Z_2] = \frac{\mu}{\eta};$$
$$\mathbb{E}_\pi\left[h^+(Z_1, X_2) + h^-(X_1, X_2)\right] = 2\mathbb{E}_\pi\left[h^+(Z_1, X_2)\right] - \frac{\gamma_1\gamma_2}{k_1}\frac{\mu}{\theta}.$$

To compute the steady-state variance $\mathrm{Var}_\pi[X_2]$, we use the following set of algebraic equations

$$\begin{aligned} \frac{d}{dt}\mathrm{Var}_\pi[X_2] = 0 &\implies \mathrm{Var}_\pi[X_2] = \frac{\mu}{\theta} + \frac{k_1}{\gamma_2}\mathrm{Cov}_\pi[X_1, X_2] \\ \frac{d}{dt}\mathrm{Cov}_\pi[X_1, X_2] = 0 &\implies \mathrm{Cov}_\pi[X_1, X_2] = \frac{k_1}{\gamma_1 + \gamma_2}\mathrm{Var}_\pi[X_1] + \frac{1}{\gamma_1 + \gamma_2}\mathrm{Cov}_\pi[X_2, h(Z_1, X_1, X_2)] \\ \frac{d}{dt}\mathrm{Var}_\pi[X_1] = 0 &\implies \mathrm{Var}_\pi[X_1] = \frac{\mathbb{E}_\pi[h^+(Z_1, X_2)] + \mathrm{Cov}_\pi[X_1, h(Z_1, X_1, X_2)]}{\gamma_1} \\ \frac{d}{dt}\mathrm{Cov}_\pi[X_2, Z_1 - Z_2] = 0 &\implies \mathrm{Cov}_\pi[X_1, Z_1 - Z_2] = \frac{\gamma_2}{k_1}\frac{\mu}{\theta} + \frac{\theta}{k_1}\mathrm{Var}_\pi[X_2], \end{aligned} \tag{S51}$$

where the last equality follows by exploiting the fact that $\text{Cov}_\pi[X_2, Z_1 - Z_2] = \mu/\theta$ from (S50). Observe that these algebraic equations cannot be solved exactly for $\text{Var}_\pi[X_2]$ because of the moment closure problem. However, to proceed, we give an approximation for the covariance terms $\text{Cov}_\pi[X_i, h(Z_1, X_1, X_2)]$ for $i = 1, 2$. The approximation essentially (1) exploits a second order Taylor expansion of the function $h$ around the stationary expected values , and (2) exploits the fact that for large $\eta$, $\mathbb{E}_\pi[Z_1 Z_2] = \frac{\mu}{\eta} \approx 0$ and $Z_2$ is assumed to be close to zero unlike $Z_1$ which takes positive values actuating the plant. In fact, these approximations are summarized below.

$$\mathbb{E}_\pi[Z_1 Z_2] \approx 0, \qquad \text{Cov}_\pi[X_1, Z_2] \approx 0, \qquad \text{Cov}_\pi[X_2, Z_2] \approx 0$$

$$h(Z_1, X_1, X_2) \approx \bar{h} + \begin{bmatrix} \partial_{z_1}\bar{h} & \partial_{x_1}\bar{h} & \partial_{x_2}\bar{h} \end{bmatrix} \begin{bmatrix} Z_1 - \mathbb{E}_\pi[Z_1] \\ X_1 - \mathbb{E}_\pi[X_1] \\ X_2 - \mathbb{E}_\pi[X_2] \end{bmatrix} + \frac{1}{2} \begin{bmatrix} Z_1 - \mathbb{E}_\pi[Z_1] \\ X_1 - \mathbb{E}_\pi[X_1] \\ X_2 - \mathbb{E}_\pi[X_2] \end{bmatrix}^T \begin{bmatrix} \partial_{z_1}^2\bar{h} & \partial_{z_1}\partial_{x_1}\bar{h} & \partial_{z_1}\partial_{x_2}\bar{h} \\ \partial_{x_1}\partial_{z_1}\bar{h} & \partial_{x_1}^2\bar{h} & \partial_{x_1}\partial_{x_2}\bar{h} \\ \partial_{x_2}\partial_{z_1}\bar{h} & \partial_{x_2}\partial_{x_1}\bar{h} & \partial_{x_2}^2\bar{h} \end{bmatrix} \begin{bmatrix} Z_1 - \mathbb{E}_\pi[Z_1] \\ X_1 - \mathbb{E}_\pi[X_1] \\ X_2 - \mathbb{E}_\pi[X_2] \end{bmatrix},$$

where $\bar{h} := h\Big(\mathbb{E}_\pi[Z_1], \mathbb{E}_\pi[X_1], \mathbb{E}_\pi[X_2]\Big)$. Using Appendix S11 (with $X := \begin{bmatrix} Z_1 & X_1 & X_2 \end{bmatrix}^T$, $F(X) = X_1$, and $G(X) = h(Z_1, X_1, X_2)$), we can approximate $\text{Cov}_\pi[X_1, h(Z_1, X_1, X_2)]$ up to first order (or second order if the stationary distribution is close to a normal distribution) as

$$\begin{aligned} \text{Cov}_\pi[X_1, h(Z_1, X_1, X_2)] &\approx \begin{bmatrix} 0 & 1 & 0 \end{bmatrix} \begin{bmatrix} \text{Var}_\pi[Z_1] & \text{Cov}_\pi[Z_1, X_1] & \text{Cov}_\pi[Z_1, X_2] \\ \text{Cov}_\pi[X_1, Z_1] & \text{Var}_\pi[X_1] & \text{Cov}_\pi[X_1, X_2] \\ \text{Cov}_\pi[X_2, Z_1] & \text{Cov}_\pi[X_2, X_1] & \text{Var}_\pi[X_2] \end{bmatrix} \begin{bmatrix} \partial_{z_1}\bar{h} \\ \partial_{x_1}\bar{h} \\ \partial_{x_2}\bar{h} \end{bmatrix} \\ &\approx \sigma_1 \text{Cov}_\pi[X_1, Z_1] - \sigma_3 \text{Var}_\pi[X_1] - \sigma_4 \text{Cov}_\pi[X_1, X_2], \end{aligned}$$

where

$$\begin{aligned} \sigma_1 &:= \partial_{z_1} h(\mathbb{E}_\pi[Z_1], \mathbb{E}_\pi[X_1], \mathbb{E}_\pi[X_2]) > 0 \\ \sigma_3 &:= -\partial_{x_1} h(\mathbb{E}_\pi[Z_1], \mathbb{E}_\pi[X_1], \mathbb{E}_\pi[X_2]) \geq 0 \\ \sigma_4 &:= -\partial_{x_2} h(\mathbb{E}_\pi[Z_1], \mathbb{E}_\pi[X_1], \mathbb{E}_\pi[X_2]) \geq 0. \end{aligned}$$

Similarly, we can approximate $\text{Cov}_\pi[X_2, h(Z_1, X_1, X_2)]$ as

$$\begin{aligned} \text{Cov}_\pi[X_2, h(Z_1, X_1, X_2)] &\approx \begin{bmatrix} 0 & 0 & 1 \end{bmatrix} \begin{bmatrix} \text{Var}_\pi[Z_1] & \text{Cov}_\pi[Z_1, X_1] & \text{Cov}_\pi[Z_1, X_2] \\ \text{Cov}_\pi[X_1, Z_1] & \text{Var}_\pi[X_1] & \text{Cov}_\pi[X_1, X_2] \\ \text{Cov}_\pi[X_2, Z_1] & \text{Cov}_\pi[X_2, X_1] & \text{Var}_\pi[X_2] \end{bmatrix} \begin{bmatrix} \partial_{z_1}\bar{h} \\ \partial_{x_1}\bar{h} \\ \partial_{x_2}\bar{h} \end{bmatrix} \\ &\approx \sigma_1 \text{Cov}_\pi[X_2, Z_1] - \sigma_3 \text{Cov}_\pi[X_2, X_1] - \sigma_4 \text{Var}_\pi[X_2]. \end{aligned}$$

Invoking the approximations $\text{Cov}_\pi[X_1, Z_2] \approx \text{Cov}_\pi[X_2, Z_2] \approx 0$ and the last equation in (S51), we obtain

$$\begin{aligned} \text{Cov}_\pi[X_1, Z_1] &\approx \text{Cov}_\pi[X_1, Z_1 - Z_2] = \frac{\gamma_2}{k_1}\frac{\mu}{\theta} + \frac{\theta}{k_1}\text{Var}_\pi[X_2] \\ \text{Cov}_\pi[X_2, Z_1] &\approx \text{Cov}_\pi[X_2, Z_1 - Z_2] = \frac{\mu}{\theta}. \end{aligned}$$

Then we have

$$\begin{aligned} \text{Cov}_\pi[X_1, h(Z_1, X_1, X_2)] &\approx \sigma_1 \frac{\gamma_2}{k_1}\frac{\mu}{\theta} + \sigma_1 \frac{\theta}{k_1}\text{Var}_\pi[X_2] - \sigma_3 \text{Var}_\pi[X_1] - \sigma_4 \text{Cov}_\pi[X_1, X_2] \\ \text{Cov}_\pi[X_2, h(Z_1, X_1, X_2)] &\approx \sigma_1 \frac{\mu}{\theta} - \sigma_3 \text{Cov}_\pi[X_1, X_2] - \sigma_4 \text{Var}_\pi[X_2]. \end{aligned}$$

Finally, by substituting for $\text{Cov}_\pi[X_1, h(Z_1, X_1, X_2)]$ and $\text{Cov}_\pi[X_2, h(Z_1, X_1, X_2)]$ in (S51), we obtain the following set of algebraic (approximate) equations

$$\begin{aligned} \text{Var}_\pi[X_2] &\approx \frac{\mu}{\theta} + \frac{k_1}{\gamma_2}\text{Cov}_\pi[X_1, X_2] \\ \text{Cov}_\pi[X_1, X_2] &\approx \frac{k_1}{\gamma_1 + \gamma_2}\text{Var}_\pi[X_1] + \frac{1}{\gamma_1 + \gamma_2}\left(\sigma_1\frac{\mu}{\theta} - \sigma_4 \text{Var}_\pi[X_2] - \sigma_3 \text{Cov}[X_1, X_2]\right) \\ \text{Var}_\pi[X_1] &\approx \frac{\gamma_2}{k_1}\frac{\mu}{\theta} + \frac{1}{\gamma_1}\left(h^-\left(\frac{\gamma_2}{k_1}r, r\right) + \partial_{x_1}\partial_{x_2} h\left(\frac{\gamma_2}{k_1}r, r\right)\text{Cov}_\pi[X_1, X_2] + \frac{1}{2}\partial_{x_1}^2 h\left(\frac{\gamma_2}{k_1}r, r\right)\text{Var}_\pi[X_1] \right. \\ &\left. + \frac{1}{2}\partial_{x_2}^2 h\left(\frac{\gamma_2}{k_1}r, r\right)\text{Var}_\pi[X_2] + \sigma_1\frac{\gamma_2}{k_1}\frac{\mu}{\theta} + \sigma_1\frac{\theta}{k_1}\text{Var}_\pi[X_2] - \sigma_3 \text{Var}_\pi[X_1] - \sigma_4 \text{Cov}_\pi[X_1, X_2]\right), \end{aligned}$$

where $r := \mu/\theta$. This can be written in matrix form as

$$\begin{bmatrix} 1+\frac{\sigma_3}{\gamma_1}-\frac{1}{2\gamma_1}\partial_{x_1}^2 h^-\left(\frac{\gamma_2}{k_1}r,r\right) & -\frac{\sigma_1}{\gamma_1}\frac{\theta}{k_1}-\frac{1}{2\gamma_1}\partial_{x_2}^2 h^-\left(\frac{\gamma_2}{k_1}r,r\right) & \frac{\sigma_4}{\gamma_1}-\frac{1}{\gamma_1}\partial_{x_1}\partial_{x_2}h^-\left(\frac{\gamma_2}{k_1}r,r\right) \\ 0 & 1 & -\frac{k_1}{\gamma_2} \\ -k_1 & \sigma_4 & \gamma_1+\gamma_2+\sigma_3 \end{bmatrix} \begin{bmatrix} \mathrm{Var}_\pi\left[X_1\right] \\ \mathrm{Var}_\pi\left[X_2\right] \\ \mathrm{Cov}_\pi\left[X_1,X_2\right] \end{bmatrix}$$
$$= \begin{bmatrix} \left(1+\frac{\sigma_1}{\gamma_1}\right)\frac{\gamma_2}{k_1}r+\frac{1}{\gamma_1}h^-\left(\frac{\gamma_2}{k_1}r,r\right) \\ \frac{\mu}{\theta} \\ \sigma_1\frac{\mu}{\theta} \end{bmatrix}.$$

Finally, solving for $\mathrm{Var}_\pi\left[X_2\right]$, we arrive at

$$\mathrm{Var}_\pi\left[X_2\right] \approx r\left[\frac{(\gamma_1+\gamma_2+\sigma_3)(\gamma_1\gamma_2+\gamma_2\sigma_3+\sigma_1 k_1)+k_1\left(\gamma_2(\gamma_1+\sigma_4)+\frac{k_1}{r}h^-\left(\frac{\gamma_2}{k_1}r,r\right)\right)}{(\gamma_1+\gamma_2+\sigma_3)(\gamma_1\gamma_2+\gamma_2\sigma_3+k_1\sigma_4)-\sigma_1 k_1\theta}\right].$$

This is a general formula that encompasses the standalone $a$I controller and the three $a$PI controllers of Class 1 (with different inhibition mechanisms), that are addressed as special cases next.

***a*I:** For this controller, the propensities of the actuation reactions are given by

$$h^+(z_1,x_2)=kz_1 \qquad \text{and} \qquad h^-(x_1,x_2)=0,$$

which implies that $h(z_1,x_1,x_2)=kz_1$. Then $\sigma_1=k$ and $\sigma_3=\sigma_4=0$.

***a*PI of Class 1 with Additive Inhibition:** The propensities of the actuation reactions are given by

$$h^+(z_1,x_2)=kz_1+\frac{\alpha}{1+(x_2/\kappa)^n} \qquad \text{and} \qquad h^-(x_1,x_2)=0,$$

which implies that $h(z_1,x_1,x_2)=kz_1+\frac{\alpha}{1+(x_2/\kappa)^n}$. Then we have

$$\sigma_1=k, \qquad \sigma_3=0, \qquad \text{and} \quad \sigma_4=\frac{\alpha}{r}\frac{n(r/\kappa)^n}{\left[1+(r/\kappa)^n\right]^2}.$$

***a*PI of Class 1 with Multiplicative Inhibition:** The propensities of the actuation reactions are given by

$$h^+(z_1,x_2)=\frac{kz_1}{1+(x_2/\kappa)^n} \qquad \text{and} \qquad h^-(x_1,x_2)=0,$$

which implies that $h(z_1,x_1,x_2)=\frac{kz_1}{1+(x_2/\kappa)^n}$. Then we have

$$\sigma_1=\frac{k}{1+(r/\kappa)^n}, \qquad \sigma_3=0, \qquad \text{and} \qquad \sigma_4=\frac{k\mathbb{E}_\pi\left[Z_1\right]}{r}\frac{n(r/\kappa)^n}{\left[1+(r/\kappa)^n)\right]^2}.$$

We are left with approximating $\mathbb{E}_\pi\left[Z_1\right]$. This can be done by recalling that $\mathbb{E}_\pi\left[h(Z_1,X_1,X_2)\right]=\frac{\gamma_1\gamma_2}{k_1}\frac{\mu}{\theta}$ and using a first order Taylor series expansion of $h$ around stationarity. That is

$$\begin{aligned} \mathbb{E}_\pi\left[h\Big(\mathbb{E}_\pi\left[Z_1\right],\mathbb{E}_\pi\left[X_1\right],\mathbb{E}_\pi\left[X_2\right]\Big)\right] &\approx \frac{\gamma_1\gamma_2}{k_1}r \\ \frac{k\mathbb{E}_\pi\left[Z_1\right]}{1+(r/\kappa)^n} &\approx \frac{\gamma_1\gamma_2}{k_1}r \\ \mathbb{E}_\pi\left[Z_1\right] &\approx \frac{\gamma_1\gamma_2}{kk_1}r\left[1+(r/\kappa)^n\right]. \end{aligned}$$

Finally, substituting for $\mathbb{E}_\pi\left[Z_1\right]$ in $\sigma_4$ yields

$$\sigma_4=\frac{\gamma_1\gamma_2}{k_1}\frac{n(r/\kappa)^n}{1+(r/\kappa)^n}.$$

***a*PI of Class 1 with Degradation Inhibition:** The propensities of the actuation reactions are given by

$$h^+(z_1, x_2) = kz_1 \qquad \text{and} \qquad h^-(x_1, x_2) = \delta x_2 \frac{x_1}{x_1 + \kappa_1},$$

which implies that $h(z_1, x_1, x_2) = kz_1 - \delta x_2 \frac{x_1}{x_1+\kappa_1}$. Then we have

$$\sigma_1 = k, \qquad \sigma_3 = \delta r \frac{\kappa_1}{(\gamma_2 r/k_1 + \kappa_1)^2}, \qquad \text{and} \quad \sigma_4 = r \frac{\delta \gamma_2 / k_1}{\gamma_2 r/k_1 + \kappa_1}.$$

The results are summarized in Table 1.

### S8 A Genetic Design of the Second Order *a*PID Controller

Here we propose and describe a particular genetic design in E.Coli that realizes the second order *a*PID controller topology presented in Figure 5. The design is very similar to that of the third order *a*PID controller presented in Figure 10. We, once again, perform numerical simulations using biologically realistic parameters to demonstrate the effectiveness of the controllers in ameliorating the dynamic performance. The genetic circuit is depicted in Figure S10(a) where the controller circuit augments the I-control module (in blue), which is based on [2], with additional circuitry to implement additional P and D controls (in red and green). The P control module is implemented via the Mflon protease which is capable of degrading the input species $\mathbf{X_1}$; whereas the D control module is implemented via the expression of the $\sigma$ factor *sigW* by using a mutated weaker version of the AraC-responsive promoter $P_{BAD}$, which is denoted here by $P^*_{BAD}$ [3]. The additional disturbance circuit (in yellow) serves as a source of external perturbation to the closed-loop circuit by degrading the regulated output $\mathbf{X_2}$. The set of ODEs describing the deterministic dynamics are also shown Figure S10(a) and the various parameters are chosen to reflect biologically realistic regimes and account for controller species dilution $\delta_c$.

In particular, the numerical values for the rates $\gamma_1 = \gamma_2 = \delta_c = 0.028$ min$^{-1}$ and $\eta = 0.05$ nM$^{-1}$ min$^{-1}$ are taken from [2] to reflect a doubling time of 25 min in bacteria and a relatively strong sequestration rate, respectively. Futhermore, the rates $k_1 = 1$ min$^{-1}$, $k = 0.05$ min$^{-1}$, $\theta = 1$ min$^{-1}$ and $\mu = 50$ nM min$^{-1}$ are also chosen to respect the ranges given in [2]. Note that the choices of $\mu$ and $\theta$ yield a set-point of 50 nM. The disturbance is chosen here to be three times the dilution rate, that is $\Delta = 3 \times 0.028$ min$^{-1}$. As for $\kappa_1$, we select it to reflect a half activation threshold equal to one fifth of the set-point, that is $\kappa_1 = 10$ nM. For I-control, we set $\delta = \beta = 0$. For PI-control, we set $\beta = 0$ and $\delta = 5 \times 0.028$ min$^{-1}$ which reflects a reasonable degradation rate which is five times faster than dilution. For PID-control we set $\delta = 5 \times 0.028$ min$^{-1}$ and $\beta = \theta/2 = 0.5$ min$^{-1}$ since both $\beta$ and $\theta$ are gene expression rates and are thus comparable. Recall that for the second order *a*PID, the set-point is given by $\mu/(\theta - \beta)$ with $\beta < \theta$. Hence to do a fair comparison with I- and PI-control, we set $\mu = 25$ nM min$^{-1}$ to keep the set-point at 50 nM. Figure S10(b) shows the simulation results for I, PI and PID control. The responses are shown for a step change of setpoint (by doubling $\mu$), which is tunable with HSL [2], at $t = 8$ hrs and for a step change of disturbance ($\Delta = 0$ to $\Delta = 5 \times 0.028$ min$^{-1}$), which is tunable with aTc, at $t = 16$ hrs. The simulations demonstrate that the full PID controller is capable of dramatically enhancing the stability and performance by not only shaping the transient dynamics but also reducing the steady-state error that can be incurred by the dilution effect (see [4], [2], [5]).

### S9 Numerical Values

In this section, we provide the numerical values of the various parameters adopted in each figure.

**Figure 4(d):**

- **Fixed Parameters:**
  For all inhibition mechanisms: $\mu = 10,\ \theta = 2,\ \eta = 100,\ k = 3,\ n = 1,\ k_1 = \gamma_1 = \gamma_2 = 2$ and $r := \mu/\theta = 5$.
  For Additive Inhibition: $\kappa = 1$
  For Degradation Inhibition: $\kappa_1 = 0.01$.

- **Variable Parameters:** $\alpha \in [0, \alpha_{\mathrm{TH}}],\ \kappa^{-1} \in [0, 0.7]$ and $\delta \in [0, 6]$ with $\alpha_{\mathrm{TH}} := \frac{\gamma_1 \gamma_2}{k_1} r \left(1 + (r/\kappa)^n\right) = 60$.

- **Remark:** Throughout the paper, when the proportional controller is realized with degradation inhibition, i.e. $h^-(x_1, x_L) = \delta x_L^n \frac{x_1}{x_1+\kappa_1}$, $\kappa_1$ is chosen to be small. This is not a requirement for the controllers; in fact, $\kappa_1$ can

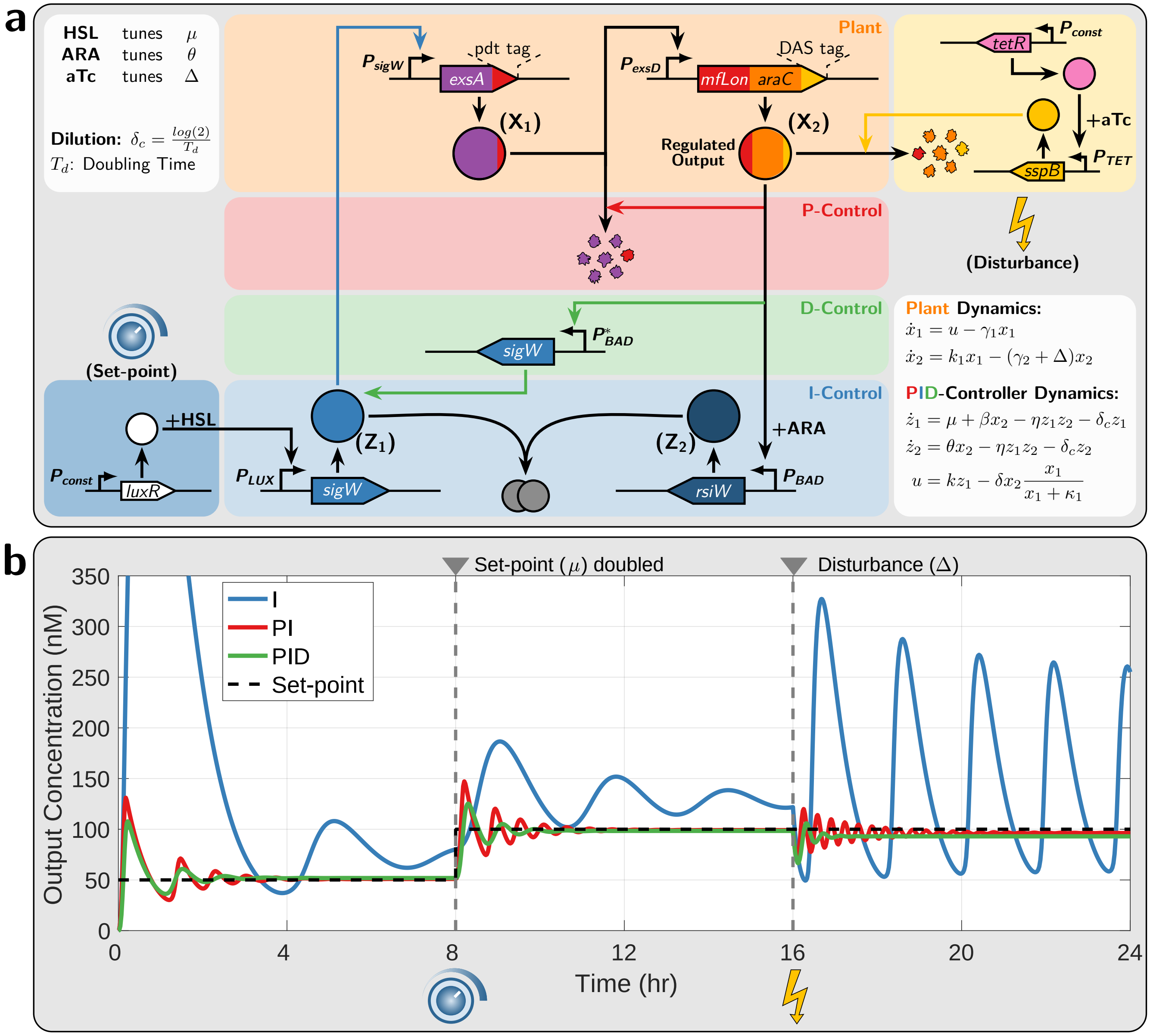

Figure S10: **A Genetic Implementation of the Second Order *a*PID Controller. (a) Circuit Design.** The genetic closed-loop circuit is comprised of the plant (in orange) involving two species $\mathbf{X_1}$ and $\mathbf{X_2}$, the *a*PID controller involving $\mathbf{Z_1}$ and $\mathbf{Z_2}$ and the disturbance network (in yellow). The objective is to force the regulated output $\mathbf{X_2}$ to track a tunable set-point, despite the injected disturbance, and enhance the transient dynamic response. The antithetic integral control is implemented via the sequestration between the $\sigma$ factor (SigW denoted by $\mathbf{Z_1}$) and anti-$\sigma$ factor (RsiW denoted by $\mathbf{Z_2}$) that are driven by the promoters $P_{LUX}$ and $P_{BAD}$ [2], respectively. The set-point is encrypted in the expression rate $\mu$ of $\mathbf{Z_1}$ which is tunable with homoserine lactone (HSL). The plant is comprised of two genes. The first gene encodes *exsA* [6] fused to the degradation pdt tag (recognized by Mflon), and is driven by a SigW-responsive promoter $P_{sigW}$. The second gene is driven by the ExsA-responsive promoter $P_{exsD}$ [6] and encodes the protease MfLon and the transcription factor *araC* fused to the ssrA(DAS) degradation tag. The disturbance circuit (in yellow) increases the degradation rate of $\mathbf{X_2}$ by expressing *sspB* at a tunable rate, with anhydrotetracycline (aTc), which in turn recognizes the DAS tag in $\mathbf{X_2}$ and sends it to the endogenous degradation machinery [7]. The MfLon in $\mathbf{X_2}$ is capable of degrading $\mathbf{X_1}$, while the AraC is capable of activating both $P_{BAD}$ and its weaker version $P^*_{BAD}$ [3] (that is $\beta < \theta$). The promoter $P_{BAD}$ drives the integral control module, while $P^*_{BAD}$ drives another copy of a gene expressing *sigW*. **(b) Deterministic Simulations.** Simulation of the closed-loop dynamics with I, PI and PID control. The plot shows the dynamic response of the regulated output $\mathbf{X_2}$ to a step change in the set-point at $t = 8$ hrs and to a disturbance injection at $t = 16$ hrs. The simulations are carried out using biologically realistic numerical values for the various parameters.

be used as an additional tuning parameter that can be shown to have benefits on shaping the dynamic response. To see this, one can look at the linearization of $h^-$ around the fixed point given by $K_P \tilde{x}_L + K'_P \tilde{x}_1$ with

$$K_P := \partial_{x_L} h^-(\bar{x}_1, \bar{x}_L) = \delta n \bar{x}_L^{n-1} \frac{\bar{x}_1}{\bar{x}_1 + \kappa_1} \quad \text{and} \quad K'_P := \partial_{x_1} h^-(\bar{x}_1, \bar{x}_L) = \delta \bar{x}_L^n \frac{\kappa_1}{(\bar{x}_1 + \kappa_1)^2}.$$

Hence the controller has two proportional feedback actions from $x_1$ and $x_L$ (state feedback). In general, the additional proportional action with gain $K'_P$ can be leveraged and exploited as yet an additional tuning knob to enhance the performance. However, our focus in this paper is to track the effects of the output feedback alone. Hence, by setting $\kappa_1$ to be small, $K'_P$ becomes negligible, and as a result, only output feedback effectively enters the dynamics.

**Figure 4(e):**

- **Fixed Parameters:** Same as the fixed parameters in Figure 4(d), except $\gamma_2 = 7$ to make the dynamics less oscillatory and focus on behavior of the stationary variance.
- **Variable Parameters:** $\alpha \in [0, \alpha_{\text{TH}}]$, $\kappa^{-1} \in [0, 5]$ and $\delta \in [0, 200]$ with $\alpha_{\text{TH}} := \frac{\gamma_1\gamma_2}{k_1} r\left(1 + (r/\kappa)^n\right) = 210$.
- **Remark:** The temporal evolution of the variance is calculated by running $N = 10^4$ stochastic trajectories, while the stationary variance is calculated by running a single simulation long enough ($t_{\text{Final}} = 2 \times 10^4$) and then taking time averages.

**Figure 6(b):**

- **Fixed Parameters:** $\mu = 10\ (t < 0)$, $\mu = 20\ (t > 0)$, $\theta = 2$, $\eta = 100$, $k = 4$, $\kappa_1 = 10^{-4}$, $n = 1$, $k_1 = \gamma_1 = \gamma_2 = 0.1$.
- **Variable Parameters:** $\delta \in [0, 1500]$.

**Figure 6(c):**

- **Fixed Parameters:** $\mu = 10\ (t < 0)$, $\mu = 20\ (t > 0)$, $\kappa_1 = 10^{-4}$, $n = 1$, $k_1 = \gamma_1 = \gamma_2 = 0.1$.
- **Variable Parameters:**
  For the second order $a$PID: $a \in \left[\frac{\gamma_1+\gamma_2}{2}, (2+\sqrt{2})\frac{\gamma_1+\gamma_2}{2}\right] = [0.1, 0.34]$. $(K_P, K_I, K_D, \omega)$ are computed for each $a$ using (S46), and $(\delta, \eta, k, \beta, \theta)$ are then computed using (S26). Only $(K_P, K_I, K_D)$ and $(\delta, \beta, k)$ are shown in the figure.
  For the third order $a$PID: $\theta = 2$, $\eta = 100$, $\kappa_0 = 300$, $\alpha_0 = 10^4$, $n_0 = 1$ and $a \in [0.05, 10]$. $(K_P, K_I, K_D, \omega)$ are computed for each $a$ using (S46), and $(\delta, \delta_0, k, \gamma_0)$ are then computed using (S31). Only $(K_P, K_I, K_D)$ and $(\delta, \delta_0, k)$ are shown in the figure.
  For the fourth order $a$PID: $\theta = 2$, $\eta = \eta_0 = 100$, $\theta_0 = 1, \bar{z}_3 = 10$ and $a \in [0.05, 10]$. $(K_P, K_I, K_D, \omega)$ are computed for each $a$ using (S46), and $(\delta, \alpha_1, \alpha_2, k, \mu_0)$ are then computed using (S36) and $\mu_0 = \theta_0(\alpha_1\bar{z}_3 + \alpha_2 r)$ with $r = \mu/\theta = 10$. Only $(K_P, K_I, K_D)$ and $(\delta, \alpha_2, k)$ are shown in the figure.

**Figure 7:**

- **Fixed Parameters:** $\mu = 3\ (t < 0)$, $\mu = 6\ (t > 0)$, $\kappa_1 = 10^{-4}$, $n = 1$, $k = 0.015$, $k_i = \gamma_i = 0.1$, $\kappa_F = 0.01$, $\gamma_F = 0.3$.
- **Variable Parameters:**
  For the second order $a$PID: $\eta = 10^{-4}, \delta \in [0, 0.3]$, $\beta \in [0, 10]$, $\theta = \beta + \mu/10$.
  For the third order $a$PID: $\theta = 3$, $\eta = 10^4$, $\kappa_0 = 10$, $\alpha_0 = 1$, $n_0 = 1$, $\gamma_0 = 0.2$ and $\delta \in [0, 0.1]$, $\delta_0 \in [0, 4]$.
  For the fourth order $a$PID: $\theta = 3$, $\eta = \eta_0 = 10^4$, $\mu_0 = 3$, $\theta_0 = 1$, $\alpha_1 = 0.2$ and $\alpha_2 \in [0, 0.1]$, $k_0 \in [0, 0.3]$.
- **Remark 1:** $\gamma_F$ is a parameter that specifies the feedback strength within the plant. It is chosen here to be larger than $\gamma_i$'s to give rise to an unstable open loop in order to challenge the controllers.
- **Remark 2:** The performance index weights are chosen to be $w_1 = w_2 = 1$ and $w_3 = 300$ to compensate for the different units (time versus concentration) and penalize the overshoot more than the settling and rise times.
- **Remark 3:** For the second order $a$PID, $\theta$ is set to be $\beta + \mu/10$ so that the set point for $t < 0$ is $\mu/(\theta - \beta) = 10$, and $\eta$ is chosen to be small because otherwise only small values of $K_D$ will be achievable (see (S4)).
- **Remark 4:** The various contours are computed using the mappings in Section S4 by finding the relationships between the biomolecular parameters of the horizontal and vertical axes of the intensity plots for a given value of the particular PID gain.

**Figure 8:** The plant parameters and initial conditions are taken from https://www.ebi.ac.uk/biomodels/. The fixed controller parameters are $\mu = 100,\ \eta = 0.1$ and $k = 100$. For the $a$I controller, we set $\beta = \delta = 0$ and $\theta = 1$. For the $a$PI, we set $\beta = 0,\ \delta = 0.01,\ \theta = 1$. For the $a$PID, we set $\beta = 50,\ \delta = 0.1,\ \theta = 51$.

**Figure 9(a):**

- **Fixed Parameters:** $\mu = 10,\ \kappa_1 = 10^{-2},\ n = 1,\ k_1 = \gamma_1 = 2,\ \gamma_2 = 7$ and $K_P = 0,\ K_I = 3,\ \omega_c = 10$.
- **Variable Parameters:**
For the second order $a$PID: $K_D \in [0, K_D^{\max}]$ with $K_D^{\max} = 1.34$. $(\delta, \eta, k, \beta, \theta)$ are computed for each $K_D$ using (S26). Only $(\delta, \beta, k)$ are shown in the figure.
For the third order $a$PID: $\theta = 2,\ \eta = 100,\ \kappa_0 = 1,\ \alpha_0 = 1,\ n_0 = 1,\ k = 3,\ \gamma_0 = 10$ and $K_D \in [0, 50]$. $(\delta, \delta_0)$ are computed for each $K_D$ using (S31).
For the fourth order $a$PID: $\theta = 2,\ \eta = \eta_0 = 100,\ \theta_0 = 1,\ \bar{z}_3 = 1,\ k = 3,\ \alpha_1 = 10$ with $K_D \in [0, 50]$. $(\alpha_2, \mu_0)$ are then computed using (S36) and $\mu_0 = \theta_0(\alpha_1 \bar{z}_3 + \alpha_2 r)$ with $r = \mu/\theta = 5$. Only $(\alpha_1, \alpha_2)$ are shown in the figure.
- **Remark 1:** For the second order $a$PID, with $K_P = 0$ and $(K_I, \omega_c)$ fixed, $K_D$ cannot be designed to be higher than $K_D^{\max} = \frac{\bar{u}}{r}\left(\frac{\bar{u}}{4K_I\mu} - \frac{1}{\omega_c}\right)$ which is derived from (S27).
- **Remark 2:** The temporal evolution of the variance and the stationary distribution are calculated by running $N = 32000$ stochastic trajectories, while the stationary variance is calculated by running a single simulation long enough ($t_{\text{Final}} = 2 \times 10^4$) and then taking time averages.

**Figure 9(b):**

- **Fixed Parameters:** $\mu = 3,\ \kappa_1 = 10^{-4},\ n = 1,\ k_i = \gamma_i = \gamma_F = 0.1, \kappa_F = 0.01$ and $K_P = 0,\ K_I = 0.03,\ \omega_c = 0.5$.
- **Variable Parameters:**
For the second order $a$PID: $K_D \in [0, K_D^{\max}]$ with $K_D^{\max} = 0.71$. $(\delta, \eta, k, \beta, \theta)$ are computed for each $K_D$ using (S26). Only $(\delta, \beta, k)$ are shown in the figure.
For the third order $a$PID: $\theta = 0.3,\ \eta = 10^4,\ \kappa_0 = 10,\ \alpha_0 = 1,\ n_0 = 1,\ k = 0.03,\ \gamma_0 = 0.5$ and $K_D \in [0, 5]$. $(\delta, \delta_0)$ are computed for each $K_D$ using (S31).
For the fourth order $a$PID: $\theta = 0.3,\ \eta = \eta_0 = 10^4,\ \theta_0 = 1,\ \bar{z}_3 = 0.01,\ k = 0.03,\ \alpha_1 = 05$ with $K_D \in [0, 2]$. $(\alpha_2, \mu_0)$ are then computed using (S36) and $\mu_0 = \theta_0(\alpha_1 \bar{z}_3 + \alpha_2 r)$ with $r = \mu/\theta = 10$. Only $(\alpha_1, \alpha_2)$ are shown in the figure.
- **Remark:** The temporal evolution of the variance and the stationary distribution are calculated by running $N = 50000$ stochastic trajectories, while the stationary variance is calculated by running a single simulation long enough ($t_{\text{Final}} = 2 \times 10^6$) and then taking time averages.

**Figure 10(b):**

- **Fixed Parameters:** $n_0 = 1,\ \kappa_1 = \kappa_0 = 10$ nM, $\gamma_1 = \gamma_2 = \delta_c = \gamma_0 = 0.028$ min$^{-1}$, $\eta = 0.05$ nM$^{-1}$ min$^{-1}$, $k_1 = \theta = 1$ min$^{-1}$, $k = 0.05$ min$^{-1}$ and $\alpha_0 = 100$ nM min$^{-1}$.
- **Variable Parameters:**
For I-control: $\delta = \delta_0 = 0$.
For PI-control: $\delta_0 = 0$ and $\delta = 5 \times 0.028$ min$^{-1}$.
For PID-control: $\delta_0 = 0.028$ min$^{-1}$ and $\delta = 5 \times 0.028$ min$^{-1}$.
For all controllers $\mu = 50$ nM min$^{-1}$ before $t = 8$ h, and it is doubled afterwards. Furthermore $\Delta = 0$ for $t < 16$ h and $\Delta = 3 \times 0.028$ min$^{-1}$ for $t \geq 16$ h.
- **Remark:** The parameters $\mu, \theta, \eta, k, k_1, \gamma_1, \gamma_2, \delta_c, \delta, \kappa_1$ and $\Delta$ are the same as those in Section S8 and are essentially extracted from [2]. Furthermore, $\gamma_0$ and $\delta_0$ are both chosen to be equal to the dilution rate since the rate of removal of proteins is dominated by dilution, and $\kappa_0$ is chosen to be the same as $\kappa_1$ to reflect a half-activation threshold equal to one fifth the set-point. Finally, since $\alpha_0$ represents the maximal gene expression rate of $\mathbf{Z_3}$, then it is comparable to $\mu$. Here we chose it to be double of $\mu$.

**Figure 11(b):** The chosen parameters for the cyberloop experiment are given in Table S2.

Table S2: Cyberloop experimental parameters

| Parameter | Value | Parameter | Value |
|---|---|---|---|
| $\mu$ | 14 min$^{-1}$ | $\delta$ | 0 (for I), 1 min$^{-1}$ (for PI and PID) |
| $\theta$ | 1 min$^{-1}$ | $\mu_0$ | 0 (for I and PI), 10 min$^{-1}$ (for PID) |
| $\eta$ | 5 min$^{-1}$ | $\theta_0$ | 0 (for I and PI), 0.1 min$^{-1}$ (for PID) |
| $k$ | 0.1 min$^{-1}$ | $\eta_0$ | 5 |
| $\alpha$ | 0.01 min$^{-1}$ | $\alpha_1$ | 0 (for I and PI), 1 min$^{-1}$ (for PID) |
| $\gamma$ | 0.1 min$^{-1}$ | $\alpha_2$ | 0 (for I and PI), 1 min$^{-1}$ (for PID) |
| $\kappa_1$ | 1 | | |

### S10 Useful Covariance Identities

Let $X, Y$ and $Z$ be two vector-valued random variables of possibly different dimensions. Let $X := \begin{bmatrix} X_1 \\ X_2 \end{bmatrix}$ and $Y := \begin{bmatrix} Y_1 \\ Y_2 \end{bmatrix}$, where $X_1, X_2, Y_1$ and $Y_2$ are all vector-valued random variables. Let $A$ and $B$ be two deterministic matrices with suitable dimensions. Furthermore, let $b$ be a deterministic vector. We have the following identities.

1. $\mathrm{Cov}\left[X, Y\right] := \mathbb{E}\left[\left(X - \mathbb{E}\left[X\right]\right)\left(Y - \mathbb{E}\left[Y\right]\right)^T\right] = \mathbb{E}\left[XY^T\right] - \mathbb{E}\left[X\right]\mathbb{E}\left[Y^T\right]$
2. $\mathrm{Cov}\left[X, Y\right] = \mathrm{Cov}\left[Y, X\right]^T$
3. $\mathrm{Cov}\left[b, X\right] = 0$
4. $\mathrm{Cov}\left[AX, BY\right] = A\mathrm{Cov}\left[X, Y\right]B^T$.
5. $\mathrm{Cov}\left[X, Y\right] = \begin{bmatrix}\mathrm{Cov}\left[X, Y_1\right] & \mathrm{Cov}\left[X, Y_2\right]\end{bmatrix} = \begin{bmatrix}\mathrm{Cov}\left[X_1, Y\right] \\ \mathrm{Cov}\left[X_2, Y\right]\end{bmatrix} = \begin{bmatrix}\mathrm{Cov}\left[X_1, Y_1\right] & \mathrm{Cov}\left[X_1, Y_2\right] \\ \mathrm{Cov}\left[X_2, Y_1\right] & \mathrm{Cov}\left[X_2, Y_2\right]\end{bmatrix}$
6. $\mathrm{Cov}\left[X_1 + X_2, Y_1 + Y_2\right] = \mathrm{Cov}\left[X_1, Y_1\right] + \mathrm{Cov}\left[X_1, Y_2\right] + \mathrm{Cov}\left[X_2, Y_1\right] + \mathrm{Cov}\left[X_2, Y_2\right]$
7. $\mathrm{Cov}\left[b^T X, X^T AX\right] = \sum_{i,j,k} b_k A_{ij}\mathrm{Cov}\left[X_k, X_i X_j\right]$
8. $\mathrm{Cov}\left[X^T AX, X^T BX\right] = \sum_{i,j,k,l} A_{ij} B_{k,l}\mathrm{Cov}\left[X_i X_j, X_k X_l\right]$

The proofs of 1 through 6 are straight forward. The proofs of 7 and 8 are given below.

*Proof of 7.*

$$\begin{aligned}
\mathrm{Cov}\left[b^T X, X^T AX\right] &= \mathbb{E}\left[b^T X X^T AX\right] - \mathbb{E}\left[b^T X\right]\mathbb{E}\left[X^T AX\right] \\
&= \mathbb{E}\left[\sum_k b_k X_k \sum_{i,j} A_{ij} X_i X_j\right] - \mathbb{E}\left[\sum_k b_k X_k\right]\mathbb{E}\left[\sum_{i,j} A_{ij} X_i X_j\right] \\
&= \sum_{i,j,k} b_k A_{ij}\Big(\mathbb{E}\left[X_k X_i X_j\right] - \mathbb{E}\left[X_k\right]\mathbb{E}\left[X_i X_j\right]\Big) \\
&= \sum_{i,j,k} b_k A_{ij}\mathrm{Cov}\left[X_k, X_i X_j\right]
\end{aligned}$$

□

*Proof of 8.*

$$\begin{aligned}
\mathrm{Cov}\left[X^T AX, X^T BX\right] &= \mathbb{E}\left[X^T AX X^T BX\right] - \mathbb{E}\left[X^T AX\right]\mathbb{E}\left[X^T BX\right] \\
&= \mathbb{E}\left[\sum_{i,j} A_{ij} X_i X_j \sum_{k,l} B_{kl} X_k X_l\right] - \mathbb{E}\left[\sum_{i,j} A_{ij} X_i X_j\right]\mathbb{E}\left[\sum_{k,l} B_{kl} X_k X_l\right] \\
&= \sum_{i,j,k,l} A_{ij} B_{kl}\Big(\mathbb{E}\left[X_i X_j X_k X_l\right] - \mathbb{E}\left[X_i X_j\right]\mathbb{E}\left[X_k X_l\right]\Big) \\
&= \sum_{i,j,k,l} A_{ij} B_{kl}\mathrm{Cov}\left[X_i X_j, X_k X_l\right]
\end{aligned}$$

□

#### S11 Useful Expectation and Covariance Approximations

Let $F, G : \mathbb{R}^n \to \mathbb{R}$ and $X \in \mathbb{R}^n$. We have the following approximations

1. $\mathbb{E}[F(X)] \approx F(\mathbb{E}[X]) + \frac{1}{2}\text{tr}\left\{\partial^2 F(\mathbb{E}[X])\text{Cov}[X]\right\}$
2. $\text{Cov}[F(X), G(X)] \approx \partial F(\mathbb{E}[X])\text{Cov}[X]\,\partial G(\mathbb{E}[X])^T$
3. If X follows a multivariate normal distribution, we have

$$\text{Cov}[F(X), G(X)] \approx \partial F(\mathbb{E}[X])\text{Cov}[X]\,\partial G(\mathbb{E}[X])^T + \frac{1}{2}\text{tr}\left\{\partial^2 F(\mathbb{E}[X])\text{Cov}[X]\,\partial^2 G(\mathbb{E}[X])\text{Cov}[X]\right\}.$$

Note that (1) and (3) are second order approximations while (2) is a first order approximation.

*Proof of 1.* A second order approximation of $F$ around the expected value of $X$, denoted here as $\bar{X}$ for convenience, can be written as

$$F(X) \approx F(\bar{X}) + \partial F(\bar{X})(X - \bar{X}) + \frac{1}{2}(X - \bar{X})^T \partial^2 F(\bar{X})(X - \bar{X}),$$

where $\partial F(\bar{X})$ (resp. $\partial^2 F(\bar{X})$) is a row vector (resp. square matrix) whose dimension is $n$ (resp. $n \times n$) that represents the directional derivative of $F$ (respectively Hessian), evaluated at $\bar{X}$. Taking the expectation of $F(X)$ yields

$$\begin{aligned}\mathbb{E}[F(X)] &\approx F(\bar{X}) + \frac{1}{2}\mathbb{E}\left[(X - \bar{X})^T \partial^2 F(\bar{X})(X - \bar{X})\right] \\ &\approx F(\bar{X}) + \frac{1}{2}\text{tr}\left\{\partial^2 F(\bar{X})\text{Cov}[X]\right\},\end{aligned}$$

where the first approximate equality follows from the fact that $\mathbb{E}\left[X - \bar{X}\right] = 0$, and the second approximate equality follows from the circular property of the trace operator. □

*Proof of 2.* Using a first order Taylor expansion for $F$ and $G$ around the expectation of $X$, denoted here by $\bar{X}$ for convenience, we proceed as follows

$$\begin{aligned}\text{Cov}[F(X), G(X)] &\approx \text{Cov}\left[F(\bar{X}) + \partial F(\bar{X})(X - \bar{X}), G(\bar{X}) + \partial G(\bar{X})(X - \bar{X})\right] \\ &\approx \text{Cov}\left[\partial F(\bar{X})(X - \bar{X}), \partial G(\bar{X})(X - \bar{X})\right] \\ &\approx \partial F(\bar{X})\text{Cov}\left[X - \bar{X}\right]\partial G(\bar{X})^T,\end{aligned}$$

which follows by exploiting identity 4 in Appendix S10. The proof is complete since $\text{Cov}\left[X - \bar{X}\right] = \text{Cov}[X]$. □

*Proof of 3.* Using a second order Taylor expansion for $F$ and $G$ around the expectation of $X$, denoted here by $\bar{X}$, we proceed as follows

$$\begin{aligned}&\text{Cov}[F(X), G(X)] \\ &\approx \text{Cov}\left[F(\bar{X}) + \partial F(\bar{X})(X - \bar{X}) + \frac{1}{2}(X - \bar{X})^T \partial^2 F(\bar{X})(X - \bar{X}), G(\bar{X}) + \partial G(\bar{X})(X - \bar{X}) + \frac{1}{2}(X - \bar{X})^T \partial^2 G(\bar{X})(X - \bar{X})\right] \\ &\approx \text{Cov}\left[\partial F(\bar{X})(X - \bar{X}) + \frac{1}{2}(X - \bar{X})^T \partial^2 F(\bar{X})(X - \bar{X}), \partial G(\bar{X})(X - \bar{X}) + \frac{1}{2}(X - \bar{X})^T \partial^2 G(\bar{X})(X - \bar{X})\right] \\ &\approx \partial F(\bar{X})\text{Cov}\left[X - \bar{X}\right]\partial G(\bar{X})^T + \frac{1}{2}\partial F(\bar{X})\text{Cov}\left[X - \bar{X}, (X - \bar{X})^T \partial^2 G(\bar{X})(X - \bar{X})\right] \\ &\qquad + \frac{1}{2}\text{Cov}\left[(X - \bar{X})^T \partial^2 F(\bar{X})(X - \bar{X}), X - \bar{X}\right]\partial G(\bar{X})^T \\ &\qquad + \frac{1}{4}\text{Cov}\left[(X - \bar{X})^T \partial^2 F(\bar{X})(X - \bar{X}), (X - \bar{X})^T \partial^2 G(\bar{X})(X - \bar{X})\right]\end{aligned}$$

Define the following deterministic vectors $a := \partial F(\bar{X})^T$ and $b := \partial G(\bar{X})^T$ and define the following symmetric matrices $A := \partial^2 F(\bar{X})$ and $B := \partial^2 G(\bar{X})$. Then we have

$$\begin{aligned}\text{Cov}[F(X), G(X)] \approx a^T \text{Cov}\left[X - \bar{X}\right] b &+ \frac{1}{2}\text{Cov}\left[a^T(X - \bar{X}), (X - \bar{X})^T B(X - \bar{X})\right] \\ &+ \frac{1}{2}\text{Cov}\left[b^T(X - \bar{X}), (X - \bar{X})^T A(X - \bar{X})\right] \\ &+ \frac{1}{4}\text{Cov}\left[(X - \bar{X})^T A(X - \bar{X}), (X - \bar{X})^T B(X - \bar{X})\right].\end{aligned}$$

Now, let's calculate each term separately. First we have that $\mathrm{Cov}\left[X-\bar{X}\right] = \mathrm{Cov}\left[X\right]$. The second term is calculated next using property 7.

$$\mathrm{Cov}\left[a^T(X-\bar{X}), (X-\bar{X})^T B (X-\bar{X})\right] = \sum_{i,j,k} a_k B_{i,j} \mathrm{Cov}\left[X_k - \bar{X}_k, (X_i-\bar{X}_i)(X_j-\bar{X}_j)\right] = 0,$$

because the odd central moments of a multivariate normal distribution are all zeros. The second term is also zero for the same reason. We are left with the last term which we calculate using property 8.

$$\begin{aligned}\mathrm{Cov}\left[(X-\bar{X})^T A (X-\bar{X}), (X-\bar{X})^T B (X-\bar{X})\right] &= \sum_{i,j,k,l} A_{ij} B_{kl} \mathbb{E}\left[(X_i-\bar{X}_i)(X_j-\bar{X}_j)(X_k-\bar{X}_k)(X_l-\bar{X}_l)\right] \\ &- \sum_{i,j,k,l} A_{ij} B_{kl} \mathbb{E}\left[(X_i-\bar{X}_i)(X_j-\bar{X}_j)\right] \mathbb{E}\left[(X_k-\bar{X}_k)(X_l-\bar{X}_l)\right]\end{aligned}$$

Using the fourth and second order moments of the multivariate normal distribution, we can write the right hand side as

$$\begin{aligned}&\sum_{i,j,k,l} A_{ij} B_{kl} \Big( \mathrm{Cov}\left[X_i, X_j\right] \mathrm{Cov}\left[X_k, X_l\right] + \mathrm{Cov}\left[X_i, X_l\right] \mathrm{Cov}\left[X_j, X_k\right] + \mathrm{Cov}\left[X_i, X_k\right] \mathrm{Cov}\left[X_l, X_j\right] - \mathrm{Cov}\left[X_i, X_j\right] \mathrm{Cov}\left[X_k, X_l\right] \Big) \\ &= \sum_{i,j,k,l} A_{ij} B_{kl} \Big( \mathrm{Cov}\left[X_i, X_l\right] \mathrm{Cov}\left[X_j, X_k\right] + \mathrm{Cov}\left[X_i, X_k\right] \mathrm{Cov}\left[X_l, X_j\right] \Big) \\ &= 2 \sum_{i,j,k,l} A_{ij} B_{kl} \mathrm{Cov}\left[X_i, X_l\right] \mathrm{Cov}\left[X_j, X_k\right],\end{aligned}$$

where the last equality follows because $A$ is symmetric. Now let's write the sum in matrix form by exploiting the symmetry of $A$ and $B$.

$$\begin{aligned}\sum_{i,j,k,l} A_{ij} B_{kl} \mathrm{Cov}\left[X_i, X_l\right] \mathrm{Cov}\left[X_j, X_k\right] &= \sum_{i,j,k,l} A_{ji} \mathrm{Cov}\left[X_i, X_l\right] B_{lk} \mathrm{Cov}\left[X_k, X_j\right] \\ &= \sum_{j,l} \left( \sum_i A_{ji} \mathrm{Cov}\left[X_i, X_l\right] \right) \left( \sum_k B_{lk} \mathrm{Cov}\left[X_k, X_j\right] \right) \\ &= \sum_{j,l} A_j \mathrm{Cov}\left[X, X_l\right] B_l \mathrm{Cov}\left[X, X_j\right] \\ &= \sum_j A_j \left( \sum_l \mathrm{Cov}\left[X, X_l\right] B_l \right) \mathrm{Cov}\left[X, X_j\right] \\ &= \sum_j A_j \mathrm{Cov}\left[X\right] B \mathrm{Cov}\left[X, X_j\right] \\ &= \sum_j e_j^T A \mathrm{Cov}\left[X\right] B \mathrm{Cov}\left[X\right] e_j = \mathrm{tr}\left\{ A \mathrm{Cov}\left[X\right] B \mathrm{Cov}\left[X\right] \right\}.\end{aligned}$$

Therefore, we have

$$\mathrm{Cov}\left[F(X), G(X)\right] \approx \partial F(\bar{X}) \mathrm{Cov}\left[X\right] \partial G(\bar{X})^T + \frac{1}{2} \mathrm{tr}\left\{ \partial^2 F(\bar{X}) \mathrm{Cov}\left[X\right] \partial^2 G(\bar{X}) \mathrm{Cov}\left[X\right] \right\}.$$

□

### References

[1] K. J. Åström and R. M. Murray, *Feedback systems: an introduction for scientists and engineers.* Princeton university press, 2010.

[2] S. K. Aoki, G. Lillacci, A. Gupta, A. Baumschlager, D. Schweingruber, and M. Khammash, "A universal biomolecular integral feedback controller for robust perfect adaptation," *Nature*, p. 1, 2019.

[3] T. Reeder and R. Schleif, "Arac protein can activate transcription from only one position and when pointed in only one direction," *Journal of Molecular Biology*, vol. 231, no. 2, pp. 205–218, 1993.

[4] C. Briat, A. Gupta, and M. Khammash, “Antithetic integral feedback ensures robust perfect adaptation in noisy biomolecular networks,” *Cell systems*, vol. 2, no. 1, pp. 15–26, 2016.

[5] Y. Qian and D. Del Vecchio, “Realizing ‘integral control’in living cells: how to overcome leaky integration due to dilution?,” *Journal of The Royal Society Interface*, vol. 15, no. 139, p. 20170902, 2018.

[6] T. Shopera, W. R. Henson, A. Ng, Y. J. Lee, K. Ng, and T. S. Moon, “Robust, tunable genetic memory from protein sequestration combined with positive feedback,” *Nucleic acids research*, vol. 43, no. 18, pp. 9086–9094, 2015.

[7] K. E. McGinness, T. A. Baker, and R. T. Sauer, “Engineering controllable protein degradation,” *Molecular Cell*, vol. 22, no. 5, pp. 701–707, 2006.
